# Distinct modes of dopamine modulation on striatopallidal synaptic transmission

**DOI:** 10.1101/2025.01.15.633147

**Authors:** Youngeun Lee, Maria Reva, Ki Jung Kim, Yemin Kim, Eunjeong Cho, Hyun-Jin Kim, Minseok Jeong, Kyungjae Myung, Yulong Li, Seung Eun Lee, Eunjoon Kim, C. Justin Lee, Christian Lüscher, Jae-Ick Kim

## Abstract

Dopamine (DA) affects voluntary movement by modulating basal ganglia function. In the classical model, DA depletion leads to overactivity of the indirect pathway and excessively inhibits the thalamus, resulting in hypokinesia. The contribution of DA on striatopallidal synapses, an initial hub in the indirect pathway connecting the striatum to the external globus pallidus (GPe), remains poorly understood because of the sparse DA innervation. Here, we combine optogenetic projection targeting, whole cell patch clamp recordings in acute brain slices from mice, and computational modeling to overcome this limitation. We show that DA activates D2R receptors (D2Rs) and D4 receptors (D4Rs) differentially in distinct GPe subregions. In a pinwheel-like fashion, dorsolateral and ventromedial GPe expresses high levels of D2Rs, which exert presynaptic inhibition, while in dorsomedial and ventrolateral GPe D4Rs cause postsynaptic inhibition. DA depletion by 6-OHDA (6-hydroxydopamine) reverses the region-specific effect of DA, shifting it in the opposite direction and contributing to hypokinesia. These findings reveal the mechanism by which the different modality information conveyed spatially through the indirect pathway is differentially modulated by DA at striatopallidal synapses.

## Introduction

The proper control of voluntary movement is vital for survival. The core brain regions modulating voluntary movement are the basal ganglia, a network of interconnected subcortical nuclei^1,2^. Sensory and motor information, widely distributed across the cortex, enters the basal ganglia, which contributes, among other functions, to select the appropriate motor strategy^3^. Central among these processes is the modulation of neuronal activity and synaptic transmission by the neuromodulator dopamine (DA) within the basal ganglia. The primary source of DA for goal-directed movement is the midbrain DA neurons located in the substantia nigra pars compacta (SNc), which predominantly innervate the striatum via the nigrostriatal pathway^4^. Within the striatum, the input nucleus of the basal ganglia, over 90% of neurons are GABAergic projection neurons known as medium spiny neurons (MSNs)^5^. Anatomically, the striatal MSNs can be categorized into two distinct groups: “direct” and “indirect” pathway MSNs. Interestingly, direct MSNs (dMSNs) projecting “directly” to the SNr/GPi (substantia nigra pars reticulata and internal globus pallidus, respectively), the primary output nuclei of the basal ganglia, express the DA D1 receptors (D1Rs), while indirect MSNs (iMSNs) projecting “indirectly” to the SNr/GPi by way of the GPe, express the DA D2 receptors (D2Rs)^1,2^. DA orchestrates the activity of MSNs through these receptors and the ensuing synaptic plasticity of excitatory afferents. In an action selection model, the direct pathway promotes appropriate motor strategies, whereas the indirect pathway suppresses other competing or inappropriate ones^3,4^. Consequently, the depletion of DA causes hyperactivity of the indirect pathway, resulting in an excessive inhibition of the thalamus and ultimately leading to the suppression of movement^5–7^.

Striatopallidal synapses in the GPe, the first synaptic connection between nuclei of the indirect pathway, are a critical node where the axon terminals of extensively distributed iMSNs converge. DA neurons in the midbrain are also known to project directly to the GPe via the nigropallidal pathway, which originates from the nigrostriatal pathway and terminates in the GPe^6,7^. However, studies to date have found only limited evidence supporting direct DA release from the nigropallidal pathway onto the GPe for the modulation of striatopallidal synaptic transmission^6,8–10^. On the other hand, it has been reported that the administration of DA or D2R agonists reduces striatopallidal GABAergic transmission with increased paired-pulse ratio (PPR), possibly attributable to the presence of presynaptic D2Rs at striatopallidal axon terminals^8–10^. In addition, GPe neurons seem to express other D2-like receptors, including D4Rs^11,12^, which may contribute to DA’s postsynaptic effect at striatopallidal synapses^11^. Despite these findings, our understanding of the dopaminergic modulation of striatopallidal transmission in the GPe is still incomplete. Notably, midbrain DA neurons are not a monolith; they encompass molecularly and functionally heterogeneous populations that can innervate anatomically and spatially distinct compartments of the striatum and other basal ganglia nuclei. This characteristic of DA neurons and their axons may explain in part the fact that each subdivision of the striatum is involved in specific animal behaviors^12,13^. In a similar vein, recent findings suggest that the indirect pathway can maintain anatomically parallel subnetworks, and striatopallidal synapses, depending on their anatomical location within the GPe, may convey different modalities of information^13,14^. However, the majority of previous studies on the GPe have treated striatopallidal synapses as uniform in their physiological properties. Hence, whether dopaminergic modulation of synaptic transmission can vary by the anatomical location of striatopallidal synapses in the GPe remains to be elucidated.

Here, we sought to determine how dopaminergic modulation shapes striatopallidal synaptic transmission in the GPe. Our data demonstrated that nigropallidal DA axons innervate the GPe with regional heterogeneity and their boutons release DA. In addition, DA D2-like receptors could modulate striatopallidal synaptic transmission in a region-specific manner across different subdivisions of the GPe. Specifically, the activation of D2-like receptors resulted in an increased paired-pulse ratio (PPR) and a decrease in GABAergic transmission within the dorsolateral (DL) and ventromedial (VM) regions of the GPe. Surprisingly, this dopaminergic effect on PPR was absent in the ventrolateral (VL) and dorsomedial (DM) GPe, despite the observed reduction in GABAergic transmission in these areas.

In contrast, DA depletion reversed the modulation of striatopallidal transmission by DA D2-like receptors in the GPe. 6-OHDA-induced DA depletion particularly promoted an increase in PPR in the VL and DM subregions of the GPe, while attenuating the rise in PPR in the DL and VM GPe. These results suggest that distinct sensory-motor information conveyed via the indirect pathway can be differentially modulated by DA, contingent upon the anatomical locations of striatopallidal synapses. Given that the structural and functional organization of basal ganglia circuits determines DA-related psychomotor functions and behaviors, our findings offer new insights into the previously overlooked role of dopaminergic modulation on striatopallidal synapses and globus pallidus in health and disease.

## Results

### DA axons capable of releasing DA innervate the GPe with regional heterogeneity

Although dopaminergic projection to the GPe via the nigropallidal pathway has been previously reported^6,7^, evidence of direct DA release onto the GPe from these DA terminals remains limited. We first examined the origin of nigropallidal DA axons in the GPe and found that the majority of TH-positive axon fibers originated from the SNc^15,16^ (Extended Data Fig. 1). Attempts to monitor DA transmission in the GPe using conventional electrochemical methods have been hindered by the sparse innervation of dopaminergic axon fibers to the GPe. However, recent advancements in fluorescent reporters for DA hold great promise for examining relatively weak DA transmission in brain regions such as the GPe^8,17,18^. To investigate whether dopaminergic axonal boutons innervating the GPe are functional and capable of releasing DA, we employed adeno-associated viruses (AAVs) encoding the genetically encoded fluorescent DA sensor GRAB_DA_ (rDA2h)^19^. These AAVs were injected into the GPe, and DA release was monitored by changes in fluorescence intensity within the imaging area. To compare the relative amplitude of DA transmission across other DA-projecting regions, AAVs expressing the GRAB_DA_ sensor were also injected into the M1 cortex and dorsolateral striatum (DLS). Three weeks post-injection, acute brain slices were prepared, and one-photon imaging combined with electrical stimulation was utilized to measure stimulus-evoked DA release. Although relatively small in amplitude compared to the DLS, electrical stimulation clearly evoked DA transmission in the GPe, indicating that DA axonal boutons in the GPe are functional and release DA (Fig. 1a,b).

**Fig. 1.**
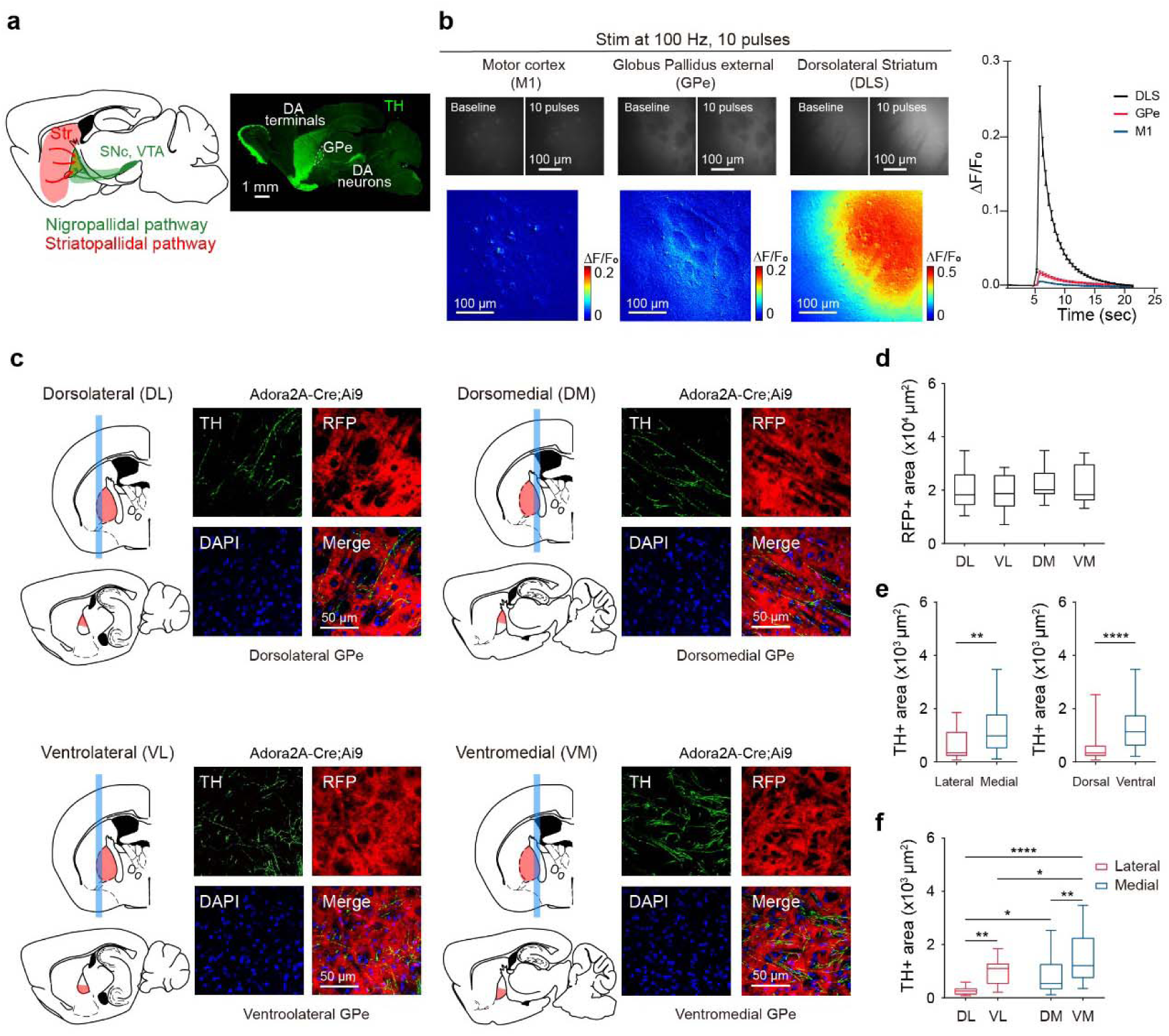
N**i**gropallidal **DA axons innervate the GPe with regional heterogeneity a**, A schematic illustration of the nigropallidal and striatopallidal pathways, and a fluorescent image of dopaminergic pathways in the brain. **b**, GRAB_DA_ (rDA2h) fluorescence responses in response to ten electrical pulses (left). A summary of the peak fluorescence changes in three brain regions in brain slices (right) (n = 15 slices from 4 mice for the GPe and M1 cortex, and n = 22 slices from 4 mice for the DLS). **c**, Schematic illustrations of the GPe subregions and representative confocal images showing TH+ nigropallidal and RFP+ striatopallidal axons in the GPe subregions of Adora2A-Cre;Ai9 mice. **d**, Quantification of RFP+ areas in the GPe subregions (n = 18 images from 3 mice per GPe subregion, one-way ANOVA; DL 19920 ± 1618 μm^2^, VL 18920 ± 1507 μm^2^, DM 22350 ± 1271 μm^2^, VM 21650 ± 1603 μm^2^; p = 0.3612). **e**, Summary statistics of TH+ area in the lateral and medial GPe subregions (left) and in the dorsal and ventral GPe subregions (right) (n = 36 images from 6 mice per each group, unpaired t-test; lateral 632.7 ± 84.3 μm^2^, medial 1190 ± 144.1 μm^2^, p = 0.0013; dorsal 570.8 ± 98.9 μm^2^, ventral 1252 ± 1216.1 μm^2^, p < 0.0001). **f,** Quantification of TH+ areas in the GPe subregions (n = 18 images from 3 mice per GPe subregion, ordinary two-way ANOVA with Holm-Sidak’s post-hoc multiple comparisons test; DL 280 ± 36.6 μm^2^, VL 985.4 ± 115.4 μm^2^, DM 861.7 ± 170.2 μm^2^, VM 1519 ± 209.3 μm^2^; DV axis effect, p < 0.0001, LM axis effect, p = 0.0003, interaction, p = 0.8713). The data are presented as box-and-whisker plots. *p < 0.05, **p < 0.01, ****p < 0.0001.

To further interrogate the innervation patterns of DA axons within the GPe subregions, we analyzed the regional variations of DA fibers by dividing the GPe into four subregions along the dorsoventral and lateromedial axes (Fig. 1c). We selectively expressed tdTomato in iMSNs by crossing Adora2A-Cre mice with Ai9 mice to visualize striatopallidal axon fibers. We quantified both striatopallidal and dopaminergic axonal areas in the GPe through double-immunostaining for tyrosine hydroxylase (TH) and tdTomato (RFP). The regional expression of tdTomato in Adora2A-Cre;Ai9 mice was consistent across the different subregions of the GPe (Fig. 1d). Interestingly, although TH-positive DA fibers are comparable within the striatal regions^20^, TH-positive DA axons innervated the GPe with significant spatial heterogeneity; DA fibers are more abundant in the medial and ventral parts of the GPe compared to the lateral and dorsal regions (Fig. 1e,f).

### DA D2-like receptors modulate striatopallidal transmission through distinct mechanisms within the subregions of the GPe

The bath application of either DA or D2-like receptors agonist quinpirole in the GPe has been documented to reduce GABAergic transmission at striatopallidal synapses while simultaneously increasing the paired pulse ratio (PPR)^10,21,22^. These findings indicate a presynaptic action of DA through D2Rs at striatopallidal axon terminals originating from iMSNs. We first revisited the functional role of striatopallidal presynaptic D2Rs on GABAergic synaptic transmission by measuring the PPR to indirectly monitor release probability^23,24^. Most of the previous studies examining striatopallidal synapses have utilized electrical stimulation to evoke striatopallidal synaptic transmission. However, this approach is inherently prone to the unintended activation of other axons including collaterals from dMSNs axons, pallidostriatal axons, and GPe collateral axons^25–27^. To avoid this possibility, we employed optogenetic techniques to selectively stimulate striatopallidal axons^25,26^. Specifically, we expressed channelrhodopsin-2 (ChR2) in iMSNs by crossing Adora2A-Cre mice with transgenic mice harboring a conditional floxed allele of ChR2 in the Rosa26 locus (Ai32 mice) (Fig. 2a). First, we validated the stable maintenance of the oIPSC (optically evoked inhibitory postsynaptic current) baseline over a 40-minute period using ChR2-mediated optogenetic stimulation (Extended Data Fig. 2a,b). Consistent with previous reports using electrical stimulation^21,22^, the application of quinpirole significantly reduced the light-evoked GABAergic transmission (oIPSCs) and increased the PPR at striatopallidal synapses (Fig. 2b,c and Extended Data Fig. 2c,d). Although differences in the oIPSC kinetics of striatopallidal synaptic transmission were observed depending on the postsynaptic GPe neuronal types (Table S1), the response to DA D2-like receptor activation did not differ between GPe neuronal types, consistent with previous studies^21^.

**Fig. 2.**
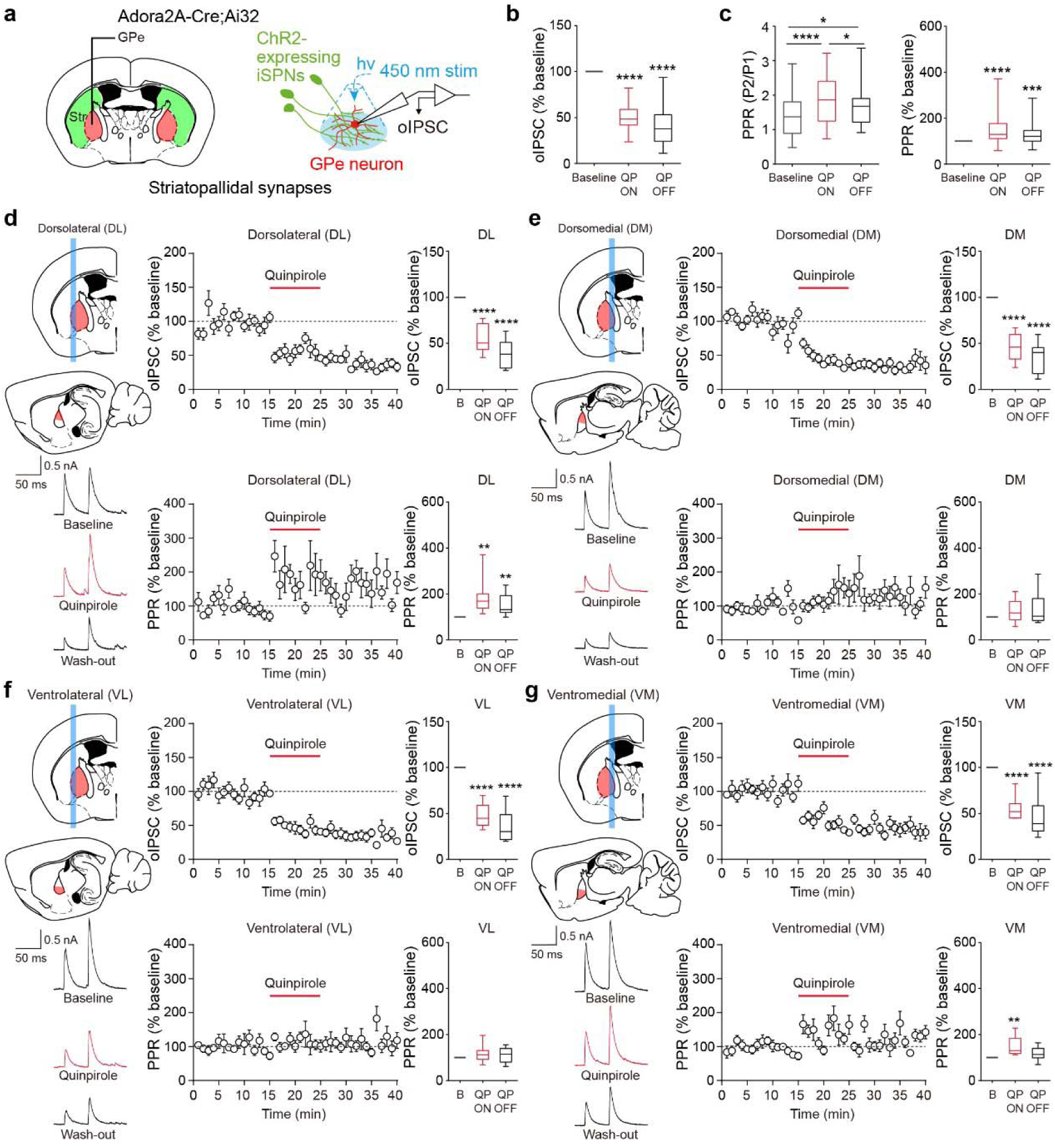
DA D2-like receptors modulate striatopallidal transmission through distinct mechanisms within the subregions of the GPe **a**, A schematic illustration depicting electrophysiological recording of striatopallidal synaptic transmission via optogenetic stimulation in Adora2A-Cre;Ai32 mice. **b,** Summary statistics of optogenetically evoked oIPSCs in GPe neurons during bath application of quinpirole (10 μM) (n = 40 cells from 26 mice, one sample t-test; Baseline 100%, QP ON 50.60 ± 2.17%, QP OFF 38.99 ± 2.77%). **c,** Summary statistics of the absolute values (left) and normalized values (right) of optogenetically evoked PPRs in striatopallidal synaptic transmission during bath application of quinpirole (10 μM) (n = 40 cells from 26 mice; left, repeated measures one-way ANOVA with Holm-Sidak’s post-hoc multiple comparisons test, Baseline 1.44 ± 0.10, QP ON 1.89 ± 0.11, QP OFF 1.69 ± 0.09, p < 0.0001; right, one sample t-test, Baseline 100%, QP ON 143.10 ± 8.88%, QP OFF 129.80 ± 7.73%). **d-g,** Schematic illustrations of the GPe subregions (top left), representative recording traces (bottom left), oIPSC amplitude (top center) and PPR plots (bottom center) normalized to baseline during bath application of quinpirole (10 μM), and summary statistics of normalized oIPSCs (top right) and normalized PPRs (bottom right). **d,** Summary statistics of normalized oIPSCs (n = 10 cells from 9 mice, one sample t-test; Baseline 100%, QP ON 54.38 ± 4.60%, QP OFF 39.33 ± 4.79%) and normalized PPRs (Baseline 100%, QP ON 184 ± 23.37%, QP OFF 154.10 ± 14.71%) in the DL GPe. **e,** Summary statistics of normalized oIPSCs (n = 10 cells from 8 mice, one sample t-test; Baseline 100%, QP ON 45.29 ± 4.53%, QP OFF 34.79 ± 5.23%) and normalized PPRs (Baseline 100%, QP ON 126.30 ± 15.15%, QP OFF 135.30 ± 22.82%) in the DM GPe. **f,** Summary statistics of normalized oIPSCs (n = 10 cells from 8 mice, one sample t-test; Baseline 100%, QP ON 47.49 ± 4.07%, QP OFF 35.84 ± 5.35%) and normalized PPRs (Baseline 100%, QP ON 115.90 ± 11.32%, QP OFF 112.20 ± 10.24%) in the VL GPe. **g,** Summary statistics of normalized oIPSCs (n = 10 cells from 8 mice, one sample t-test; Baseline 100%, QP ON 55.24 ± 3.85%, QP OFF 45.99 ± 6.78%) and normalized PPRs (Baseline 100%, QP ON 146.30 ± 12.77%, QP OFF 117.50 ± 8.48%) in the VM GPe. The data are presented as box-and-whisker plots. *p < 0.05, **p < 0.01, ***p < 0.001, ****p < 0.0001.

As mentioned above, most prior studies have assumed that striatopallidal synapses in the GPe possess homogeneous physiological characteristics. Given the regional heterogeneity in dopaminergic innervation within the GPe that we observed, we next investigated whether DA D2-like receptors differentially modulate striatopallidal synaptic transmission across different anatomical locations in the GPe. Based on the distinct innervation patterns of nigropallidal DA axons within the GPe (Fig. 1c-f), we assessed the effect of quinpirole on GABAergic transmission and the PPR at striatopallidal synapses by dividing the GPe into four subregions along the dorsoventral and lateromedial axes (Fig. 2d-g). In addition, we deliberately aimed to record neurons at a distance from the overlapping areas within the four GPe subregions to more clearly capture the potential physiological heterogeneity in striatopallidal transmission. Most importantly, we observed that the effect of quinpirole at striatopallidal synapses exhibited significant variation dependent on the specific subregions of the GPe. While quinpirole universally suppressed GABAergic transmission across all subregions of the GPe, it induced distinct alterations in the PPR. Quinpirole treatment increased the PPR in the dorsolateral (DL) and ventromedial (VM) subregions, whereas no significant change in PPR was observed in the ventrolateral (VL) and dorsomedial (DM) subregions of the GPe (Fig. 2d-g and Extended Data Fig. 2e-h). Coe cient of variation analysis also showed that VL and DM GPe were plotted closer to the horizontal axis compared to DL and VM GPe, suggesting that the effects of D2-like receptors activation on striatopallidal synaptic transmission may involve a complex interplay of both presynaptic and postsynaptic mechanisms (Extended Data Fig. 2i).

In light of the regional heterogeneity observed in dopaminergic modulation within the GPe, we hypothesized the presence of an underlying anatomical factor that might partially account for this variation. To explore this, we quantified the nearest neighbor distances between putatively functional DA boutons and striatopallidal synapses. Utilizing Adora2A-Cre;Ai9 mice, we performed multi-color immunostaining coupled with enhanced confocal microscopy. The co- localization of the presynaptic protein Bassoon and TH was considered as functional DA boutons^28,29^. Furthermore, striatopallidal GABAergic synapses were identified by the co- localization of presynaptic RFP and postsynaptic GABA_A_R (Extended Data Fig. 3a,b). Interestingly, the DL and VM regions of the GPe, which exhibited elevated PPR in response to quinpirole, displayed shorter nearest neighbor distances between functional DA boutons and striatopallidal GABAergic synapses compared to the VL and DM GPe (Extended Data Fig. 3c-e). This anatomical characteristics within the GPe may also have functional implications for the region-dependent variability in dopaminergic modulation of striatopallidal transmission.

### Region-specific dopaminergic modulation of striatopallidal synaptic transmission in the GPe is driven by differential contributions of presynaptic D2Rs and postsynaptic D4Rs

To determine how the activation of D2-like receptors differently modulates striatopallidal transmission across various subregions of the GPe, we compared the quantal properties of striatopallidal GABAergic transmission before and after treatment with quinpirole. For quantal analysis, Ca^2+^ was substituted with Sr^2+^ in the bath solution to facilitate asynchronous release. Striatopallidal GABAergic transmission in some GPe neurons did not respond to Sr^2+^ treatment; consequently, these cells were excluded from the quantal analysis. It is noteworthy that neurons in the lateral GPe regions (DL, VL) exhibited a lower propensity for evoking asynchronous release. When Sr^2+^ was applied, no significant differences were observed in the quantal amplitude and frequency of oIPSCs among the GPe subregions. Following quinpirole treatment with Sr^2+^ replacement, a reduction in quantal amplitude was observed across all GPe subregions (Fig. 3a-d). However, a decrease in quantal frequency was specifically found in the DL and VM GPe subregions (Fig. 3a,d,e), where an elevation in PPR by quinpirole was also detected. These findings suggest that the activation of postsynaptic D2-like receptors in the GPe universally suppresses GABAergic transmission across all subregions, while presynaptic D2-like receptors in the DL and VM GPe modulate GABA release.

**Fig. 3.**
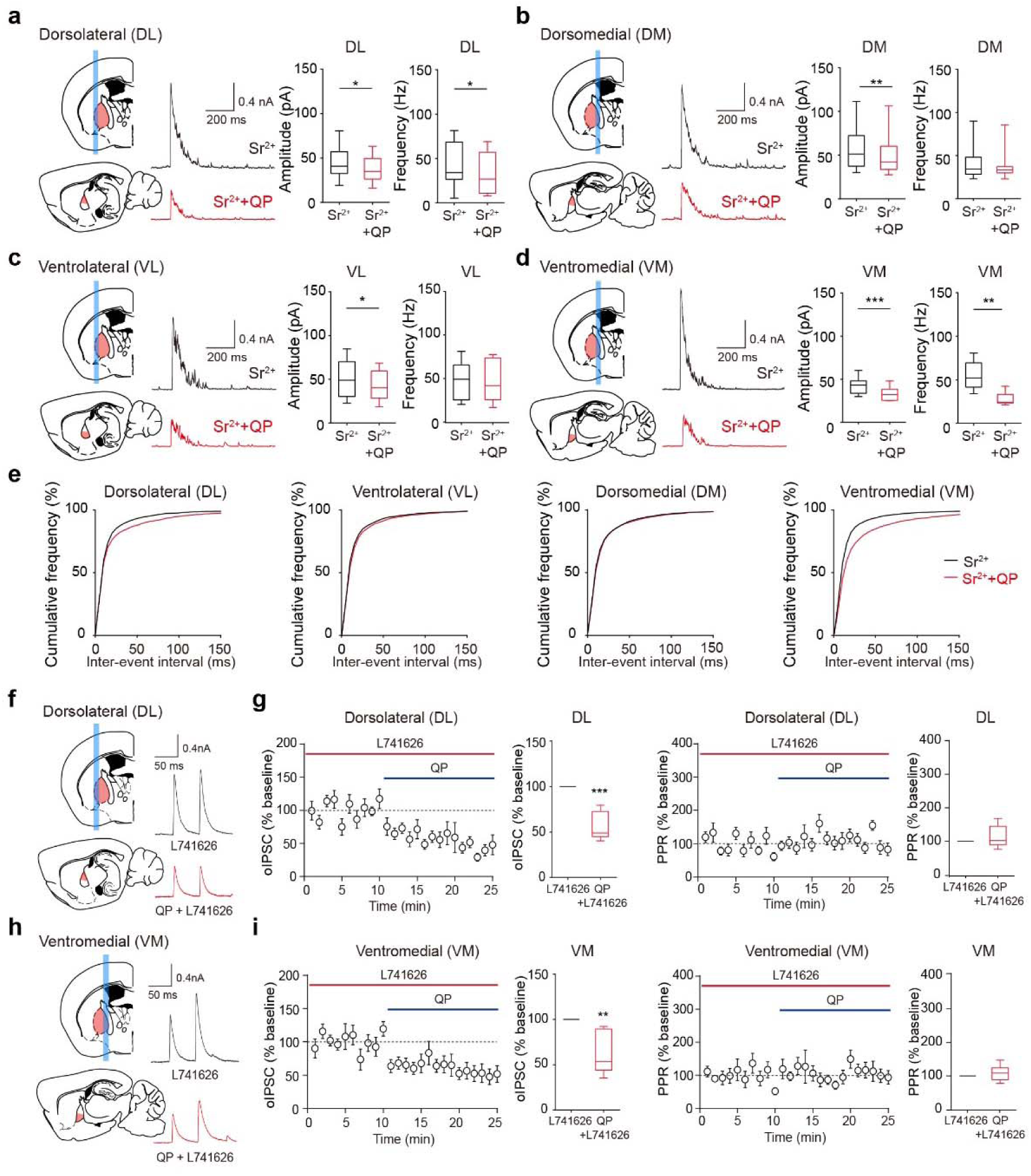
R**e**gion**-specific dopaminergic modulation of striatopallidal synaptic transmission in the GPe is driven by differential contributions of presynaptic D2Rs and postsynaptic D4Rs a-d**, Schematic illustrations of the GPe subregions and representative recording traces of optogenetically evoked asynchronous GABA release (quantal transmission) in Adora2A- Cre;Ai32 mice during bath application of strontium (4 mM) and quinpirole (10 μM) (left). Summary statistics of quantal oIPSC amplitude and frequency (right). **a**, Summary statistics of quantal oIPSC amplitude (n = 8 cells from 6 mice, paired t-test; Sr^2+^ 44.48 ± 6.68 pA, Sr^2+^ + QP 37.54 ± 5.32 pA) and frequency (Sr^2+^ 42.80 ± 9.12 Hz, Sr^2+^ + QP 31.80 ± 8.30 Hz) in the DL GPe. **b**, Summary statistics of quantal oIPSC amplitude (n = 16 cells from 15 mice, paired t-test; Sr^2+^ 52.31 ± 6.07 pA, Sr^2+^ + QP 47.91 ± 6.03 pA) and frequency (Sr^2+^ 41.03 ± 4.74 Hz, Sr^2+^ + QP 40.26 ± 4.35 Hz) in the DM GPe. **c**, Summary statistics of quantal oIPSC amplitude (n = 7 cells from 7 mice, paired t-test; Sr^2+^ 52.84 ± 8.51 pA, Sr^2+^ + QP 45.05 ± 6.83 pA) and frequency (Sr^2+^ 46.85 ± 7.95 Hz, Sr^2+^ + QP 43.29 ± 8.96 Hz) in the VL GPe. **d**, Summary statistics of quantal oIPSC amplitude (n = 9 cells from 8 mice, paired t-test; Sr^2+^ 42.62 ± 3.15 pA, Sr^2+^ + QP 32.38 ± 2.64 pA) and frequency (Sr^2+^ 54.32 ± 5.36 Hz, Sr^2+^ + QP 28.63 ± 2.38 Hz) in the VM GPe. **e**, Cumulative plots of inter-event intervals for quantal oIPSCs at striatopallidal synapses. **f,h,** Schematic illustrations and representative recording traces of optogenetically evoked oIPSCs in the DL and VM GPe during bath application of L-741626 (100 nM) and quinpirole (100 nM). **g,** Normalized oIPSC amplitude plot and summary statistics in the DL GPe (n = 7 cells from 5 mice, one sample t-test; L-741626 100%, L-741626 + QP 56.04 ± 5.73%) (left). Normalized PPR plot and summary statistics in the DL GPe (L-741626 100%, L-741626 + QP 111.70 ± 12.13%) (right). **i,** Normalized oIPSC amplitude plot and summary statistics in the VM GPe (n = 7 cells from 4 mice, one sample t-test; L-741626 100%, L-741626 + QP 61.37 ± 8.55%) (left). Normalized PPR plot and summary statistics in the VM GPe (L-741626 100%, L-741626 + QP 106.8 ± 8.90%). The data are presented as box-and-whisker plots. *p < 0.05, **p < 0.01, ***p < 0.001.

To explore potential regional variations in the quantal kinetics of striatopallidal synaptic transmission in the GPe, we conducted an AI-based clustering analysis on Sr^2+^-induced quantal responses. This analysis aimed to determine whether specific quantal characteristics varied across the DL, VL, DM, and VM regions of the GPe. By applying principal component analysis (PCA) for dimensionality reduction and K-means clustering, we classified the quantal responses into up to 50 distinct categories based on their amplitude, rise time, and decay time (Extended Data Fig. 4a). Comparison of these categories across the GPe subregions revealed a notable divergence in the VM region. Unlike the DL, VL, and DM regions, which shared similar quantal characteristics, the VM region exhibited unique quantal categories with significantly different kinetics (Extended Data Fig. 4b-d). This distinct pattern may indicate that striatopallidal synapses in the VM region of the GPe may originate from functionally different sources in the striatum or employ a unique set of presynaptic mechanisms, potentially reflecting functional specialization within the GPe circuitry. As a further investigation into the synaptic vesicle release machinery, we examined the individual isoforms of synaptotagmin at striatopallidal axon terminals to determine potential subregional variations in the presynaptic fusion machinery within the GPe (Extended Data Fig. 5a,d,g). Our findings showed that striatopallidal terminals are predominantly co-localized with synaptotagmin 1 across all GPe subregions (Extended Data Fig. 5j,k). Synaptotagmin 7 exhibited slightly higher expression in the medial and dorsal regions relative to the lateral and ventral regions of the GPe (Extended Data Fig. 5e,f), while synaptotagmin 5/9 was preferentially localized in the dorsal GPe compared to the ventral GPe (Extended Data Fig. 5h,i). These observations implicate subregion-specific heterogeneity of striatopallidal synapses in basal synaptic features including quantal characteristics and synaptic release machinery.

We next focused on identifying D2-like receptors at presynaptic and postsynaptic sites of striatopallidal synapses across the different GPe subregions. GPe neurons in rodents are reported to express DA D4 receptors (D4Rs)^11,12^, yet there are conflicting studies regarding the postsynaptic dopaminergic effects on striatopallidal synaptic transmission^21,22,30^. We pharmacologically evaluated the postsynaptic effect of D4Rs on striatopallidal transmission using optogenetic stimulation combined with a highly selective D4R agonist A-412997 (2- (3′,4′,5′,6′-tetrahydro-2′H-[2,4′] bipyridinyl-1′-yl)-N-m-tolyl-acetamide). If GABAergic transmission at striatopallidal synapses is suppressed by postsynaptic D4Rs, treatment with a selective D4R agonist A-412997 would result in an attenuation of oIPSCs without altering the PPR. Consistent with our quantal analysis of striatopallidal transmission, treatment with A- 412997 led to a widespread reduction in oIPSCs across all GPe subregions, without affecting the PPR (Extended Data Fig. 6). This finding suggests that postsynaptic D4Rs inhibit GABAergic transmission uniformly throughout the GPe. To clearly validate the region-specific role of presynaptic D2Rs in regulating GABA release at striatopallidal synapses, we assessed the effects of quinpirole on oIPSCs and PPR under continuous bath application of the D2R-selective antagonist L-741626. We found that quinpirole continued to decrease GABAergic transmission in the presence of L-741626 application; however, the previously observed increase in PPR induced by quinpirole was no longer present in the DL and VM GPe regions (Fig. 3f-i).

We next questioned whether differences in the localization and spatial distribution of D2Rs at striatopallidal axon terminals might contribute, at least in part, to the region-specific dopaminergic modulation of synaptic transmission in the GPe. To selectively label striatopallidal axons and synaptic terminals, we injected a Cre-inducible adeno-associated virus (AAV) expressing both mGFP and synaptophysin-mRuby into the entire striatum of Adora2A-Cre mice (Fig. 4a,b). Subsequently, we performed multi-color immunostaining and enhanced confocal imaging to quantify the spatial localization of D2Rs at striatopallidal axon terminals (RFP+) or axons (GFP+) within each GPe subregion (Fig. 4c). Our results revealed that the DL and VM GPe subregions exhibited a higher degree of co-localization of D2Rs on striatopallidal axons (D2R+ GFP+) and axon terminals (D2R+ RFP+) compared to the VL and DM GPe subregions (Fig. 4d). Additionally, the nearest neighbor distance between D2Rs on striatopallidal axons or axon terminals was shorter in the DL and VM GPe compared to the VL and DM GPe (Fig. 4e). We also investigated the expression of D4Rs and GABA_A_Rs on postsynaptic neurons in the GPe. In Adora2A-Cre;Ai9 mice, striatopallidal axons were labeled with RFP and GABA_A_Rs were immunostained to mark the dendrites of GPe neurons (Fig. 4f,g). Using multi-color immunostaining and 3D reconstruction, we were able to clearly identify the separation between striatopallidal axons (RFP+) and postsynaptic neuronal dendrites labeled with GABA_A_R in the GPe (Fig. 4h). We found that both the volume and number of D4R-positive signals were significantly greater in the VL and DM GPe subregions compared to the DL and VM GPe, whereas no significant differences were observed in the volume and number of GABA_A_R- positive signals (Fig. 4i,j). Furthermore, the nearest neighbor distances between D4Rs and GABA_A_Rs were markedly shorter in the VL and DM GPe compared to the DL and VM GPe (Fig. 4k). These findings suggest that region-specific differences in the expression, localization, and spatial distribution of presynaptic D2Rs and postsynaptic D4Rs may partially underly the distinct dopaminergic modulation of striatopallidal transmission observed across GPe subregions.

**Fig. 4.**
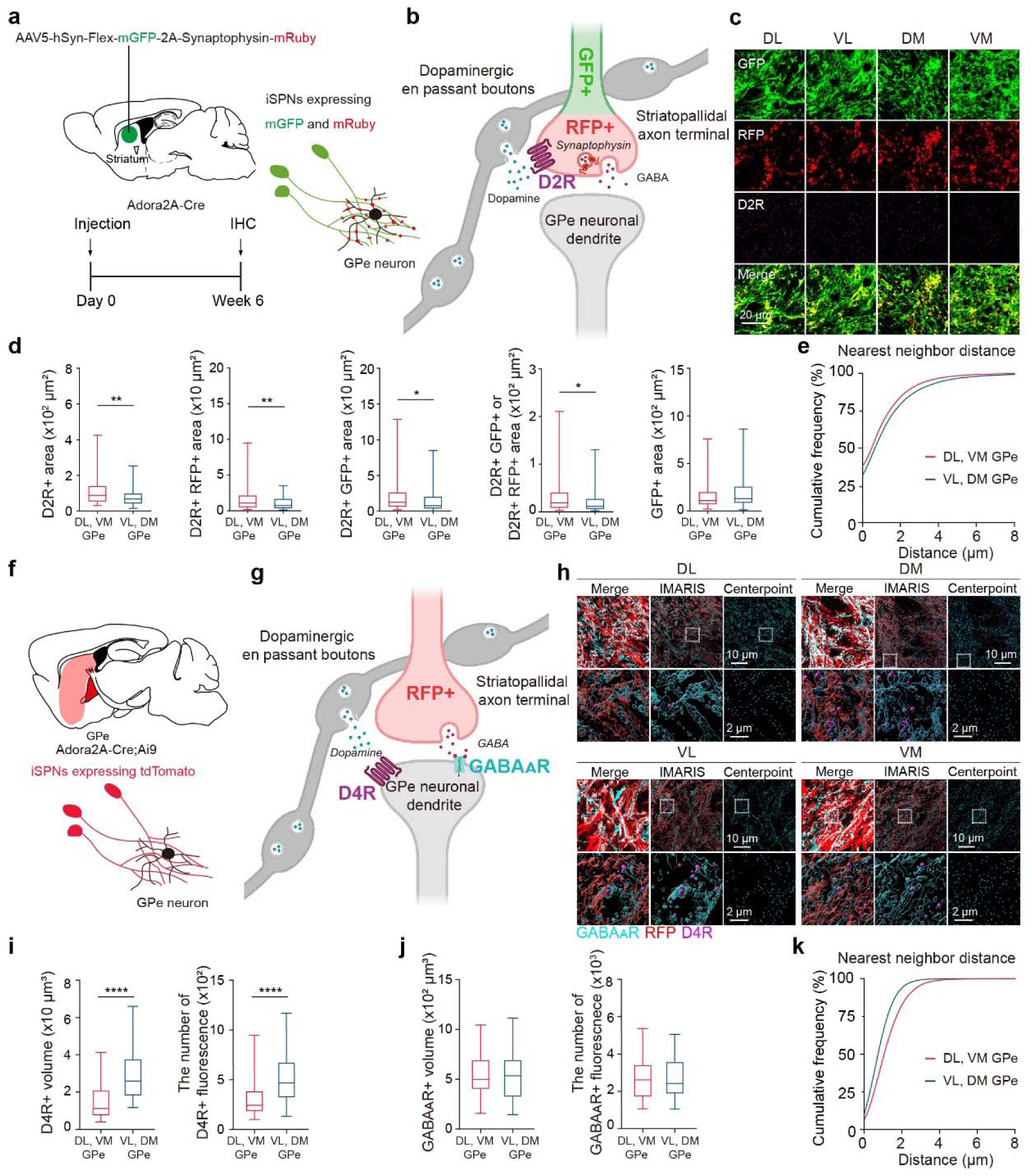
The distinct spatial localization and distribution of presynaptic D2Rs and postsynaptic D4Rs in the GPe subregions may contribute to the region-specific dopaminergic modulation of striatopallidal transmission **a**, A schematic illustration describing the injection of AAV5-hSyn-Flex-mGFP-2A- Synaptophysin-mRuby virus into the striatum of Adora2A-Cre mice to selectively label striatopallidal axons and terminals. **b**, A schematic illustration depicting fluorescently labeled striatopallidal axons (GFP), axon terminals (RFP), and D2Rs on iMSNs. **c**, Representative enhanced confocal images of striatopallidal axons, axon terminals, and D2Rs in the GPe subregions. **d,** Summary statistics of D2R+ area (n = 95 images from 9 mice per GPe subgroup, unpaired t-test; DL-VM GPe 109.70 ± 7.91 μm^2^, VL-DM GPe 80.39 ± 4.99 μm^2^), D2R+ area colocalized with RFP+ area (n = 93 images from 9 mice per GPe subgroup; DL-VM GPe 15.61 ± 1.52 μm^2^, VL-DM GPe 10.80 ± 0.90 μm^2^), D2R+ area colocalized with GFP+ area (n = 76 images from 9 mice per GPe subgroup; DL-VM GPe 21.96 ± 2.80 μm^2^, VL-DM GPe 14.51 ± 1.66 μm^2^), D2R+ area colocalized with GFP+ or RFP+ area (n = 92 images from 9 mice per GPe subgroup; DL-VM GPe 29.78 ± 3.41 μm^2^, VL-DM GPe 21.41 ± 2.36 μm^2^), and GFP+ area (n = 96 images from 9 mice per GPe subgroup, DL-VM GPe 163.60 ± 14.08 μm^2^, VL-DM GPe 187.40 ± 14.62 μm^2^). **e,** Cumulative plots for the nearest neighbor distance from striatopallidal axon terminals (RFP+) to D2Rs on striatopallidal axons or axon terminals (D2R+ RFP+ or D2R+ GFP+). **f**, A schematic illustration depicting fluorescently labeled striatopallidal axons (RFP) in the GPe, projected from iMSNs of Adora2A-Cre;Ai9 mice. **g**, A schematic illustration describing fluorescently labeled striatopallidal axons (RFP), GABA_A_Rs, and D4Rs in the GPe of Adora2A- Cre;Ai9 mice. **h,** Representative enhanced confocal images, restructured 3D images by IMARIS, and centerpoints of GABA_A_Rs and D4Rs in the GPe subregions. **i,** Summary statistics of D4R+ fluorescence volume (n = 47 images from 6 mice per GPe subgroup, unpaired t-test; DL-VM GPe 14.58 ± 1.32 μm^3^, VL-DM GPe 29.06 ± 1.84 μm^3^) and number (DL-VM GPe 317.80 ± 30.08, VL-DM GPe 534.10 ± 32.20). **j,** Summary statistics of GABA_A_R+ fluorescence volume (n = 47 images from 6 mice per GPe subgroup, unpaired t-test; DL-VM GPe 533.50 ± 29.48 μm^3^, VL-DM GPe 536.60 ± 31.58 μm^3^) and number (DL-VM GPe 2686.30 ± 169.60, VL-DM GPe 2685.80 ± 143.30). **k,** Cumulative plots for the nearest neighbor distance from postsynaptic GABA_A_Rs to postsynaptic D4Rs in the GPe. The data are presented as box-and-whisker plots. *p < 0.05, **p < 0.01, ****p < 0.0001.

### Pre- and postsynaptic DA receptors fine-tune ongoing activity in the GPe through region-specific modulations

Information spanning a wide range of frequencies, from low to high, propagates through the basal ganglia circuits including the indirect pathway. To further explore the functional implications of region-specific dopaminergic modulations on striatopallidal transmission, beyond their effects on PPR, we investigated how DA receptor activation influences ongoing activity at striatopallidal synapses. We optogenetically stimulated striatopallidal axon terminals with trains of 10 light pulses at 20 Hz, and compared the inhibitory postsynaptic responses across the GPe subregions^31^. We found that postsynaptic responses to 10-pulse stimulation varied significantly depending on the anatomical location within the GPe (Fig. 5a,d,g,j). High-frequency stimulation often leads to short-term depression at synapses, primarily due to the depletion of the readily releasable pool (RRP)^32^. In the GPe, such stimulation induced pronounced short-term depression in the DL and VL subregions, with GABAergic transmission significantly suppressed throughout the stimulation trains (Fig. 5a-f). In contrast, in the VM GPe, oIPSCs were initially facilitated but ultimately transitioned to synaptic depression (Fig. 5j-l). Notably, striatopallidal synapses in the DM GPe consistently exhibited short-term facilitation, underscoring the heterogeneity of synaptic modulation across GPe subregions (Fig. 5g-i,m).

**Fig. 5.**
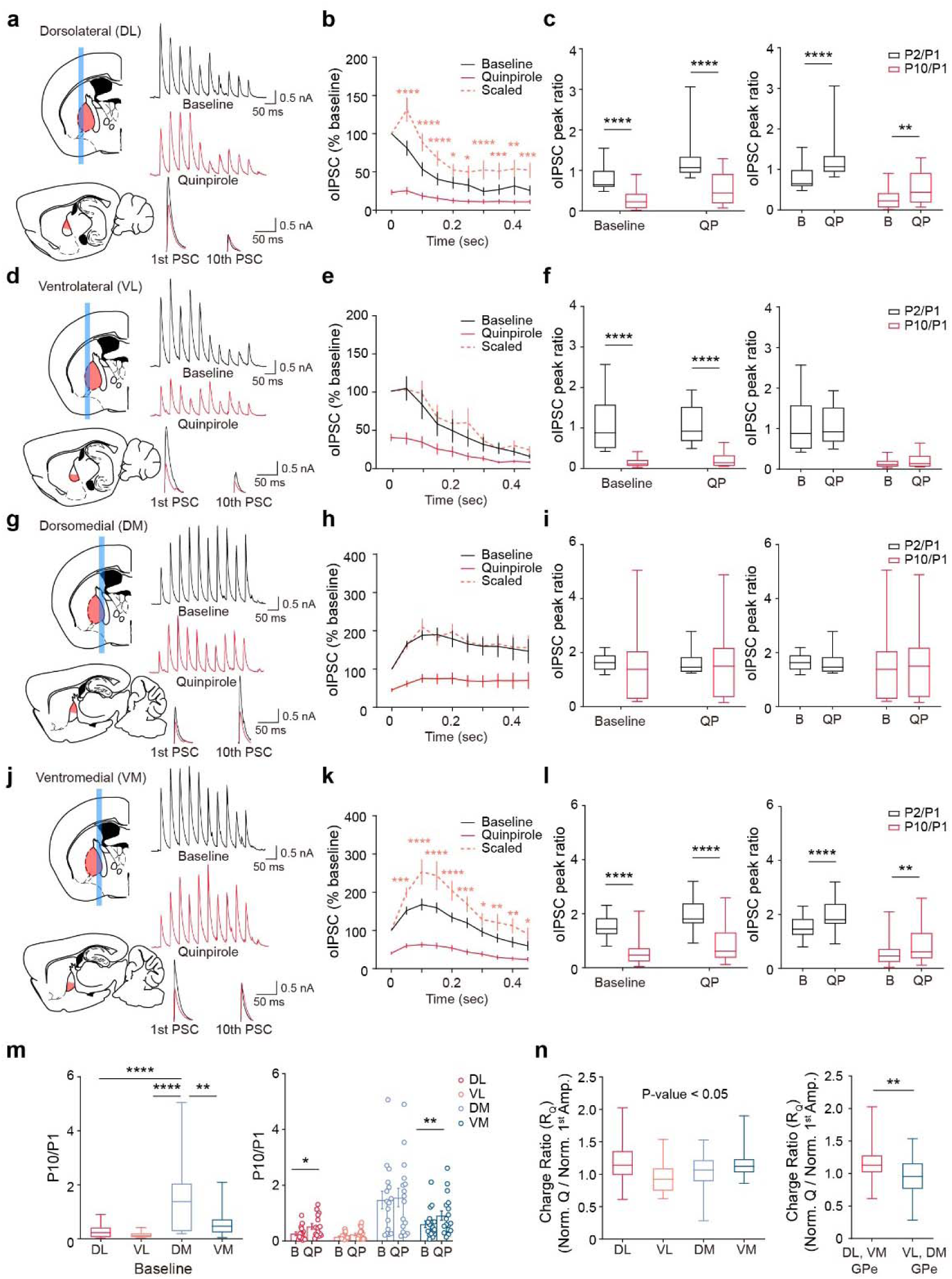
Pre- and postsynaptic DA receptors fine-tune ongoing activity in the GPe through region-specific modulations **a,d,g,j,** Schematic illustrations of the GPe subregions (left) and representative recording traces of optogenetically evoked oIPSCs (right), generated by ten light pulses at 20 Hz before and after bath application of quinpirole (10 μM). **b,e,h,k,** Summary plots of ongoing activity in the GPe subregions. **c**, Summary statistics of oIPSC amplitude peak ratios for comparative analysis between P2/P1 and P10/P1 in the DL GPe (n = 15 cells from 13 mice, repeated measures two- way ANOVA with Holm-Sidak’s post-hoc multiple comparisons test; Baseline P2/P1 0.795 ± 0.089, Baseline P10/P1 0.258 ± 0.065, QP P2/P1 1.259 ± 0.149, QP P10/P1 0.520 ± 0.101; QP treatment, p = 0.0003, peak ratio, p < 0.0001, interaction, p = 0.0648). **f**, Summary statistics of oIPSC amplitude peak ratios for comparative analysis between P2/P1 and P10/P1 in the VL GPe (n = 15 cells from 13 mice, repeated measures two-way ANOVA with Holm-Sidak’s post-hoc multiple comparisons test; Baseline P2/P1 1.037 ± 0.159, Baseline P10/P1 0.149 ± 0.028, QP P2/P1 1.023 ± 0.113, QP P10/P1 0.232 ± 0.051; QP treatment, p = 0.5132, peak ratio, p < 0.0001, interaction, P = 0.2939). **i**, Summary statistics of oIPSC amplitude peak ratios for comparative analysis between P2/P1 and P10/P1 in the DM GPe (n = 16 cells from 14 mice, repeated measures two-way ANOVA with Holm-Sidak’s post-hoc multiple comparisons test; Baseline P2/P1 1.654 ± 0.075, Baseline P10/P1 1.467 ± 0.317, QP P2/P1 1.618 ± 0.108, QP P10/P1 1.555 ± 0.335; QP treatment, p = 0.8014, peak ratio, p = 0.6871, interaction, p = 0.3852). **l**, Summary statistics of oIPSC amplitude peak ratios for comparative analysis between P2/P1 and P10/P1 in the VM GPe (n = 17 cells from 15 mice, repeated measures two-way ANOVA with Holm-Sidak’s post-hoc multiple comparisons test; Baseline P2/P1 1.520 ± 0.103, Baseline P10/P1 0.601 ± 0.125, QP P2/P1 1.991 ± 0.135, QP P10/P1 0.909 ± 0.170; QP treatment, p < 0.0001, peak ratio, p < 0.0001, interaction, p = 0.1611). **m**, Summary statistics of P10/P1 ratios in the GPe subregions under basal state (left) (one-way ANOVA with Holm-Sidak’s post-hoc multiple comparisons test; DL 0.2584 ± 0.0650, VL 0.1489 ± 0.0280, DM 1.4670 ± 0.3174, VM 0.6009 ± 0.1249; p < 0.0001) and before and after bath application of quinpirole (right) (repeated measures two-way ANOVA with Holm-Sidak’s post-hoc multiple comparisons test; Baseline DL 0.258 ± 0.065, QP DL 0.520 ± 0.101, Baseline VL 0.149 ± 0.028, QP VL 0.232 ± 0.051, Baseline DM 1.467 ± 0.317, QP DM 1.555 ± 0.335, Baseline VM 0.601 ± 0.125, QP VM 0.909 ± 0.170; QP treatment, p = 0.0004, GPe subregions, p < 0.0001, interaction, p = 0.2442). **n**, Summary statistics of charge ratios in the GPe subregions (left) (one-way ANOVA with Holm-Sidak’s post- hoc multiple comparisons test; DL 1.2260 ± 0.0914, VL 0.9444 ± 0.0654, DM 1.0430 ± 0.0767, VM 1.1770 ± 0.0586; p = 0.0389) and between the GPe subgroups (right) (n = 32 cells from 28 mice in the DL-VM GPe and n = 31 cells from 27 mice in the VL-DM GPe, unpaired t-test; DL-VM GPe 1.20 ± 0.0522, VL-DM GPe 0.9951 ± 0.0506). The data are presented as box-and- whisker plots or mean ± SEM. *p < 0.05, **p < 0.01, ***p < 0.001, ****p < 0.0001.

Upon application of quinpirole at striatopallidal synapses, GABAergic transmission during 10-pulse stimulation was significantly inhibited by DA receptor activation in the DL and VM GPe. However, presynaptic mechanisms partially counteracted this inhibition, sustaining short-term facilitation to mitigate frequency-dependent suppression (Fig. 5b,c,k,l,n). In contrast, oIPSCs were consistently suppressed by high-frequency stimulation in the VL GPe, where GABAergic transmission had previously been diminished by quinpirole without any change in the PPR. To assess the difference between the initial and final responses elicited by the 20 Hz stimulation, the ratio of the oIPSC amplitude for the tenth stimulus (P10) to that of the first stimulus (P1) was calculated. The lack of change in the P10/P1 ratio following quinpirole treatment suggests that DA receptor activation in the VL GPe affects striatopallidal transmission exclusively through gain modulation (Fig. 5e,f). Similarly, in the dorsomedial (DM) GPe, quinpirole attenuated GABAergic transmission, and the P10/P1 ratio remained unchanged, further indicating gain modulation. Despite this, striatopallidal synapses in the DM GPe sustained synaptic transmission by maintaining short-term facilitation, thereby functioning as a high-pass filter independent of DA receptor activation (Fig. 5h,i). Additionally, the charge ratio, which quantifies the relative suppression of trains of oIPSCs normalized to the suppression of single oIPSC^31^, differed between the DL-VM GPe and VL-DM GPe (Fig. 5n). These findings collectively suggest that ongoing activity at striatopallidal synapses is differentially shaped based on the anatomical location within the GPe. Moreover, while GABAergic transmission at striatopallidal synapses is universally suppressed by DA receptor activation throughout the GPe, pre- and postsynaptic DA receptors fine-tune this transmission in a region-specific manner.

### Dopaminergic axons are differentially denervated by 6-OHDA across the GPe subregions

Given that the dopaminergic projections from the SNc primarily follow the nigrostriatal pathway, which predominantly innervates the striatum, the degeneration of TH-positive dopaminergic axons, a key pathological hallmark of Parkinson’s disease (PD), has been extensively characterized within the striatum. However, several studies have also reported pathological alterations in the nigropallidal pathway both in human patients and animal models^33–35^. This led us to investigate whether the degeneration of dopaminergic axons in animal models of DA depletion exhibits regional heterogeneity within the GPe. To induce DA depletion pharmacologically, we unilaterally infused 6-hydroxydopamine (6-OHDA) into the medial forebrain bundle (MFB) of Adora2A-Cre;Ai9 mice (Extended Data Fig. 7a). We then examined the time-dependent denervation of DA axons in the GPe subregions at 1, 3, and 5 days post-6- OHDA administration using double immunostaining for TH and RFP (Extended Data Fig. 7b). Under these conditions, DA axons were significantly diminished in the DLS, the primary projection target of SNc DA neurons, as early as one day after the 6-OHDA injection (Extended Data Fig. 7c,f,g). Most notably, the degeneration of DA axons in the DL and VM regions of the GPe, similar to the DLS, was observed as early as one day post-6-OHDA injection. In contrast, denervation in the VL and DM GPe subregions became evident only from day 3 post-injection (Extended Data Fig. 7d). As depicted in Figure 1, the heterogeneity in DA axon innervation across GPe subregions was also preserved in the contralateral hemisphere (Extended Data Fig. 7e). These findings suggest that DA axons in different GPe subregions exhibit varying susceptibility to DA depletion, with degeneration occurring earlier in the DL and VM GPe.

### DA depletion reverses the region-specific dopaminergic modulation of striatopallidal synaptic transmission

Studies in both nonhuman primates and humans have demonstrated increased oscillatory synchronization in the GPe of Parkinsonian models and PD patients^36,37^. In addition, several studies in rodent models suggest that the heightened striatopallidal GABAergic inhibition of the GPe under DA-depleted conditions may contribute to the pathophysiological changes observed in the GPe^10,38^. We sought to determine whether dopaminergic modulation of striatopallidal transmission in the GPe is also altered in a region-specific manner under DA-depleted conditions. To unravel the physiological changes in striatopallidal transmission under DA-depleted conditions, we unilaterally injected 6-OHDA into the MFB of Adora2A-Cre;Ai32 mice (Fig. 6a). Three days post-6-OHDA infusion, we bath-applied quinpirole and measured oIPSCs and PPR changes across different GPe subregions (Fig. 6b-e). In the contralateral hemispheres, we replicated the region-specific dopaminergic modulation of striatopallidal synaptic transmission; oIPSCs decreased with an increased PPR in the DL and VM GPe, whereas oIPSCs were reduced without a change in PPR in the VL and DM GPe (Extended Data Fig. 8,9).

**Fig. 6.**
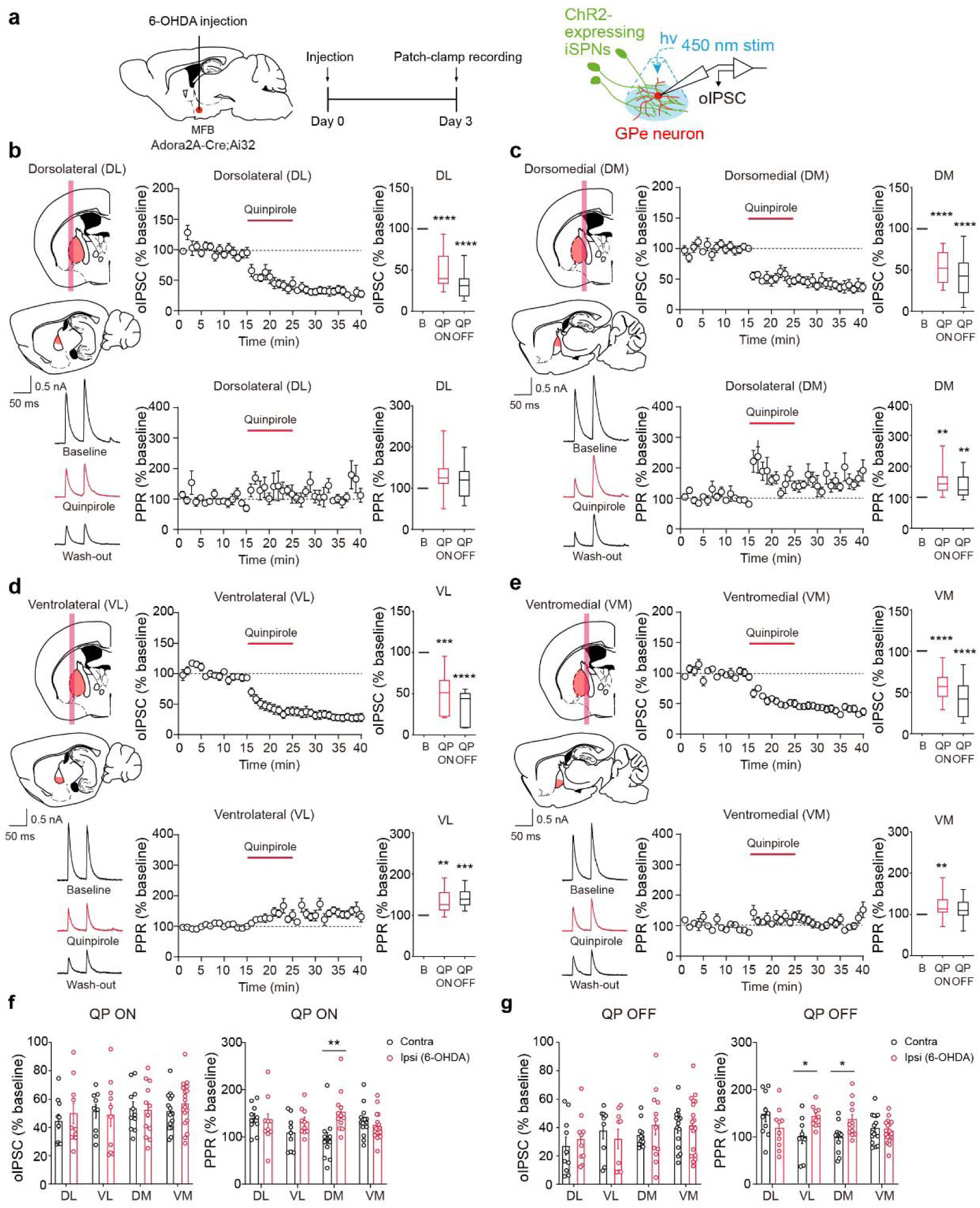
DA depletion reverses the region-specific dopaminergic modulation of striatopallidal synaptic transmission **a**, A schematic illustration describing the unilateral injection of 6-OHDA into the MFB (left) and electrophysiological recording of synaptic transmission at striatopallidal synapses via optogenetic stimulation in Adora2A-Cre;Ai32 mice (right). **b-e,** Schematic illustrations of the GPe subregions (top left), representative recording traces (bottom left), oIPSC amplitude (top center) and PPR plots (bottom center) normalized to the baseline during bath application of quinpirole (10 μM), and summary statistics of normalized oIPSCs (top right) and normalized PPRs (bottom right) in the DA-depleted condition. **b,** Summary statistics of normalized oIPSCs (n = 10 cells from 10 mice, one sample t-test; Baseline 100%, QP ON 49.95 ± 7.23%, QP OFF 31.65 ± 5.41%) and normalized PPRs (Baseline 100%, QP ON 132.70 ± 15.88%, QP OFF 117.40 ± 13.59%) in the DL GPe. **c,** Summary statistics of normalized oIPSCs (n = 12 cells from 12 mice, one sample t-test; Baseline 100%, QP ON 52.51 ± 5.44%, QP OFF 41.80 ± 7.14%) and normalized PPRs (Baseline 100%, QP ON 153.30 ± 12.68%, QP OFF 136.0 ± 10.86%) in the DM GPe. **d,** Summary statistics of normalized oIPSCs (n = 10 cells from 9 mice, one sample t-test; Baseline 100%, QP ON 48.86 ± 8.03%, QP OFF 31.79 ± 6.97%) and normalized PPRs (Baseline 100%, QP ON 132.50 ± 9.07%, QP OFF 142.50 ± 7.04%) in the VL GPe. **e,** Summary statistics of normalized oIPSCs (n = 18 cells from 16 mice, one sample t-test; Baseline 100%, QP ON 56.77 ± 3.77%, QP OFF 41.40 ± 4.70%) and normalized PPRs (Baseline 100%, QP ON 122.0 ± 6.69%, QP OFF 111.70 ± 5.74%) in the VM GPe. **f,** Summary statistics of normalized oIPSC amplitudes (left) (ordinary two-way ANOVA with Holm-Sidak’s post-hoc multiple comparisons test; Contra DL 44.347 ± 5.088%, Ipsi DL 49.954 ± 7.226%, Contra VL 51.447 ± 4.930%, Ipsi VL 48.861 ± 8.030%, Contra DM 53.383 ± 4.646%, Ipsi DM 52.514 ± 5.437%, Contra VM 51.443 ± 3.434%, Ipsi VM 56.773 ± 3.770%; GPe subregion, p = 0.5448, 6-OHDA, p = 0.6188, interaction, p = 0.8144) and normalized PPRs (right) (ordinary two-way ANOVA with Holm-Sidak’s post-hoc multiple comparisons test; Contra DL 139.859 ± 9.380%, Ipsi DL 132.732 ± 15.879%, Contra VL 108.932 ± 12.277%, Ipsi VL 132.508 ± 9.071%, Contra DM 95.106 ± 13.944%, Ipsi DM 153.346 ± 12.678%, Contra VM 134.012 ± 8.134%, Ipsi VM 122.040 ± 6.694%; GPe subregion, p = 0.5846, 6-OHDA, p = 0.0470, interaction, p = 0.0042) during QP ON state. **g,** Summary statistics of normalized oIPSC amplitudes (left) (ordinary two-way ANOVA with Holm-Sidak’s post-hoc multiple comparisons test; Contra DL 26.774 ± 6.324%, Ipsi DL 31.651 ± 5.413%, Contra VL 37.657 ± 5.798%, Ipsi VL 31.790 ± 6.968%, Contra DM 34.370 ± 2.790%, Ipsi DM 41.805 ± 7.140%, Contra VM 39.535 ± 3.705%, Ipsi VM 41.398 ± 4.70%; GPe subregion, p = 0.1840, 6-OHDA, p = 0.5974, interaction, p = 0.7010) and normalized PPRs (right) (ordinary two-way ANOVA with Holm-Sidak’s post-hoc multiple comparisons test; Contra DL 147.0 ± 14.870%, Ipsi DL 117.371 ± 13.592%, Contra VL 99.991 ± 14.058%, Ipsi VL 142.466 ± 7.039%, Contra DM 97.972 ± 9.605%, Ipsi DM 135.955 ± 10.857%, Contra VM 117.347 ± 8.066%, Ipsi VM 111.744 ± 5.741%; GPe subregion, p = 0.3303, 6- OHDA, p = 0.1248, interaction, p = 0.002) during QP OFF state. The data are presented as box-and-whisker plots or mean ± SEM. *p < 0.05, **p < 0.01, ***p < 0.001, ****p < 0.0001.

Upon quinpirole treatment, oIPSCs were similarly decreased across all ipsilateral GPe subregions, mirroring the effects observed in the contralateral GPe (Fig. 6f). Interestingly, we found that the alterations in PPR following quinpirole treatment shifted in opposite directions between the DL-VM and VL-DM subregions of the ipsilateral GPe compared to the contralateral GPe (Fig. 6f,g). For instance, DA depletion negated the quinpirole-induced elevation of PPR in striatopallidal GABAergic transmission in the DL GPe (Fig. 6b and Extended Data Fig 9a,b). Moreover, the effect of quinpirole on PPR changes became less pronounced following DA depletion in the VM GPe (Fig. 6e and Extended Data Fig 9g,h). On the other hand, striatopallidal synapses in the VL and DM GPe, where PPR remained unchanged by quinpirole under normal conditions, unexpectedly began to exhibit increased PPR under DA-depleted conditions (Fig. 6c,d and Extended Data Fig 9c-f). Interestingly, during the baseline recording, the PPR was markedly diminished in the VL GPe of the ipsilateral hemisphere (Extended Data Fig 9c,d,i,j). In contrast, striatopallidal synapses in the DM region of the GPe, which sustained ongoing activity through gain modulation alone, showed an enhancement of PPR after quinpirole treatment, without any alterations in baseline PPR (Extended Data Fig. 9e,f,i,j). These findings collectively suggest that dopaminergic modulation of striatopallidal short-term plasticity (STP) is differentially affected by DA depletion in the DL-VM and VL-DM subregions of the GPe.

### DA depletion induces region-specific alterations in the spatial distribution of presynaptic D2Rs at striatopallidal axon terminals

In light of the possibility that variations in the spatial distribution and localization of D2Rs at striatopallidal axon terminals may contribute to region-specific dopaminergic modulation of striatopallidal transmission within the GPe, we further investigated whether DA depletion induces region-specific alterations in the spatial distribution and localization of D2Rs. Striatopallidal axons and axon terminals were labeled with GFP and RFP (mRuby), respectively, by injecting AAVs into the entire striatum (Fig. 7a). Following 6-OHDA injection, we performed multi-color immunostaining and utilized enhanced confocal imaging with airyscan in each subregion of the GPe (Fig. 7b). We replicated our findings from Figure 4, demonstrating that D2Rs were more localized in the striatopallidal axons and axon terminals within the DL and VM GPe compared to the VL and DM GPe in the contralateral hemisphere (Fig. 7c). To further delve into the spatial relationship between presynaptic D2Rs and striatopallidal axon terminals in each subregion of the GPe, we conducted clustering analysis using Ripley’s H function, which is particularly sensitive to spatial point patterns^39,40^. When the nearest distances from striatopallidal axon terminals (RFP+) to D2Rs in the VL and DM subregions of the GPe were used as a null model, clustering analysis revealed that the DL and VM GPe exhibited a greater tendency for clustered features between striatopallidal axon terminals and D2Rs (Fig. 7e). The results were reversed when the null model was replaced by the nearest distances from striatopallidal axon terminals to D2Rs in the DL and VM GPe, indicating that striatopallidal axon terminals and D2Rs appeared to be less clustered in the VL and DM GPe (Fig. 7e).

**Fig. 7.**
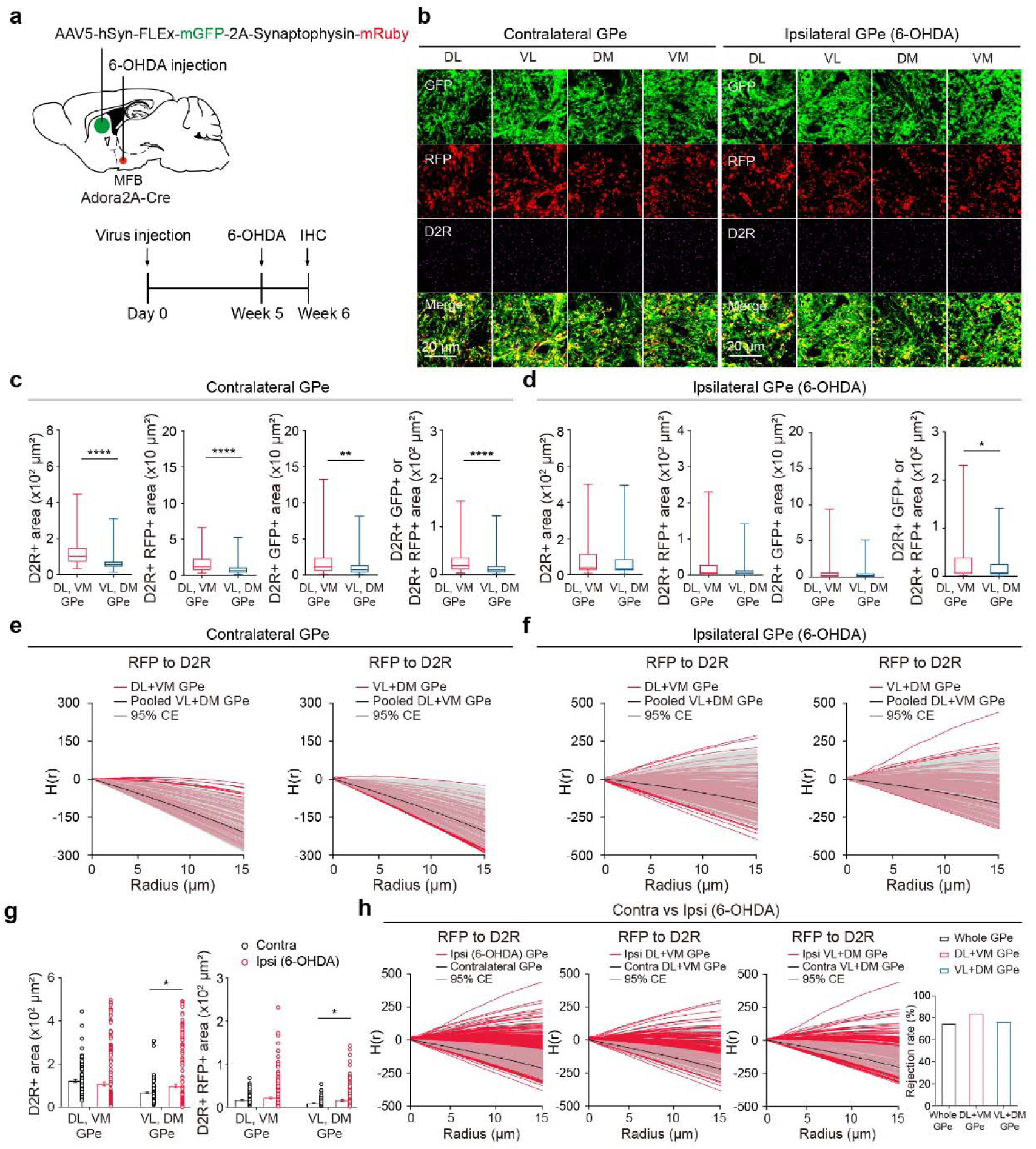
DA depletion induces region-specific alterations in the spatial distribution of presynaptic D2Rs at striatopallidal axon terminals **a**, A schematic illustration depicting the injection of AAV5-hSyn-Flex-mGFP-2A- Synaptophysin-mRuby virus into the striatum of Adora2A-Cre mice, along with the unilateral injection of 6-OHDA into the MFB. **b**, Representative enhanced confocal images of striatopallidal axon terminals and D2Rs in the GPe subregions of the contralateral and ipsilateral (6-OHDA) hemispheres. **c,** Summary statistics of D2R+ area (n = 120 images from 11 mice per GPe subgroup, unpaired t-test; DL-VM GPe 122.0 ± 6.91 μm^2^, VL-DM GPe 67.76 ± 4.38 μm^2^), D2R+ area colocalized with RFP+ area (DL-VM GPe 16.23 ± 1.11 μm^2^, VL-DM GPe 8.83 ± 0.73 μm^2^), D2R+ area colocalized with GFP+ area (DL-VM GPe 19.92 ± 2.11 μm^2^, VL-DM GPe 12.29 ± 1.30 μm^2^), and D2R+ area colocalized with GFP+ or RFP+ area (DL-VM GPe 27.79 ± 2.34 μm^2^, VL-DM GPe 15.97 ± 1.65 μm^2^) between GPe subgroups in the contralateral hemisphere. **d,** Summary statistics of D2R+ area (n = 120 images from 11 mice per GPe subgroup, unpaired t-test; DL-VM GPe 104.0 ± 9.10 μm^2^, VL-DM GPe 97.36 ± 9.25 μm^2^), D2R+ area colocalized with RFP+ area (DL-VM GPe 21.56 ± 2.44 μm^2^, VL-DM GPe 15.94 ± 1.85 μm^2^), D2R+ area colocalized with GFP+ area (DL-VM GPe 6.18 ± 0.83 μm^2^, VL-DM GPe 5.09 ± 0.55 μm^2^), and D2R+ area colocalized with GFP+ or RFP+ area (DL-VM GPe 25.74 ± 2.44 μm^2^, VL-DM GPe 18.91 ± 1.83 μm^2^) between GPe subgroups in the ipsilateral hemisphere. **e,** Edge-corrected Ripley’s H function analysis of the distance from RFP to D2R in the DL-VM GPe and VL-DM GPe (null model, CE: confidence envelope) (left), and the distance from RFP to D2R in the VL-DM GPe and DL-VM GPe (null model, CE: confidence envelope) (right) in the contralateral GPe. **f,** Edge-corrected Ripley’s H function analysis of the distance from RFP to D2R in the DL-VM GPe and VL-DM GPe (null model, CE: confidence envelope) (left), and the distance from RFP to D2R in the VL-DM GPe and DL-VM GPe (null model, CE: confidence envelope) (right) in the ipsilateral GPe. **g,** Summary statistics of D2R+ area (left) (ordinary two- way ANOVA with Holm-Sidak’s post-hoc multiple comparisons test; Contra DL-VM 122.01 ± 6.91 μm^2^, Ipsi DL-VM 107.96 ± 9.16 μm^2^, Contra VL-DM 67.76 ± 4.38 μm^2^, Ipsi VL-DM 97.36 ± 9.25 μm^2^; GPe subgroup, p = 0.0004, 6-OHDA, p = 0.3946, interaction, p = 0.0171) and D2R+ area colocalized with RFP+ area (right) (ordinary two-way ANOVA with Holm-Sidak’s post-hoc multiple comparisons test; Contra DL-VM 16.23 ± 1.11 μm^2^, Ipsi DL-VM 21.56 ± 2.44 μm^2^, Contra VL-DM 8.83 ± 0.73 μm^2^, Ipsi VL-DM 15.94 ± 1.85 μm^2^; GPe subgroup, p = 0.0020, 6- OHDA, p = 0.0032, interaction, p = 0.6695) comparing the contralateral and ipsilateral hemispheres. **h,** Edge-corrected Ripley’s H function analysis of the distance from RFP to D2R in the ipsilateral GPe and contralateral GPe (null model, CE: confidence envelope) (left), the distance from RFP to D2R in the DL-VM GPe of the ipsilateral hemisphere and contralateral hemisphere (null model, CE: confidence envelope) (center), and the distance from RFP to D2R in VL-DM GPe in the ipsilateral hemisphere and contralateral hemisphere (null model, CE: confidence envelope). Rejection rate of the distance from RFP to D2R in the ipsilateral hemisphere, evaluated using the DCLF test (compared to the null model of the contralateral hemisphere) (right). The data are presented as box-and-whisker plots or mean ± SEM. *p < 0.05, **p < 0.01, ****p < 0.0001.

However, following DA depletion induced by 6-OHDA infusion, the differences in D2R localization at striatopallidal axons and axon terminals between the DL-VM and VL-DM subregions of the ipsilateral GPe were no longer observed (Fig. 7d). Under this pathological condition, the difference in the level of clustering between striatopallidal axon terminals and D2Rs also became less pronounced between the DL-VM and VL-DM subregions of the GPe (Fig. 7f). Interestingly, following DA depletion, the expression of D2Rs on striatopallidal axon terminals increased significantly in the VL and DM GPe subregions (Fig. 7g). Moreover, the post-DA depletion led to a marked enhancement in the clustering of striatopallidal axon terminals and D2Rs in the ipsilateral GPe, in contrast to the contralateral GPe. This finding was further supported by rejection rates exceeding 70%, calculated as the proportion of samples for which the null model was rejected using the Diggle-Cressie-Loosmore-Ford (DCLF) test^41^ (Fig. 7h).

Under the same experimental conditions (Extended Data Fig. 10a), we also examined the spatial relationship between dopaminergic boutons and striatopallidal axon terminals after DA depletion. The co-localization of TH and vesicular monoamine transporter 2 (VMAT2) immunofluorescence was considered indicative of potential dopaminergic boutons, while RFP (mRuby)-positive fluorescence was used to identify putative striatopallidal axon terminals (Extended Data Fig. 10b,c). We found no changes in the area of striatopallidal axons (GFP+) or axon terminals (RFP+) within the GPe following DA depletion (Extended Data Fig. 10d,e). Notably, in the contralateral hemisphere, more contact areas between dopaminergic boutons and striatopallidal axon terminals (or axons) were observed in the DL and VM GPe compared to the VL and DM GPe. This regional difference, however, was absent in the 6-OHDA-lesioned ipsilateral hemisphere (Extended Data Fig. 10f). In the ipsilateral hemisphere, the areas of both dopaminergic axons and boutons were significantly reduced across all GPe subregions (Extended Data Fig. 10g). Furthermore, no significant differences in the nearest neighbor distances between dopaminergic boutons and striatopallidal axon terminals were observed among the GPe subregions, nor were there significant changes in the spatial relationship between striatopallidal axon terminals (Extended Data Fig. 10h,i). Taken together, our findings reveal that denervation of dopaminergic axons and DA depletion can lead to region-specific alterations in the expression, localization, and spatial distribution of D2Rs on striatopallidal axon terminals within the GPe. These changes may contribute to the functional modifications observed in the dopaminergic modulation of striatopallidal transmission.

### DA depletion may lead to region-specific changes in calcium dynamics at striatopallidal synapses

To further investigate the physiological mechanisms underlying the heterogeneous dopaminergic modulation of striatopallidal STP under both control and DA-depleted conditions, we utilized a computational modeling approach based on the Tsodyks-Markram model, which describes the dynamics of STP under general conditions^42,43^ (Fig. 8a). Consistent with our quantal analysis (Fig. 3a-e), the computational model successfully reproduced the experimentally observed subregion-specific patterns of dopaminergic modulation by incorporating the calcium channel open probability as a key variable modulated by quinpirole^44,45^ (Fig. 8b). Under control conditions, the computational model predicted a selective decrease in calcium channel open probability and intracellular calcium concentration within the striatopallidal axon terminals of the DL and VM GPe subregions (Fig. 8c-f). In contrast, under DA-depleted conditions, the model indicated a marked reduction in calcium channel open probability and calcium concentration across all GPe subregions, except for the DL GPe (Fig. 8g-j). Together, these results suggest that DA depletion drives distinct synaptic adaptations at striatopallidal synapses depending on their spatial locations within the GPe, further highlighting the region-specific mechanisms that govern synaptic transmission in this nucleus.

**Fig. 8.**
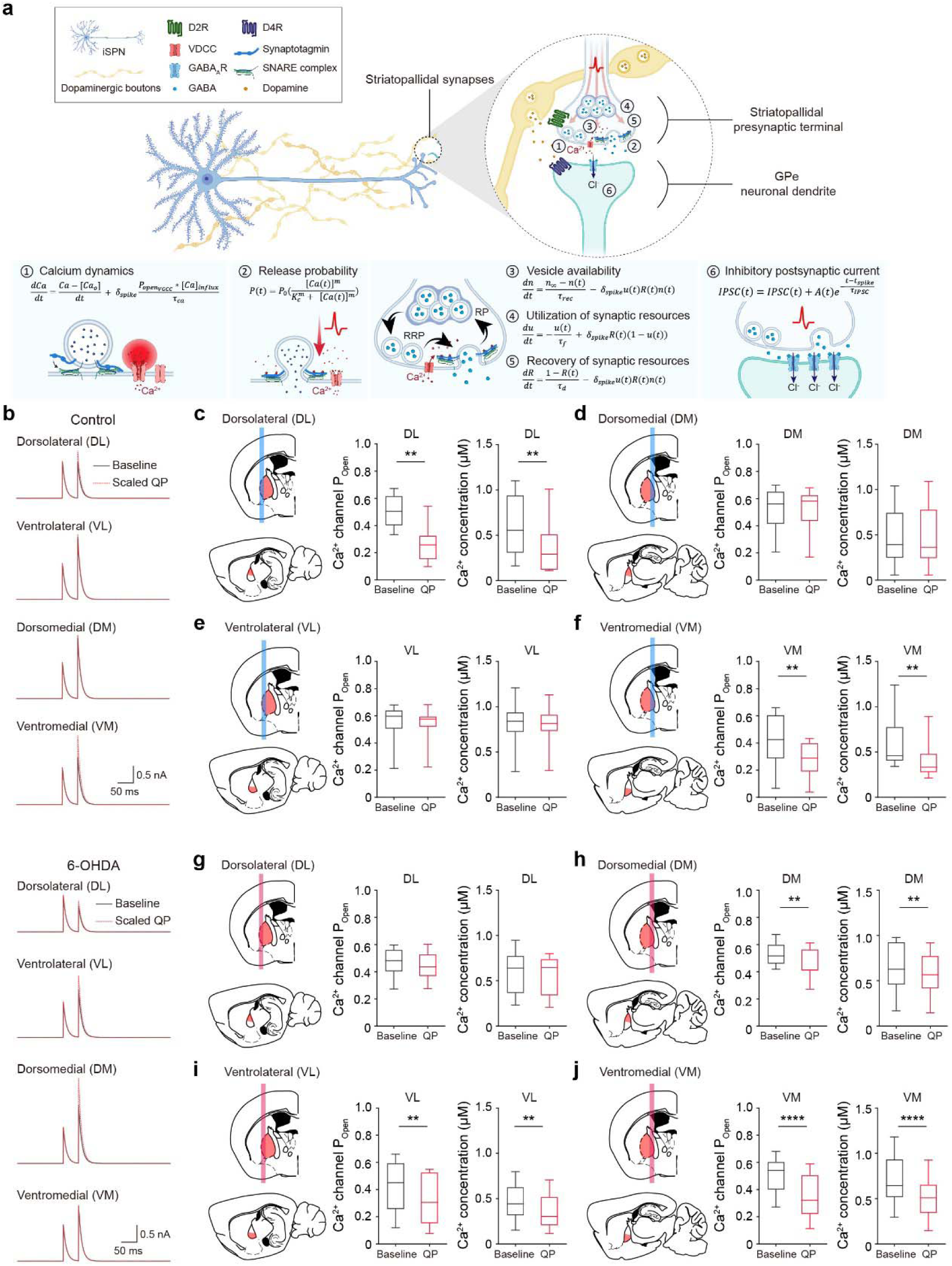
Computational modeling of striatopallidal synapses indicates region-specific changes in calcium dynamics **a,** A schematic illustration depicting the mathematical framework of computational modeling at striatopallidal synapses. **b,** Representative simulated PPR traces of GPe subregions under control conditions (Control) and DA-depleted conditions (6-OHDA). **c-f,** Schematic illustrations of the GPe subregions (left), summary statistics of simulated Ca²^+^ channel open probability (middle), and Ca²^+^ concentration at the axon terminals (right) under control conditions. **c,** Summary statistics of simulated Ca²^+^ channel open probability (n = 10, Wilcoxon matched pairs signed rank test; Baseline 0.5097 ± 0.0349, QP 0.2664 ± 0.0405) and Ca²^+^ concentration (n = 10, Wilcoxon matched pairs signed rank test; Baseline 0.6020 ± 0.1056 μM, QP 0.3679 ± 0.0908 μM) in the DL GPe. **d,** Summary statistics of simulated Ca²^+^ channel open probability (n = 10, Wilcoxon matched pairs signed rank test; Baseline 0.5167 ± 0.0479, QP 0.5203 ± 0.0486) and Ca²^+^ concentration (n = 10, Wilcoxon matched pairs signed rank test; Baseline 0.4880 ± 0.1019 μM, QP 0.4960 ± 0.1090 μM) in the DM GPe. **e,** Summary statistics of simulated Ca²^+^ channel open probability (n = 10, Wilcoxon matched pairs signed rank test; Baseline 0.5549 ± 0.0434, QP 0.5390 ± 0.0389) and Ca²^+^ concentration (n = 10, Wilcoxon matched pairs signed rank test; Baseline 0.8195 ± 0.0764 μM, QP 0.7952 ± 0.0680 μM) in the VL GPe. **f,** Summary statistics of simulated Ca²^+^ channel open probability (n = 10, Wilcoxon matched pairs signed rank test; Baseline 0.4178 ± 0.0614, QP 0.2778 ± 0.0421) and Ca²^+^ concentration (n = 10, Wilcoxon matched pairs signed rank test; Baseline 0.5936 ± 0.0883 μM, QP 0.4043 ± 0.0636 μM) in the VM GPe. **g-j,** Schematic illustrations of the GPe subregions (left), summary statistics of simulated Ca²^+^ channel open probability (middle), and Ca²^+^ concentration at the axon terminals (right) under DA-depleted conditions. **g,** Summary statistics of simulated Ca²^+^ channel open probability (n = 9, Wilcoxon matched pairs signed rank test; Baseline 0.4732 ± 0.0353, QP 0.4407 ± 0.0354) and Ca²^+^ concentration (n = 9, Wilcoxon matched pairs signed rank test; Baseline 0.5746 ± 0.0812 μM, QP 0.5404 ± 0.0736 μM) in the DL GPe. **h,** Summary statistics of simulated Ca²^+^ channel open probability (n = 9, Wilcoxon matched pairs signed rank test; Baseline 0.5309 ± 0.0280, QP 0.4528 ± 0.0358) and Ca²^+^ concentration (n = 9, Wilcoxon matched pairs signed rank test; Baseline 0.6650 ± 0.0935 μM, QP 0.5680 ± 0.0802 μM) in the DM GPe. **i,** Summary statistics of simulated Ca²^+^ channel open probability (n = 9, Wilcoxon matched pairs signed rank test; Baseline 0.4341 ± 0.0639, QP 0.3318 ± 0.0608) and Ca²^+^ concentration (n = 9, Wilcoxon matched pairs signed rank test; Baseline 0.4544 ± 0.0682 μM, QP 0.3648 ± 0.0651 μM) in the VL GPe. **j,** Summary statistics of simulated Ca²^+^ channel open probability (n = 17, Wilcoxon matched pairs signed rank test; Baseline 0.5001 ± 0.0314, QP 0.3607 ± 0.0370) and Ca²^+^ concentration (n = 17, Wilcoxon matched pairs signed rank test; Baseline 0.6953 ± 0.0625 μM, QP 0.5048 ± 0.0506 μM) in the VM GPe. The data are presented as box-and-whisker plots. **p < 0.01, ****p < 0.0001.

## Discussion

We demonstrate here that information transmitted via the indirect pathway can be modulated through presynaptic D2Rs and postsynaptic D4Rs in a subregion-specific manner within the GPe. DA depletion exerts differential and region-specific effects on the presynaptic modulation of GABA release mediated by D2Rs. The GPe, previously considered a simple relay station in the indirect pathway, possesses a heterogeneous neuronal composition with distinct characteristics and functions^46–49^. These findings unveil a previously unrecognized role of dopaminergic modulation in the GPe, highlighting its complex, anatomically specific effects on striatopallidal synaptic transmission.

The distinct DA modulation of striatopallidal transmission within the GPe subregions through D2-like DA receptors was topographically organized in a pinwheel-like fashion along two orthogonal, diagonal axes: the DL-VM and VL-DM orientation. This anatomical organization is particularly intriguing when considered alongside a recent work demonstrating that GPe projections to the subthalamic nucleus (STN) are also topographically aligned linearly along two diagonal gradient axes^50^. Given the parallel connectivity of the indirect pathway within the cortex-basal ganglia-thalamic network^13^ the topographic alignment of dopaminergic modulation in the GPe subregions may play a critical role in shaping information processing in the downstream nuclei of the basal ganglia circuitry.

Monitoring DA with fast-scan cyclic voltammetry (FSCV) has been challenging due to the limited sensitivity of electrochemical techniques. The fluorescent DA sensor GRAB_DA_, allowed for direct monitoring of DA release from the nigropallidal axons in the GPe. We took advantage of optogenetic stimulation to selectively drive striatopallidal transmission. The most intriguing finding in our investigation of striatopallidal transmission is that, while the D2-like DA receptor agonist quinpirole ubiquitously attenuated GABAergic transmission at striatopallidal synapses, it increased the PPR only in the DL and VM GPe. In contrast, the PPR remained unaffected in the VL and DM regions of the GPe. By analyzing the effects of quinpirole on quantal properties of striatopallidal transmission across the GPe subregions and employing D2R- and D4R-selective antagonists, we discovered that postsynaptic D4Rs at striatopallidal synapses mediate the suppression of GABAergic transmission irrespective of the anatomical locations within the GPe. However, presynaptic D2Rs effectively inhibited presynaptic GABA release exclusively in the DL and VM GPe. Regional differences in the expression, localization, and spatial distribution of presynaptic D2Rs and postsynaptic D4Rs at striatopallidal synapses may thus represent the mechanisms accounting for the distinct dopaminergic modulations observed across the GPe subregions. The efficacy of presynaptic D2R activation and its signal transduction may also vary depending on the alternatively spliced isoforms of D2Rs expressed on striatopallidal axon terminals in each subregion of the GPe^51,52^. Moreover, presynaptic modulation of total calcium influx via D2Rs could exhibit significant divergence across the GPe subregions.

Various STP regulation modes have been identified across several brain regions, including the cortex^53^, cerebellum^54^, and hippocampus^55^. In addition, both canonical and noncanonical presynaptic modulation can coexist at the same synapses in the prefrontal cortex^31^, suggesting the presence of diverse mechanisms for precisely and finely tuning synaptic functions. Striatopallidal synapses, serving as the initial gateway of the indirect pathway, represent a critical node that can profoundly influence the overall output of this pathway. Synaptic DA modulation may become even more diverse when ongoing synaptic activity is propagated through the striatopallidal synapses. When stimulated with trains of light pulses, the entire sequence of oIPSCs progressively depressed in the lateral GPe, including the DL and VL regions. Moreover, stimulation of D2-like receptors with quinpirole led to an additional suppression of striatopallidal transmission in the DL and VL GPe. However, STP at striatopallidal synapses in the DL GPe allowed ongoing activity to be sustained toward the end of the stimulation. In contrast, synaptic activities in the VL GPe were continuously suppressed, with the last oIPSC showing the strongest attenuation.

In the case of medial GPe, including the DM and VM regions, continuing activity facilitated synaptic transmission at striatopallidal synapses. This facilitation was observed throughout the entire sequence of all ten oIPSCs in the DM GPe or at least during the first half of the synaptic transmission in the VM GPe. Although the activation of D2-like receptors in the medial GPe universally attenuated GABAergic transmission, as observed in the lateral GPe, a high-pass filtering mechanism consistently promoted synaptic transmission in the DM GPe. Notably, this facilitation occurred independently of D2-like receptor activation. Neurons in the GPe also receive excitatory inputs directly from the STN and indirectly from the cortex. Given the facilitating nature of GABAergic transmission at striatopallidal synapses in the DM GPe, it is likely that only robust excitatory inputs are capable of triggering the firing of GPe neurons in this region. In contrast to the DM GPe, presynaptic D2Rs in the VM regions of the GPe function as a high-pass filter on striatopallidal transmission, effectively sustaining ongoing activity.

Our findings collectively suggest that ongoing synaptic information transmitted through striatopallidal synapses is differently shaped along the lateral-to-medial axis of the GPe. Simultaneously, presynaptic mechanisms mediated by D2Rs sustain continuing synaptic activity, particularly in the DL and VM regions of the GPe. It is important to note that although high-frequency, ongoing synaptic activity was gain-modulated without STP in both the VL and DM regions of the GPe, the effects of this modulation differed. In the VL GPe, the synaptic gain was modulated to primarily inhibit striatopallidal transmission, whereas gain modulation functioned to augment synaptic transmission in the DM regions of the GPe. In light of the recent studies revealing topographically graded organization in the basal ganglia circuit^13,50^, it is very likely that diverse modalities of information, including sensory, motor, associative, and limbic inputs, may propagate in parallel through the basal ganglia network. Supporting this, diffusion tensor imaging (DTI) studies in human subjects have shown the existence of three parallel channels originating from association, motor, and limbic cortices, which project to distinct striatal regions^56^. Building on these findings, our study offers new insights into how distinct cortical information may be differentially processed and shaped within anatomically parallel subnetworks of the basal ganglia, particularly through the indirect pathway.

A comprehensive understanding of the anatomical organization and functional features of dopaminergic modulation within the basal ganglia circuitry, including striatopallidal synapses in the GPe, is essential for elucidating the pathophysiology of DA-related brain disorders. We found that dopaminergic projections through the nigropallidal pathway exhibit varying susceptibility to DA depletion depending on their specific innervation sites; DA axons in the DL and VM GPe are more vulnerable to depletion than those in the VL and DM GPe. Given the heterogeneous molecular characteristics of DA neurons in the midbrain, this differential susceptibility of nigropallidal DA axons to DA depletion may stem from their distinct origins within specific DA neuronal subtypes^57–59^.

In 6-OHDA-lesioned mice, striatopallidal synapses in the VL and DM subregions of the GPe, where GABAergic transmission was initially suppressed by the activation of D2-like receptors without changes in the PPR, exhibited a significant elevation in PPR following treatment with quinpirole. However, these alterations in PPR observed in the VL and DM GPe may arise from distinct mechanisms. Considering the observed decrease in baseline PPR after DA depletion and the absence of a significant difference in absolute PPR values between ipsilateral and contralateral hemispheres following quinpirole application, it is likely that the enhanced PPR in the VL GPe is due to altered presynaptic release probability resulting from DA depletion^60^. In the case of striatopallidal transmission in the DM GPe under DA-depleted conditions, the PPR in the ipsilateral GPe showed a marked increase following quinpirole application despite no baseline PPR differences between hemispheres. This result potentially indicates either D2R upregulation or increased D2R supersensitivity due to DA depletion^59,61,62^.

Furthermore, the enhanced STP at striatopallidal synapses in the DM GPe, induced by D2- like receptor activation, may raise the activation threshold of GPe neurons. Notably, in contrast to the changes in the VL and DM GPe, the increase in PPR by quinpirole treatment was attenuated after DA depletion in the DL and VM subregions of the GPe. These findings, together with the observations in the VL and DM GPe, suggest that subregion-specific alterations in the expression, localization, and spatial distribution of D2Rs on striatopallidal axon terminals within the GPe may partially, though not entirely, account for the physiological changes in the DL and VM GPe under DA-depleted conditions.

Expanding on these findings, computational modeling revealed subregion-specific differences in calcium dynamics under DA-depleted conditions. By incorporating calcium channel open probability as a critical variable modulated by quinpirole, the modeling outcomes effectively recapitulated the experimentally observed differences in PPR, reinforcing the validity of our experimental findings across GPe subregions. Notably, the modeling indicated that the VM GPe retained calcium channel modulation in response to quinpirole under DA-depleted conditions. Although our electrophysiological data revealed a trend toward reduced PPR in both the DL and VM GPe under dopamine depletion, only the VM GPe maintained a statistically significant elevation of PPR in response to quinpirole. Given that DL and VM GPe exhibited weakened spatial clustering between striatopallidal axon terminals and D2Rs following DA depletion, our electrophysiological recordings and computational modeling underscore the intricate nature of striatopallidal synaptic adaptations to DA depletion. The ability of the VM GPe to preserve PPR modulation by quinpirole under DA-depleted conditions may be attributed to its sustained regulation of calcium dynamics, as suggested by the modeling results. Furthermore, AI-based clustering analysis of Sr² -induced quantal responses revealed that the VM GPe exhibits distinct quantal properties compared to other subregions. These findings suggest that the release machinery within the striatopallidal axon terminals of the VM GPe may possess specialized characteristics that differentiate it from other subregions. Our findings demonstrate that striatopallidal synapses within the GPe exhibit dynamic, region-specific responses to DA depletion, reflecting the functional heterogeneity of GPe subregions. Our data also indicate the spatially organized and functionally diverse roles of GPe subregions in maintaining information processing within the basal ganglia under pathological conditions.

In summary, our findings provide the first evidence that sensory-motor-associative information transmitted via the indirect pathway can be differentially modulated by DA receptor activation under both normal and DA-depleted conditions, depending on the anatomical locations of striatopallidal synapses along the dorsoventral and lateromedial axes of the GPe. Our data further suggest that region-specific differences in the expression, localization, spatial distribution, and functional properties of presynaptic D2Rs and postsynaptic D4Rs at striatopallidal synapses may underlie the distinct dopaminergic modulation observed across different subregion of the GPe. These results provide novel insights into the functional role of nigropallidal pathway and its dopaminergic modulation of striatopallidal synapses within the GPe in both health and disease.

## Methods

### Animals

All experimental procedures were conducted by protocols approved by the Institutional Animal Care and Utilization Committee of the Ulsan National Institute of Science and Technology (UNIST). All mice were maintained in C57BL/6J background. Mice were group-housed under a 12 hr light / dark schedule (lights on from 6 AM to 6 PM) and given *ad libitum* access to food and water. Mice were group-housed (up to 5 mice per cage) and bred under standard pathogen-free housing conditions in the animal facility of UNIST. The majority of experiments were performed using male and female mice aged 10 to 15 weeks. Given the health and condition of the GPe neurons in acute brain slices, patch clamp recording experiments were conducted on mice aged 3 to 4 weeks. To specifically express ChR2 in indirect pathway medium spiny neurons (iMSNs), Adora2A-Cre (B6.FVB(Cg)-Tg(Adora2a-cre)KG139Gsat/Mmucd, MMRRC stock number: 036158-UCD) mice were crossed with Ai32 (B6.Cg-Gt(ROSA)26Sor^tm32(CAG-COP4*H134R/EYFP)Hze^/J, Jackson stock number: 024109) mice, producing Adora2A-Cre;Ai32. These mice were used for optogenetic stimulation of striatopallidal synaptic terminals in whole-cell patch clamp recording experiments. To visualize striatopallidal pathway (axons and axon terminals), Adora2A-Cre;Ai9 mice were generated by crossing Ai9-tdTomato (B6.Cg-Gt(ROSA)26Sor^tm9(CAG-tdTomato)Hze^/J) mice with Adora2A-Cre mice. These mice were also used in immunohistochemistry experiments. Adora2A-Cre mice were used for immunohistochemistry and fluorescence imaging experiments.

### One-photon fluorescence imaging in the GPe in acute brain slices

Adult C57BL/6J mice were anesthetized via intraperitoneal injection of zoletil (60 mg/kg, Virbac Korea) and rompun (15 mg/kg, Bayer Korea) mixture solution (zoletil : rompun : saline = 4 : 1 : 20), and AAVs expressing GRAB_DA_ (AAV9-hSyn-rDA1.2a, 3.28 × 10^13^ vg/ml, WZ Biosciences) were injected (250 nl per injection site at a rate of 100 nl/min) into the GPe (coordinates used, AP: -0.45 mm from the bregma, ML: ±2.6 mm from the bregma, DV: −3.5 mm, -3.9 mm from the dura). Three weeks after virus injection, the mice were deeply anesthetized with isoflurane (Piramal Critical Care), then decapitated and the brains were immediately removed and briefly exposed to ice-chilled artificial cerebrospinal fluid (ACSF) containing 125 mM NaCl, 2.5 mM KCl, 1.25 mM NaH_2_PO_4_, 25 mM NaHCO_3_, 1 mM MgCl_2_, 2mM CaCl_2_ and 15 mM glucose oxygenated with 95% O_2_ and 5% CO_2_. Acute brain slices were prepared as 300-μm-thick sagittal sections using a tissue vibratome (Leica, VT 1200S). One-photon imaging was performed using a BX51WI microscope (Olympus) equipped with a 40x/0.8 NA water-immersion objective (Olympus) and a SOLIS-565C high-power LED (Thorlabs). A 565 nm LED was used to excite the rDA1.2a sensor, and fluorescence was collected using a 569–594 nm filter (Chroma). Fluorescence images were further acquired using a scientific complementary metal-oxide-semiconductor (sCMOS) camera (Orca-Flash4.0 V3, Hamamatsu) in conjunction with HCImage Live software (Hamamatsu). For electrical stimulation, a bipolar concentric electrode (CBAPB75, FHC) was positioned on the GPe under fluorescence guidance. Dopamine release was evoked by electrical stimulation (0.2 ms pulse duration, ten pulses), controlled using a DS-3 stimulus isolator (Digitimer, UK). Dopamine release were imaged at a resolution of 512 x 512 pixels/frame, 10 frames/s, with an exposure time of 10 ms. For image analyses, regions of interest (ROIs) expressing rDA1.2a sensor were manually selected and analyzed using ImageJ software (NIH).

### Immunohistochemistry and confocal imaging

Mice were deeply anesthetized by intraperitoneal injection of zoletil (60 mg/kg, Virbac Korea) and rompun (15 mg/kg, Bayer Korea) mixture solution (zoletil : rompun : saline = 4 : 1 : 20) and perfused transcardially with phosphate buffer (PB), followed by 4% paraformaldehyde (PFA, Sigma-Aldrich). Brains were rapidly removed and postfixed in 4% PFA at 4°C for overnight. Fixed brains were transferred to 30% sucrose in 0.01 M PB for cryoprotection. Brain sections were made into 20 μm thick sagittal slices using frozen section technique (Microtome SM2010, Leica). Obtained free-floating brain sections were washed with PBS, PBST (0.5% Triton X-100) and blocked with PBST containing 10% normal goat serum (NGS, Sigma-Aldrich) and 2% bovine serum albumin (BSA, Sigma-Aldrich). After blocking, brain sections were incubated with an anti-tyrosine hydroxylase antibody (rabbit polyclonal, 1:1000, ab112, Abcam), an anti-tyrosine hydroxylase antibody (chicken polyclonal, 1:500, ab76442, Abcam), an anti-RFP antibody (Guinea pig polyclonal, 1:1000, 390 005, Synaptic Systems), an anti-RFP antibody (mouse monoclonal, 1:1000, MA5-15257, Thermo Fisher Scientific), an anti-GFP antibody (chicken polyclonal, 1:1000, GFP-1010, Aves Labs), an anti-GFP antibody (mouse monoclonal, 1:1000, sc-9996, Santa Cruz Biotechnology), an anti-bassoon antibody (guinea pig polyclonal, 1:500, 147 004, Synaptic Systems), an anti-bassoon antibody (mouse monoclonal, 1:500, ADI-VAM-PS003-D, Enzo Life Sciences), an anti-VMAT2 antibody (rabbit polyclonal, 1:1000, VMAT2-Rb-Af720, Nittobo Medical), an anti-GABA_A_Rα1 antibody (rabbit polyclonal, 1:2000, GABAARa1-Rb-Af660, Nittobo Medical), an anti-synaptotagmin 1 antibody (mouse monoclonal, 1:1000, 105 011, Synaptic Systems), an anti-synaptotagmin 7 antibody (rabbit polyclonal, 1:200, 105 173, Synaptic Systems), an anti-synaptotagmin 5/9 antibody (rabbit polyclonal, 1:500, 105 053, Synaptic Systems), an anti-VGAT antibody (rabbit polyclonal, 1:1000, 131 003, Synaptic Systems), an anti-VGAT antibody (guinea pig polyclonal, 1:1000, 131 004, Synaptic Systems), an anti-Dopamine D2 receptor antibody (rabbit polyclonal, 1:500, AB5084P, Sigma-Aldrich), an anti-Dopamine D4 receptor antibody (rabbit polyclonal, 1:500, PA5-28756, Thermo Fisher Scientific) in blocking solution at 4°C overnight, followed by secondary antibodies (goat anti-rabbit, goat anti-mouse, goat anti-chicken, goat anti-guinea pig, 1:1000, Invitrogen) conjugated to Alexa 405, Alexa 488, Alexa 594, and Alexa 647 fluorophores in blocking solution at room temperature for 2 hours. After washing with PBST and PBS, brain slices were mounted onto slides using a mounting medium (P36934 or P36935, Invitrogen).

For quantitative analysis, fluorescence images were captured by an FV1000 confocal laser scanning microscope (Olympus) using a 40x/0.95 NA water immersion objective (zoom factor: 1.5, image size: 512 x 512 pixels), an LSM780N multi-photon confocal laser scanning microscope with airyscan (Carl Zeiss) using a 63x/1.46 NA oil immersion objective (zoom factor: 3, image size: 1248 x 1248 pixels), and an LSM880 multi-photon confocal laser scanning microscope with airyscan (Carl Zeiss) using a 63x/1.4 NA oil immersion objective (zoom factor: 3, image size: 5827 x 5827 pixels for stitched images or 1248 x 1248 pixels). Imaging areas were randomly selected and then images were captured from each target region. Images were further analyzed by Zen software (Carl Zeiss), ImageJ program (NIH, measure function), and MATLAB (MathWorks, custom codes). Confocal 3D images were acquired using the z-stack function of LSM880 confocal laser scanning microscope with Airyscan. Images were collected at 0.185 μm intervals and 3.6993 μm depth with a 63x oil-immersion objective, 3x optical zoom, and a resolution of 1248 x 1248 pixels. 3D images were reconstructed and further analyzed by IMARIS 9.6 software.

### Brain slice preparation for electrophysiology

Sagittal brain slices containing the GPe and striatum (250 μm thick) were prepared for whole-cell patch clamp recording. Mice were anesthetized with isoflurane (Piramal Critical Care), decapitated, and the brain was briefly exposed to cold ACSF containing 125 mM NaCl, 2.5 mM KCl, 1.25 mM NaH_2_PO_4_, 25 mM NaHCO_3_, 1 mM MgCl_2_, 2mM CaCl_2_ and 15 mM glucose oxygenated with 95% O_2_ and 5% CO_2_. Acute brain slices were obtained using a tissue vibratome (Leica, VT 1200S) in ice-cold ACSF. Brain slices were first maintained in ACSF for 20 min at 34°C and then another 30 min at room temperature. After recovery, slices were transferred to a submerged recording chamber perfused with ACSF at a rate of 2 – 3 ml/min at 30 – 31°C. Brain slices were used for electrophysiological recordings within 5 hours following slice preparation.

### Electrophysiology, optogenetic stimulation, and pharmacology

GPe neurons were visually identified by conventional IR-DIC optics (BX51WI, Olympus). Whole-cell voltage clamp recordings were made with borosilicate glass pipettes (3.5 – 5.5 MΩ) filled with Cs+ based low Cl- internal solution containing 135 mM CsMeSO3, 10 mM HEPES, 1 mM EGTA, 3.3 mM QX-314, 0.1 mM CaCl2, 4 mM Mg-ATP, 0.3 mM Na3-GTP, 8 mM Na2-phosphocreatine (290 – 300 mOsm, pH 7.3 with CsOH). To measure inhibitory synaptic currents from the GPe neurons, the membrane potential was held at +0 mV (reversal potential of ionotropic glutamate receptors) under the voltage clamp mode with liquid junction potential correction. Access resistance was maintained between 10 – 20 MΩ, and only cells with less than a 20% change in access resistance were included in the analysis. Whole-cell patch clamp recordings were performed using Multiclamp 700B (Molecular Devices) and signals were filtered at 2 kHz and digitized at 10 kHz (NI PCIe-6259, National Instruments). Recording data were monitored and acquired by WinWCP (Strathclyde software, http://spider.science.strath.ac.uk/sipbs/software_ses.htm), further analyzed offline using Clampfit 10.7 software (Molecular Devices) and OriginPro 2017 (OriginLab). To stimulate ChR2- expressing striatopallidal axons, blue laser light (450 nm, 0.2 ms pulses with 40 sec intervals, 50% of saturation power under the objective less than 1 mW) from diode laser (MDL-III-450, Opto Engine LLC) was focused on the back focal plane of the objective to generate wide-field illumination.

To test the physiological effects of D2-like receptors on striatopallidal GABAergic transmission, we used the following agonists and antagonists: quinpirole, a DA D2-like receptors agonist ((-)-Quinpirole hydrochloride, 10 μM, Tocris, 1061), A-412997, a D4R-selective agonist (A 412997 dihydrochloride, 50 nM, 100 nM, Tocris, 4552), and L-741626, a D2R-selective antagonist (L-741,626, 100nM, Tocris, 1003). These drugs were bath-applied during the recording session after stable baseline IPSC traces were established. To measure the paired pulse ratio (PPR) and ongoing activity of GABA transmission at striatopallidal synapses in the GPe subregions, we optogenetically stimulated striatopallidal axon terminals by using paired light stimuli (50 ms interval) for PPR assessment and trains of 10 light pulses at 20 Hz for ongoing activity, respectively. For the experiment measuring strontium-induced asynchronous GABA release from striatopallidal synaptic terminals, strontium chloride hexahydrate (4 mM, Sigma-Aldrich, Cat. No. 255521) was added to the ACSF in place of calcium. After recording oIPSCs in normal ACSF, the calcium-containing ACSF was replaced with strontium-containing ACSF. The amplitude and frequency of strontium-induced asynchronous IPSCs were analyzed using Mini Analysis software (Synaptosoft). To quantify the relative suppression level of ongoing activity from individual IPSCs, the charge ratio was calculated using the following equation^31^.

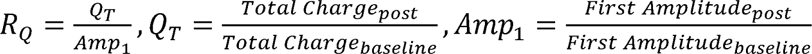

### AI-based strontium mini clustering analysis

Strontium mini responses were analyzed using Mini Analysis software (Synaptosoft). Three parameters—amplitude, rise time, and decay time—were extracted from each mini response for clustering analysis. Prior to clustering, all mini response data from cells across the four regions (DL, VL, DM, and VM) were combined into a single dataset. Two methods were employed for clustering analysis: dimensionality reduction and unsupervised clustering. For dimensionality reduction, principal component analysis (PCA) was applied to reduce noise and explain variance in the three-dimensional data (MATLAB). Subsequently, K-means clustering was employed as the unsupervised clustering method, with the number of clusters set to 50 (MATLAB). The number of mini responses in each cluster was compared across all possible combinations of regions using unpaired Student’s t-tests. If more than one cluster exhibited statistically significant differences in mini response counts, each region was considered to have different characteristics of mini responses. Clusters containing fewer than 1% of the total mini responses were excluded from the analysis to avoid the influence of outliers. This entire process was repeated 1000 times, and the probability of observing significant differences across all regional combinations was calculated.

### Biocytin labeling and immunostaining of biocytin-filled neurons

For examining the cell type of the recorded GPe neurons, Cs-based internal solution containing 0.2% biocytin (B4261, Sigma-Aldrich) was loaded into GPe neurons for 20 min. After the delicate retraction of the glass pipette, brain slices were fixed in 4% PFA overnight at 4°C. On the following day, slices were washed with PBS, PBST for 1 hours and blocked with solution containing 10% NGS and 2% BSA for 4 hours. After blocking, brain sections were incubated with an anti-Parvalbumin antibody (guinea pig polyclonal, 1:1000, 195 004, Synaptic Systems) and an anti-FOXP2 antibody (rabbit polyclonal, 1:500, ab16046, Abcam), in blocking solution at 4°C for 3 days, followed by secondary antibodies (goat anti-rabbit, goat anti-guinea pig, 1:1000, Invitrogen) conjugated to Alexa 594 and Alexa 647 fluorophores and Alexa 405-conjugated streptavidin (1:500, Invitrogen) in blocking solution at 4°C for 2 days. After washing with PBST and PBS for 12 hours, brain slices were mounted onto slides using a mounting medium (P36934, Invitrogen). Images were obtained by an FV1000 confocal laser scanning microscope (Olympus) and an LSM880 multi-photon confocal laser scanning microscope with airyscan (Carl Zeiss). Cell type clarification was conducted using ImageJ.

### Virus purchase

AAV9-hSyn-rDA2h was purchased from WZ Biosciences, with permission from Dr. Yulong Li (Peking University School of Life Sciences). The production titer was 3.28 × 10^13^ virus molecules/ml.

### AAV vector production

For adeno-associated virus (AAV)-based synaptophysin expression, we purchased pAAV-hSyn- FLEx-mGFP-2A-Synaptophysin-mRuby (Addgene, #71760), which allows expression of mGFP and Synaptophysin-mRuby in Cre-expressing neurons, and pAAV8-CAG-FRT-Rev-3xGFP (Addgene, #191204), a FlpO-dependent GFP expression vector. AAVs were thereafter purified by iodixanol gradient ultracentrifugation by the KIST Virus Facility. The production titers of virus were as follows: pAAV-hSyn-FLEx-mGFP-2A-Synaptophysin-mRuby (1.19 × 10^13^ genome copy/ml), pAAV8-CAG-FRT-Rev-3xGFP (1.86 × 10^13^ genome copy/ml).

### Stereotaxic viral injection for striatopallidal axon labeling

Stereotaxic virus injections for striatopallidal axon labeling were conducted in Adora2A-Cre mice. Mice were deeply anesthetized by intraperitoneal injection of zoletil (60 mg/kg, Virbac Korea) and rompun (15 mg/kg, Bayer Korea) mixture solution (zoletil : rompun : saline = 4 : 1 : 20) and mounted in a stereotaxic frame (51730, Stoelting). A small hole was drilled after exposing the skull, and AAVs were injected through a glass micropipette with a long, narrow tip (size: 10 – 20 μm), which was fabricated using a micropipette puller (P-1000, Sutter Instrument). The glass pipette was slowly advanced to the coordinates specified for each target area and left in place for 5 min prior to virus injection. The virus solution was injected at an infusion rate of 100 nl/min, and the glass pipette was withdrawn 10 min after the completion of the injection. Following the injection, the scalp was sutured, and the mice were returned to their home cages for a minimum of 21 days before the subsequent experiments. A total volume of 1600 nl virus solution (400 nl per injection site, AAV5-hSyn-Flex-mGFP-2A-Synaptophysin-mRuby) was injected into the striatum (coordinates used, AP: +0.8 mm, ML: ±2.65 mm from the bregma, DV: -4.2 mm and -2.5 mm for the lateral striatum from the exposed dura mater, AP: +0.85 mm, ML: ±2.00 mm from the bregma, DV: -3.2 mm and -1.85 mm for the medial striatum from the exposed dura mater) to achieve sufficient viral expression across the entire population of striatal neurons. Mice injected with the virus were subsequently used for immunohistochemistry 6 weeks post-injection.

### Stereotaxic 6-OHDA injection

Stereotaxic 6-OHDA injections were conducted on mice (Adora2A-Cre, Adora2A-Cre;Ai32, and Adora2A-Cre;Ai9) using a stereotaxic system (51730, Stoelting). Before surgery, mice were deeply anesthetized by intraperitoneal injection of zoletil (60 mg/kg, Virbac Korea) and rompun (15 mg/kg, Bayer Korea) mixture solution (zoletil : rompun : saline = 4 : 1 : 20). A total volume of 300 nl 6-OHDA solution (2.5 mg/ml, dissolved in 0.9% sterile saline with 0.02% ascorbic acid) was injected unilaterally into the left MFB (coordinates used, AP: -0.7 mm, ML: +1.2 mm from the bregma, DV: -4.80 mm from the exposed dura mater for mice aged 3 to 4 weeks, AP: -1.2 mm, ML: +1.2 mm from the bregma, DV: -4.75 mm from the exposed dura mater for mice older than 8 weeks). A glass micropipette with a long, narrow tip (size: 10 – 20 μm) was made using a micropipette puller (P-1000, Sutter Instrument) to deliver 6-OHDA. The glass pipette was slowly advanced to the target area and left in place for 5 min prior to the 6-OHDA injection. 6-OHDA solution was injected at an infusion rate of 100 nl/min, and the glass pipette was withdrawn 10 min after the end of injection. After the injection, the scalp was sutured, and the mice were returned to their home cages. Mice injected with 6-OHDA were used for electrophysiology and immunohistochemistry experiments at 1, 3, 5, and 7 days post-injection.

### Spot detection and synapse extraction analysis

Spot detection and synapse extraction analysis were performed similarly as previously described^20,63,64^. To extract synapses from the acquired images, we implemented a general method for spot detection. The method utilizes mathematical morphological processing based on set theory. Signal enhancement and spot detection can be achieved by treating objects within images as sets and utilizing combinations of logical operators in set theory. This method consists of three steps: denoising, signal enhancement, and spot extraction. In the denoising step, the acquired images are processed using Gaussian filter with σ = 0.5. For signal enhancement, a combination of three thresholding algorithms with top-hat transform, hard thresholding, and Otsu thresholding are applied to the resulting images. Top-hat transform of an image *I*(*m*) is represented as:

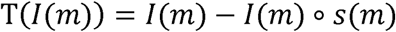

where ° denotes the opening operator in morphological processing and *s*(*m*) denotes the structuring element. Our structuring element *s*(*m*) for applying top-hat transform is circular shape with size of 10 X10. Hard thresholding is expressed as:

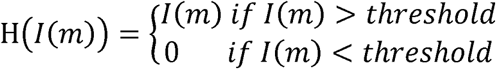

Threshold value of hard thresholding is 45000. After applying top-hat transform and hard thresholding to the acquired images, Otsu’s thresholding method is exploited to the processed images to enhance fluorescent signal from the actual objects while suppressing background noise. From the signal-enhanced images, spots are extracted using a combination of morphological filters, which include two key operations: filling and opening. The filling is used to fill out holes in fluorescent spots, while the opening operation smooths spot contours and removes irrelevant fluorescent signals unrelated to synaptic structures. The extracted spots are clustered within the set of images (presynaptic and postsynaptic) corresponding to distance among the spots, which allows the images to delineate presynaptic and postsynaptic structures. By calculating the distance between presynaptic and postsynaptic clusters, those with a separation of less than 1 pixel (40 nm) were merged to form synapses.

### Point pattern analysis

The spatial analysis of clustered features between striatopallidal axon terminals and D2Rs was conducted using Ripley’s analysis on both single group and group vs. group scale^65^. Ripley’s function describes spatial characteristics of the sample (clustering or dispersion) over a range of distances. The edge corrected Ripley’s function is defined as:

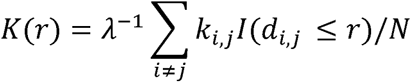

where λ is the density of a point pattern; *k_i,j_* is the inverse of a portion of circumference with center at point , and radius i, that lies within the boundary and passes through point ,; function I is an indicator function; d is the Euclidean distance between points i and j; N is the number of points in the pattern. A variance stabilized version of the Ripley’s K function was used in this study:

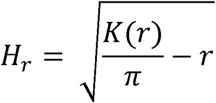

Ripley’s function was calculated for each observed point pattern. Group vs. group analysis was also used to determine whether point patterns from one group had similar spatial properties with patterns from another (null) group. We assume that all m (the total number of generated patterns) patterns in the null group are generated using the identical point method.

### Diggle-Cressie-Loosmore-Ford (DCLF) test

Diggle-Cressie-Loosmore-Ford (DCLF) test was performed similarly as previously described20,66. The DCLF test quantifies the difference between the H function of the observed pattern and the H function for the null group at the given spatial scale. We computed *H*(*r*) for the test pattern *H*(*r*) and each pattern in the null group *H_obs_*(*r*), then we estimated the summary function *H*(*r*) for the null group as follows:

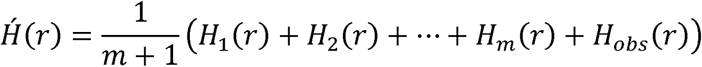

Then, the maximum vertical separation between *H*(*r*) and *H*(*r*) is:

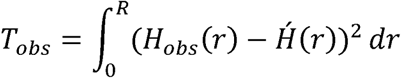

where R is maximum value of the interaction distance. To compute p-value, the following formula can be used: 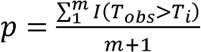, where 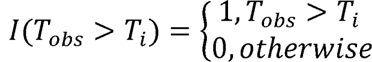 and *T*_*i*_ computed between the observed pattern and each pattern in the test group 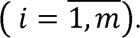.

### Computational model of synaptic transmission

A computational model is based on the Tsodyks-Markram model^42^ to simulate synaptic responses and to understand the underlying mechanisms affecting neurotransmitter release probability and synaptic plasticity under different experimental conditions. The model incorporated both facilitation and depression dynamics, adapting established frameworks for synaptic transmission modeling. The following list outlines the key variables and parameters utilized in this modeling approach.

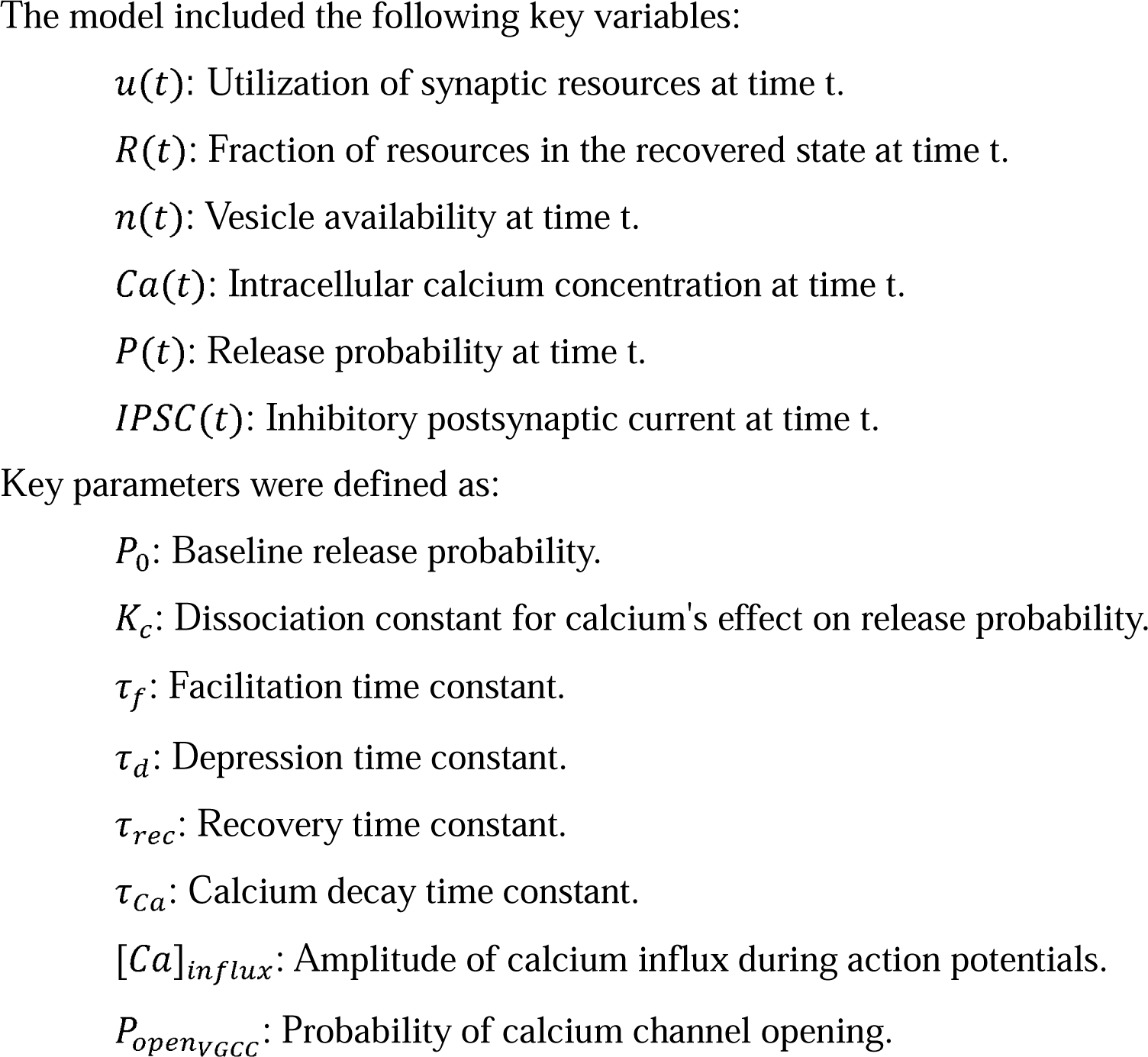

### Mathematical framework

The model was governed by a set of differential equations:

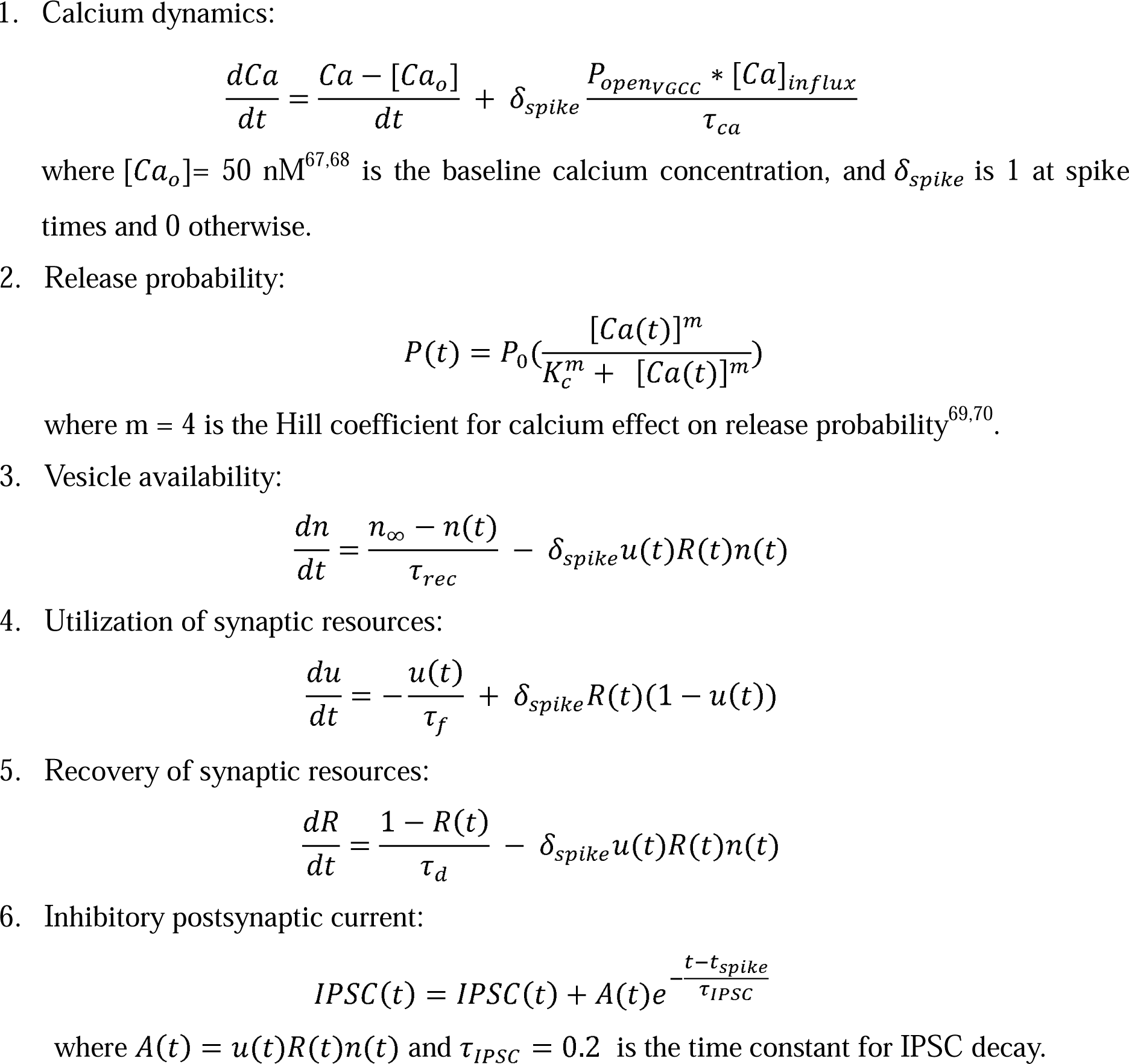

### Simulation protocol

Simulations were conducted over a 1-second duration with a time step of dt = 0.001 seconds. Action potentials were simulated at 0.5 and 0.55 seconds to represent paired-pulse stimulation. The differential equations were numerically integrated using the Euler method. Initial conditions were set based on physiological values, and variables were constrained to remain within plausible biological ranges to prevent computational errors.

### Parameter optimization

An optimization routine was employed to fit the model to the experimental PPR data. The objective function minimized the squared difference between the simulated PPR and the observed PPR for each condition. The optimization process involved:

1. Control condition: Parameters were initially optimized for the control condition using the differential evolution. To ensure physiological plausibility, bounds were specified for each parameter, and the number of iterations was capped at 100. To accurately capture the synaptic behavior observed in different conditions, the bounds for the baseline release probability (P0) were dynamically adjusted based on the observed paired-pulse ratio (PPR). Specifically, for the facilitating synapses (PPR > 1.05) the bounds for P0 were set to (0.1, 0.5). This range allows P0 to vary within values that support an increasing release probability upon successive stimuli, characteristic of facilitating synapses. For synapses that are non-facilitating or depressing (PPR ≈ 1 or PPR < 1) the bounds for P0 were adjusted to (0.55, 0.9).
2. Quinpirole condition: Starting from the optimized control parameters, scaling factors the VGCC open probability (*P_openvGCC_*). The optimization aimed to reproduce the were introduced to simulate the effect of quinpirole, a dopamine D2 receptor agonist on for the *P_openvGCC_* was optimized, while the rest of the parameters remained the same as observed changes in PPR under quinpirole treatment. In this case the scaling parameter in the optimized ‘control conditions’. If the synapses changed from non-facilitating (PPR ≈ 1) to facilitating (PPR > 1) under ‘quinpirole condition’, the following parameters were manually adjusted to guarantee facilitation.
3. Summary of parameter bounds 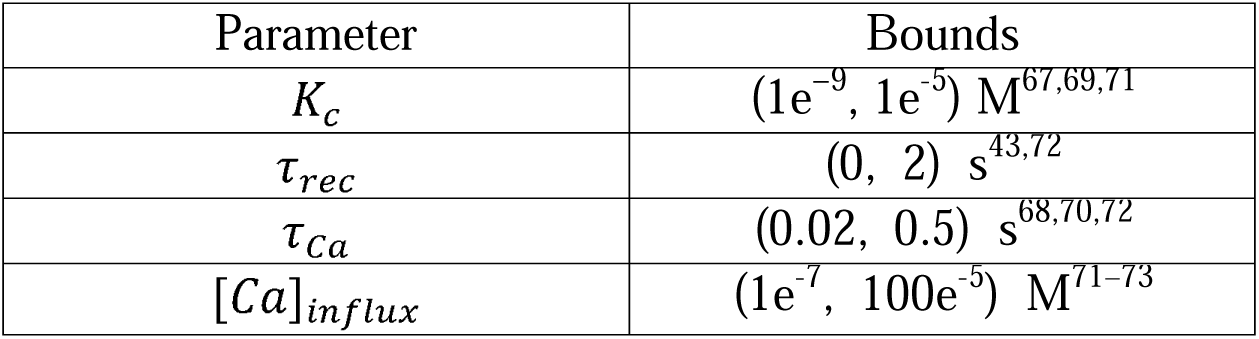

### Statistical analysis and exclusion criteria for computational modeling

Statistical comparisons between conditions were performed using non-parametric tests due to the sample size and data distribution. The Wilcoxon Signed-Rank Test was applied for paired comparisons of parameters, and the Mann-Whitney Test was applied for unpaired comparisons within the same cells under different conditions. Cells were excluded from analysis if the simulated PPR under quinpirole did not match the observed behavior (e.g., the model predicted facilitation when depression was observed experimentally, or vice versa). Additionally, cells with discrepancies exceeding a threshold (e.g., absolute difference in PPR greater than 0.4) were omitted to ensure the reliability of the model fitting.

### Software and libraries

All computations were performed using Python 3.9, with key libraries including: NumPy: For numerical computations and array manipulations.

SciPy: For optimization routines and statistical tests. Pandas: For data handling and manipulation.

### Statistics

All data were analyzed using the following software: Clampfit 10.7 (Molecular Devices), Mini Analysis (version 6.0.7, Synaptosoft), OriginPro 2017 (OriginLab), Zen software (version 2.5, Carl Zeiss), ImageJ (version 1.53c, NIH), and MATLAB (version R2021a, MathWorks). Statistical analyses were performed using GraphPad Prism (version 10, GraphPad software). Summary statistics were presented as either box-and-whisker plots or mean ± SEM. Unpaired and paired student t-tests, one-way ANOVA with Holm-Sidak’s post-hoc multiple comparison test, two-way repeated measures ANOVA with Holm-Sidak’s post-hoc multiple comparison test, linear regression, and the Kolmogorov-Smirnov test were employed to assess statistical differences (two-sided) between the groups. A P-value of less than 0.05 was considered statistically significant. *p < 0.05, **p < 0.01, ***p < 0.001, ****p < 0.0001.

### Data availability

All data generated and analyzed in the current study are available from the corresponding author upon reasonable request.

### Code availability

Python and MATLAB codes used in data analysis are available from the corresponding author upon reasonable request.

## Acknowledgements

The authors express their gratitude to Dr. Hyoung-Ro Lee and Dr. Suk-Ho Lee for their invaluable assistance with axon terminal Ca^2+^ imaging and to the members of Kim laboratory for their insightful discussions. This research was supported by Basic Science Research Program through the National Research Foundation of Korea (NRF) funded by the Ministry of Science and ICT (2021R1A4A1031644, 2021M3A9G8022960, 2022M325E8017907, 2023R1A2C1006489 to JIK), intramural research fund of UNIST (1.210115.01 to JIK), and the Institute for Basic Science (IBS-R022-D1 to KM). Figures 4b,g, Figures 8a, and Extended Data Figures 3a and 10b were created with BioRender.com.

## Author contributions

Conceptualization: YL, JIK.

Methodology: YL, MR, KJK, HJK, MSJ, KM, YL, SEL, EJK, CJL, JIK

Investigation: YL, MR, KJK, YK, EC, JIK

Interpretation of Data: YL, MR, KJK, EJK, CJL, CL, JIK

Writing - Original Draft: YL, JIK

Writing - Review and Editing: YL, CL, JIK

Funding Acquisition: KM, JIK

Supervision: JIK

## References

1. Kravitz, A. V. et al. Regulation of parkinsonian motor behaviours by optogenetic control of basal ganglia circuitry. Nature 466, 622–626 (2010).

2. Arber, S. & Costa, R. M. Connecting neuronal circuits for movement. Science *(80-. ).* **360**, 1403–1404 (2018).

3. Roth, R. H. & Ding, J. B. Cortico-basal ganglia plasticity in motor learning. Neuron 112, 2486–2502 (2024).

4. Gerfen, C. R. Molecular effects of dopamine on striatal-projection pathways. Trends Neurosci. 23, 64–70 (2000).

5. Kreitzer, A. C. & Malenka, R. C. Striatal Plasticity and Basal Ganglia Circuit Function. Neuron 60, 543–554 (2008).

6. Hedreen, J. C. Tyrosine hydroxylase-immunoreactive elements in the human globus pallidus and subthalamic nucleus. J. Comp. Neurol. 409, 400–410 (1999).

7. Smith, Y. & Kieval, J. Z. Anatomy of the dopamine system in the basal ganglia. Trends Neurosci. 23, 28–33 (2000).

8. Meszaros, J. et al. Evoked transients of ph-sensitive fluorescent false neurotransmitter reveal dopamine hot spots in the globus pallidus. Elife 7, 1–19 (2018).

9. Lindvall, O. & Björklund, A. Dopaminergic innervation of the globus pallidus by collaterals from the nigrostriatal pathway. Brain Res. 172, 169–173 (1979).

10. Mamad, O., Delaville, C., Benjelloun, W. & Benazzouz, A. Dopaminergic control of the globus pallidus through activation of D2 receptors and its impact on the electrical activity of subthalamic nucleus and substantia nigra reticulata neurons. PLoS One 10, 1–16 (2015).

11. Marjorie A. Ariano, Jean Wang, Kurtis L. Noblett, Eric R.Larson, D. R. S. Cellular distribution of the rat D4 dopamine receptor protein in t anti-receptor antisera. Brain Res. 752, 26–34 (1997).

12. Mauger, C. et al. Development and characterization of antibodies directed against the mouse D4 dopamine receptor. Eur. J. Neurosci. 10, 529–537 (1998).

13. Foster, N. N. et al. The mouse cortico–basal ganglia–thalamic network. Nature 598, 188– 194 (2021).

14. Lee, J., Wang, W. & Sabatini, B. L. Anatomically segregated basal ganglia pathways allow parallel behavioral modulation. Nat. Neurosci. 23, 1388–1398 (2020).

15. Smith, Y., Lavoie, B., Dumas, J. & Parent, A. Evidence for a distinct nigropallidal dopaminergic projection in the squirrel monkey. Brain Res. 482, 381–386 (1989).

16. Fuchs, H. & Hauber, W. Dopaminergic innervation of the rat globus pallidus characterized by microdialysis and immunohistochemistry. Exp. Brain Res. 154, 66–75 (2004).

17. Patriarchi, T. et al. Ultrafast neuronal imaging of dopamine dynamics with designed genetically encoded sensors. Science *(80-. ).* **360**, (2018).

18. Sun, F. et al. A Genetically Encoded Fluorescent Sensor Enables Rapid and Specific Detection of Dopamine in Flies, Fish, and Mice. Cell 174, 481–496.e19 (2018).

19. Sun, F. et al. Next-generation GRAB sensors for monitoring dopaminergic activity in vivo. Nat. Methods 17, 1156–1166 (2020).

20. Kim, H. J. et al. GABAergic-like dopamine synapses in the brain. Cell Rep. 42, 113239 (2023).

21. Cooper, A. J. & Stanford, I. M. Dopamine D2 receptor mediated presynaptic inhibition of striatopallidal GABAA IPSCs in vitro. Neuropharmacology 41, 62–71 (2001).

22. Watanabe, K., Kita, T. & Kita, H. Presynaptic actions of D2-like receptors in the rat cortico-striato-globus pallidus disynaptic connection in vitro. J. Neurophysiol. 101, 665– 671 (2009).

23. Zych, S. M. & Ford, C. P. Divergent properties and independent regulation of striatal dopamine and GABA co-transmission. Cell Rep. 39, 110823 (2022).

24. Yamamoto, K. & Kobayashi, M. Opposite roles in short-term plasticity for N-type and P/Q-type voltage-dependent calcium channels in gabaergic neuronal connections in the rat cerebral cortex. J. Neurosci. 38, 9814–9828 (2018).

25. Sims, R. E., Woodhall, G. L., Wilson, C. L. & Stanford, I. M. Functional characterization of GABAergic pallidopallidal and striatopallidal synapses in the rat globus pallidus in vitro. Eur. J. Neurosci. 28, 2401–2408 (2008).

26. Perez-Rosello, T. et al. Enhanced striatopallidal gamma-aminobutyric acid (GABA)A receptor transmission in mouse models of huntington’s disease. Mov. Disord. 34, 684–696 (2019).

27. Bevan, M. D. Selective innervation of neostriatal interneurons by a subclass of neuron in the globus pallidus of the rat. J. Neurosci. 18, 9438–9452 (1998).

28. Liu, C., Kershberg, L., Wang, J., Schneeberger, S. & Kaeser Correspondence, P. S. Dopamine Secretion Is Mediated by Sparse Active Zone-like Release Sites In Brief Secretion of dopamine requires specialized release machinery. Cell 172, 706–709.e15 (2018).

29. Kershberg, L., Banerjee, A. & Kaeser, P. S. Protein composition of axonal dopamine release sites in the striatum. Elife 11, 1–29 (2022).

30. Shin, R. M. et al. Dopamine D4 Receptor-Induced Postsynaptic Inhibition of GABAergic Currents in Mouse Globus Pallidus Neurons. J. Neurosci. 23, 11662–11672 (2003).

31. Burke, K. J., Keeshen, C. M. & Bender, K. J. Two Forms of Synaptic Depression Produced by Differential Neuromodulation of Presynaptic Calcium Channels. Neuron 99, 969–984.e7 (2018).

32. Kaeser, P. S. & Regehr, W. G. The readily releasable pool of synaptic vesicles. Curr. Opin. Neurobiol. 43, 63–70 (2017).

33. Katayama, S. et al. Slowly progressive L-DOPA nonresponsive pure akinesia due to nigropallidal degeneration: A clinicopathological case study. J. Neurol. Sci. 161, 169–172 (1998).

34. Tan, W. Q. et al. Deterministic Tractography of the Nigrostriatal-Nigropallidal Pathway in Parkinson’s Disease. Sci. Rep. 5, 2–7 (2015).

35. Parent, A., Lavoie, B., Smith, Y. & Bédard, P. The dopaminergic nigropallidal projection in primates: distinct cellular origin and relative sparing in MPTP-treated monkeys. Adv. Neurol. 53, 111–116 (1990).

36. Jones, J. A., Higgs, M. H., Olivares, E., Peña, J. & Wilson, C. J. Spontaneous Activity of the Local GABAergic Synaptic Network Causes Irregular Neuronal Firing in the External Globus Pallidus. J. Neurosci. 43, 1281–1297 (2023).

37. Mallet, N., Delgado, L., Chazalon, M., Miguelez, C. & Baufreton, J. Cellular and synaptic dysfunctions in Parkinson’s disease: Stepping out of the striatum. Cells 8, 1–29 (2019).

38. Galvan, A. & Wichmann, T. Pathophysiology of Parkinsonism. Clin. Neurophysiol. 119, 1459–1474 (2008).

39. Kiskowski, M. A., Hancock, J. F. & Kenworthy, A. K. On the use of Ripley’s K-function and its derivatives to analyze domain size. Biophys. J. 97, 1095–1103 (2009).

40. Rebola, N. et al. Distinct Nanoscale Calcium Channel and Synaptic Vesicle Topographies Contribute to the Diversity of Synaptic Function. Neuron 1–18 (2019). doi:10.1016/j.neuron.2019.08.014

41. N Bert Loosmore, E. D. F. Statistical inference using the g or K point pattern spatial statistics. Ecology 87, 1925–1931 (2006).

42. Tsodyks, M. V. & Markram, H. The neural code between neocortical pyramidal neurons depends on neurotransmitter release probability. Proc. Natl. Acad. Sci. U. S. A. 94, 719– 723 (1997).

43. Markram, H., Wang, Y. & Tsodyks, M. Differential signaling via the same axon of neocortical pyramidal neurons. Proc. Natl. Acad. Sci. U. S. A. 95, 5323–5328 (1998).

44. Kisilevsky, A. E. & Zamponi, G. W. D2 dopamine receptors interact directly with N-type calcium channels and regulate channel surface expression levels. Channels 2, 269–277 (2008).

45. Scarnati, M. S., Clarke, S. G., Pang, Z. P. & Paradiso, K. G. Presynaptic Calcium Channel Open Probability and Changes in Calcium Influx Throughout the Action Potential Determined Using AP-Waveforms. Front. Synaptic Neurosci. 12, 1–18 (2020).

46. Mastro, K. J., Bouchard, R. S., Holt, H. A. K. & Gittis, A. H. Transgenic mouse lines subdivide external segment of the globus pallidus (GPe) neurons and reveal distinct GPe output pathways. J. Neurosci. 34, 2087–2099 (2014).

47. Beier, K. T. et al. Rabies screen reveals GPe control of cocaine-triggered plasticity. Nature 549, 345–350 (2017).

48. Dodson, P. D. et al. Distinct developmental origins manifest in the specialized encoding of movement by adult neurons of the external globus pallidus. Neuron 86, 501–513 (2015).

49. Mastro, K. J. et al. Cell-specific pallidal intervention induces long-lasting motor recovery in dopamine-depleted mice. Nat. Neurosci. 20, 815–823 (2017).

50. Jeon, H. et al. Topographic connectivity and cellular profiling reveal detailed input pathways and functionally distinct cell types in the subthalamic nucleus. Cell Rep. 38, 110439 (2022).

51. Lindgren, N. et al. Distinct roles of dopamine D2L and D2S receptor isoforms in the regulation of protein phosphorylation at presynaptic and postsynaptic sites. Proc. Natl. Acad. Sci. U. S. A. 100, 4305–4309 (2003).

52. Alessandro Usiello, Ja-Hyun Baik, FrancËoise RougeÂ-Pont, Roberto Picetti, AndreÂe Dierich, Marianne LeMeur, P. V. P. & E. B. Distinct functions of the two isoforms of dopamine D2 receptors. Nature 408, 199–203 (2000).

53. Wang, Y. et al. Heterogeneity in the pyramidal network of the medial prefrontal cortex. Nat. Neurosci. 2006 94 9, 534–542 (2006).

54. Bao, J., Reim, K. & Sakaba, T. Target-Dependent Feedforward Inhibition Mediated by Short-Term Synaptic Plasticity in the Cerebellum. J. Neurosci. 30, 8171–8179 (2010).

55. Afia B. Ali and Alex M. Thomson. Facilitating pyramid to horizontal oriens—alveus interneuroneinputs: dual intracellular recordings in slices of rathippocampus. Journal of physiology 185–199 (1998).

56. Lehéricy, S. et al. Diffusion tensor fiber tracking shows distinct corticostriatal circuits in humans. Ann. Neurol. 55, 522–529 (2004).

57. Lee, J. H., Kim, D. S., Baik, S. K. & Nam, S. O. Nigropallidal iron accumulation in pantothenate kinase-associated neurodegeneration demonstrated by susceptibility- weighted imaging. J. Neurol. 257, 661–662 (2010).

58. Brichta, L. & Greengard, P. Molecular determinants of selective dopaminergic vulnerability in Parkinson’s disease: An update. Front. Neuroanat. 8, 122879 (2014).

59. Carmichael, K. et al. Function and Regulation of ALDH1A1-Positive Nigrostriatal Dopaminergic Neurons in Motor Control and Parkinson’s Disease. Front. Neural Circuits 15, 644776 (2021).

60. Matthias H. Hennig. Theoretical models of synaptic short term plasticity. Comput. Neurosci. 7, (2013).

61. Wei, W. et al. Supersensitive presynaptic dopamine D2 receptor inhibition of the striatopallidal projection in nigrostriatal dopamine-deficient mice. J. Neurophysiol. 110, 2203–2216 (2013).

62. Nambu, A. & Tachibana, Y. Mechanism of parkinsonian neuronal oscillations in the primate basal ganglia: Some considerations based on our recent work. Front. Syst. Neurosci. 8, 67336 (2014).

63. Aishwarya, N., Phamila, Y. A. V. & Amutha, R. Multi-focus image fusion using multi- structure top-hat transform and image variance. Int. Conf. Commun. Signal Process. ICCSP 2013 - Proc. 352–356 (2013). doi:10.1109/iccsp.2013.6577073

64. Kimori, Y., Baba, N. & Morone, N. Extended morphological processing: A practical method for automatic spot detection of biological markers from microscopic images. BMC Bioinformatics 11, (2010).

65. Ripley, B. D. Modelling Spatial Patterns. 39, 172–212 (1977).

66. Baddeley, A. et al. On tests of spatial pattern based on simulation envelopes. Ecol. Monogr. 84, 477–489 (2014).

67. Usowicz, M. M., Sugimori, M., Cherksey, B. & Llinás, R. P-type calcium channels in the somata and dendrites of adult cerebellar purkinje cells. Neuron 9, 1185–1199 (1992).

68. Neher, E. Vesicle pools and Ca2+ microdomains: New tools for understanding their roles in neurotransmitter release. Neuron 20, 389–399 (1998).

69. Dodge, F. A. & Rahamimoff, R. Co operative action of calcium ions in transmitter release at the neuromuscular junction. J. Physiol. 193, 419–432 (1967).

70. Augustine, George J. Milton P. Charlton, S. J. S. & Howard. Calcium action in synaptic transmitter release. Annu. Rev. Neurosci. 10th, 633–693 (1987).

71. Bollmann, J. H., Sakmann, B. & Borst, J. G. G. Calcium sensitivity of glutamate release in a calyx-type terminal. Science *(80-. ).* **289**, 953–957 (2000).

72. Zucker, R. S. & Regehr, W. G. Short-term synaptic plasticity. Annu. Rev. Physiol. 64, 355–405 (2002).

73. Schneggenburger, R. & Erwin Neher. Intracellular calcium dependence of transmitter release rates at a fast central synapse. Nature 406, 889–893 (2000).

