## Supplementary Materials for "Distinct modes of dopamine modulation on striatopallidal synaptic transmission"

**This PDF file includes**

1. Extended data figures and figure legends
2. Supplementary Table S1
3. Supplementary Table S2

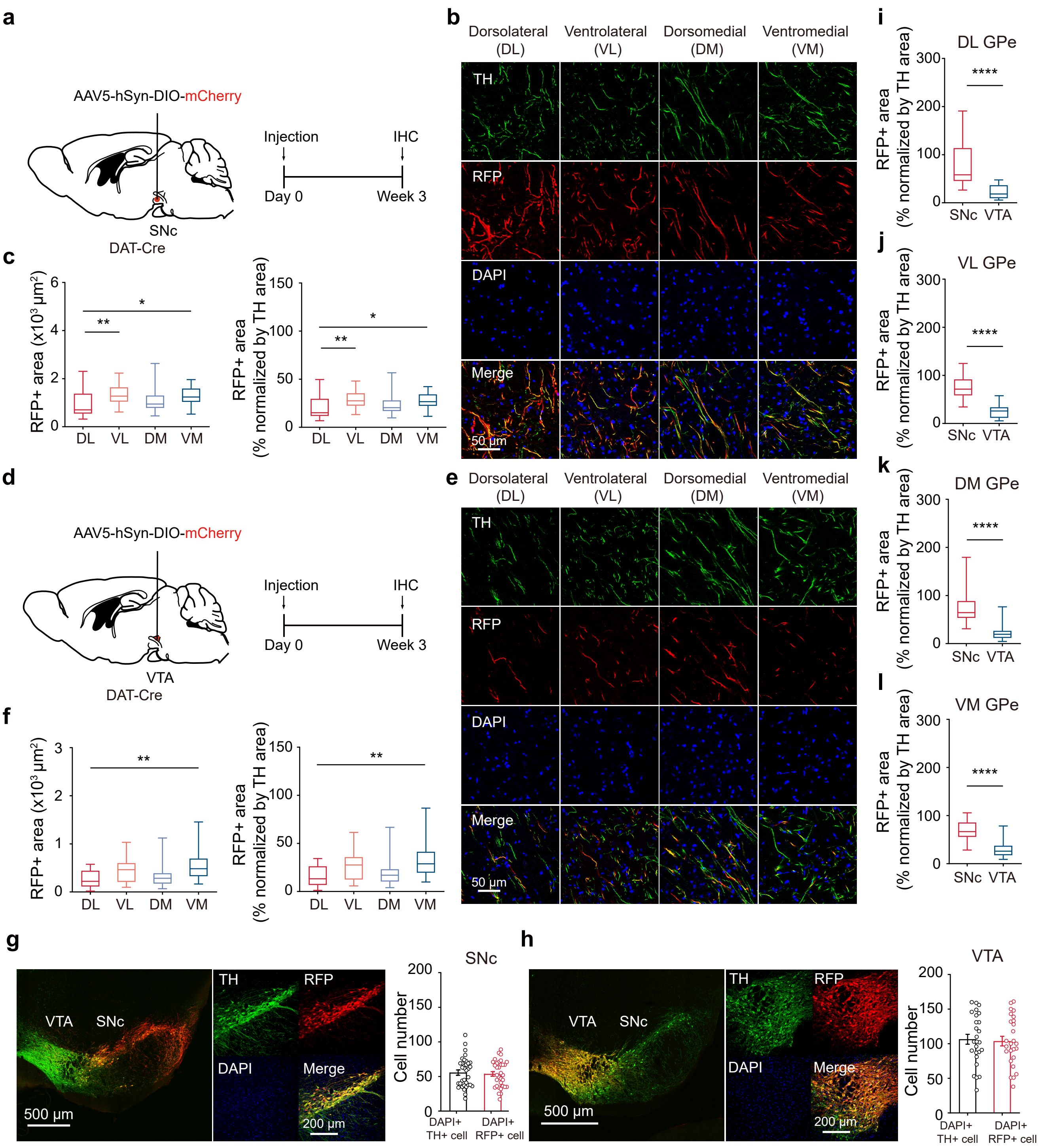
a
AAV5-hSyn-DIO-mCherry
SNc
DAT-Cre
Injection
IHC
Day 0
Week 3
b
Dorsolateral (DL)
Ventrolateral (VL)
Dorsomedial (DM)
Ventromedial (VM)
TH
RFP
DAPI
Merge
50 µm
c
RFP+ area (x10³ µm²)
RFP+ area (% normalized by TH area)
DL VL DM VM
d
AAV5-hSyn-DIO-mCherry
VTA
DAT-Cre
Injection
IHC
Day 0
Week 3
e
Dorsolateral (DL)
Ventrolateral (VL)
Dorsomedial (DM)
Ventromedial (VM)
TH
RFP
DAPI
Merge
50 µm
f
RFP+ area (x10³ µm²)
RFP+ area (% normalized by TH area)
DL VL DM VM
i
DL GPe
j
VL GPe
k
DM GPe
l
VM GPe
RFP+ area (% normalized by TH area)
SNc VTA
g
VTA SNc
500 µm
TH RFP DAPI Merge
200 µm
SNc
Cell number
DAPI+ TH+ cell
DAPI+ RFP+ cell
h
VTA SNc
500 µm
TH RFP DAPI Merge
200 µm
VTA
Cell number
DAPI+ TH+ cell
DAPI+ RFP+ cell

**Extended Data Fig. 1 | The SNc is the principal source of the nigropallidal pathway projecting to the GPe**

**a,d,** A schematic illustrations describing the injection of AAV5-hSyn-DIO-mCherry virus into the SNc and VTA of DAT-Cre mice to selectively label dopaminergic axons from the SNc and VTA. **b,e,** Representative confocal images of TH+ dopaminergic and RFP+ axons originating from the SNc and VTA dopamine neurons. **c,** Summary statistics of virally expressed RFP+ axonal areas in the GPe subregions, originating from the SNc (left) (n = 32 images from 6 mice per each group, one-way ANOVA with Holm-Sidak's post-hoc multiple comparisons test; DL 935.72 ± 89.33 $\mu m^2$, VL 1322.18 ± 70.73 $\mu m^2$, DM 1096.91 ± 92.89 $\mu m^2$, VM 1284.85 ± 62.93 $\mu m^2$; $p = 0.0026$) and RFP+ areas normalized to the TH+ areas (right) (DL 20.17 ± 1.93%, VL 28.50 ± 1.53%, DM 23.64 ± 2.0%, VM 27.69 ± 1.36%). **f,** Summary statistics of virally expressed RFP+ axonal areas in the GPe subregions, originating from the VTA (left) (n = 23 images from 5 mice per each group, one-way ANOVA with Holm-Sidak's post-hoc multiple comparisons test; DL 273.04 ± 37.82 $\mu m^2$, VL 465.54 ± 52.02 $\mu m^2$, DM 369.82 ± 65.53 $\mu m^2$, VM 571.82 ± 67.85 $\mu m^2$; $p = 0.0029$) and RFP+ areas normalized to the TH+ areas (right) (DL 16.25 ± 2.25%, VL 27.71 ± 3.10%, DM 22.01 ± 3.90%, VM 34.03 ± 4.04%). **g,h,** Representative confocal images of TH+ dopamine neurons and RFP+ neurons in the SNc or VTA of DAT-Cre mice (left), and summary statistics of TH+ and RFP+ cell numbers (right) in the SNc (n = 33 images from 6 mice per each group, paired t-test; DAPI+ TH+ cell 55.64 ± 3.79, DAPI+ RFP+ cell 53.73 ± 3.33) or in the VTA (n = 26 images from 5 mice per each group, paired t-test; DAPI+ TH+ cell 106.40 ± 7.16, DAPI+ RFP+ cell 103.70 ± 6.94). **i-l,** Summary statistics of virally expressed RFP+ axonal areas normalized to the TH+ areas in the DL GPe (paired t-test; SNc 77.41 ± 7.39%, VTA 22.59 ± 3.13%), VL GPe (SNc 73.96 ± 3.96%, VTA 26.04 ± 2.91%), DM GPe (SNc 74.79 ± 6.33%, VTA 25.21 ± 4.47%), and VM GPe (SNc 69.20 ± 3.39%, VTA 30.80 ± 3.65%). The data are presented as box-and-whisker plots or mean ± SEM. $^*p < 0.05$, $^{**}p < 0.01$, $^{****}p < 0.0001$.

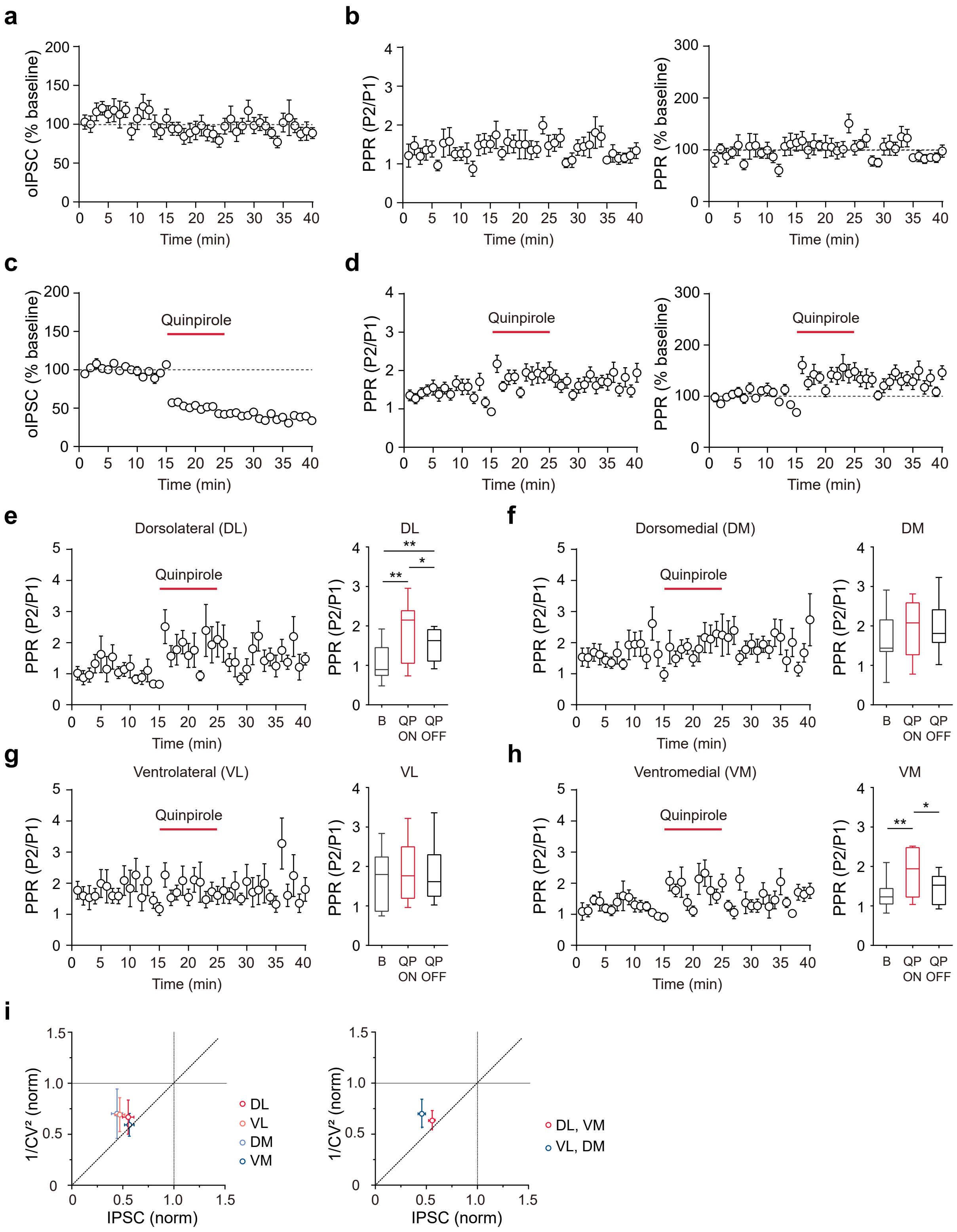
a
b
c
d
e
f
g
h
i
oIPSC (% baseline)
PPR (P2/P1)
PPR (% baseline)
Time (min)
Quinpirole
Dorsolateral (DL)
Dorsomedial (DM)
Ventrolateral (VL)
Ventromedial (VM)
DL
DM
VL
VM
B
QP ON
QP OFF
**
*
1/CV² (norm)
IPSC (norm)
DL
VL
DM
VM
DL, VM
VL, DM

**Extended Data Fig. 2 | Distinct dopaminergic modulation of striatopallidal synaptic transmission through D2-like receptors in the GPe subregions**

**a,b,** Optogenetically evoked striatopallidal synaptic transmission (normalized oIPSC amplitude) and PPR plots. **c,d,** Normalized oIPSC amplitude, PPR plot (left), and normalized PPR plot (right) with the bath application of quinpirole (10 μM). **e-h,** PPR plots and summary statistics of PPR in the DL GPe (repeated measures one-way ANOVA with Holm-Sidak's post-hoc multiple comparisons test; Baseline 1.073 ± 0.147, QP ON 1.867 ± 0.234, QP OFF 1.511 ± 0.126; $p = 0.0014$), DM GPe (Baseline 1.673 ± 0.212, QP ON 1.953 ± 0.217, QP OFF 1.976 ± 0.212; $p = 0.2845$), VL GPe (Baseline 1.717 ± 0.230, QP ON 1.868 ± 0.240, QP OFF 1.801 ± 0.239; $p = 0.6651$), and VM GPe (Baseline 1.285 ± 0.115, QP ON 1.854 ± 0.184, QP OFF 1.470 ± 0.116; $p = 0.0013$). **i,** CV analysis of oIPSCs and $1/CV^2$ after quinpirole treatment, normalized to the baseline in the GPe subregions (left) (oIPSC, DL 0.544 ± 0.046, VL 0.475 ± 0.041, DM 0.453 ± 0.045, VM 0.552 ± 0.039; $1/CV^2$, DL 0.674 ± 0.138, VL 0.694 ± 0.138, DM 0.701 ± 0.201, VM 0.611 ± 0.092), (right) (oIPSC, DL-VM GPe 0.548 ± 0.029, VL-DM GPe 0.464 ± 0.030; $1/CV^2$, DL-VM GPe 0.643 ± 0.081, VL-DM GPe 0.697 ± 0.118). The data are presented as box-and-whisker plots or mean ± SEM. $^*p < 0.05$, $^{**}p < 0.01$.

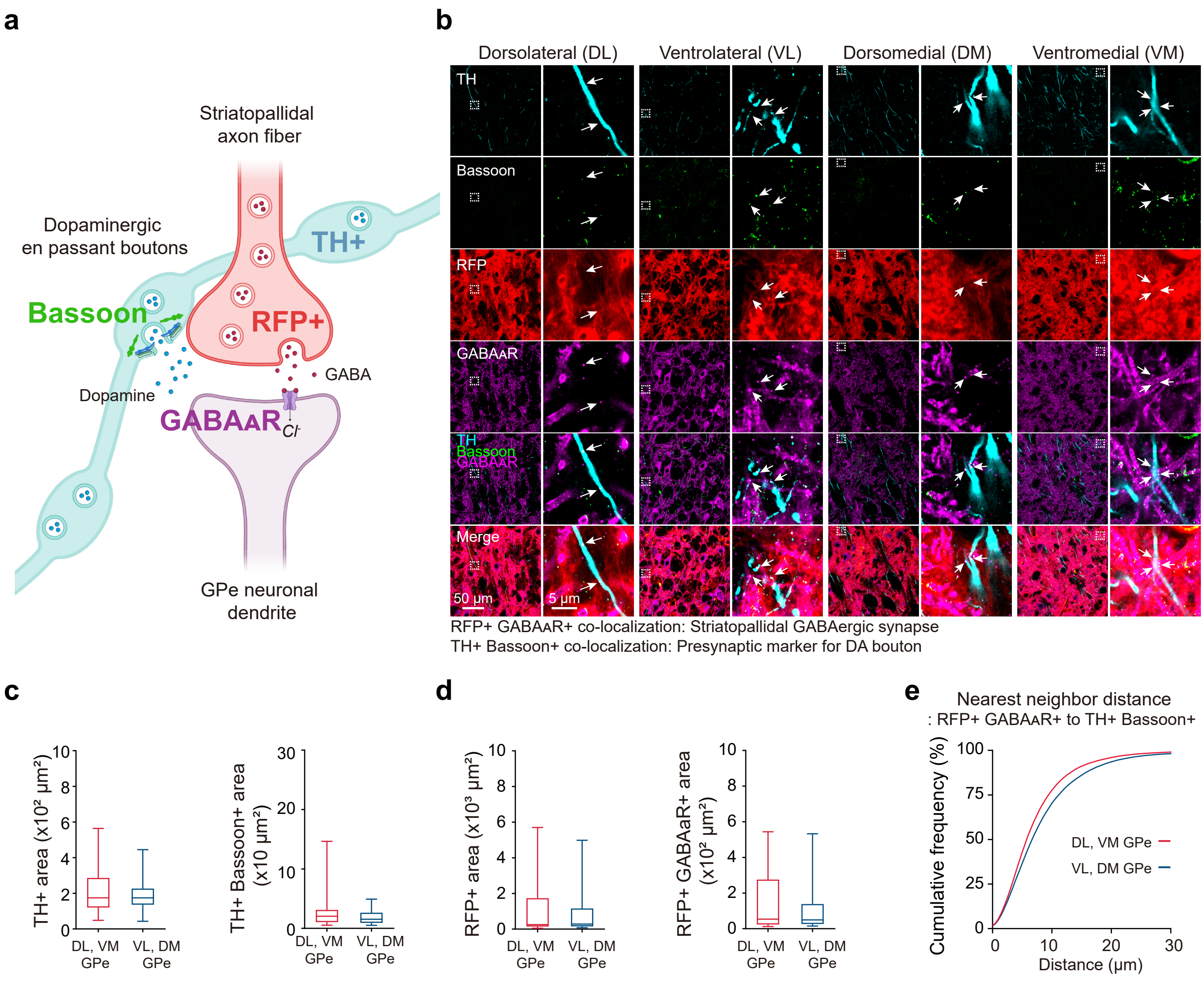
a
Striatopallidal
axon fiber
Dopaminergic
en passant boutons
TH+
Bassoon
RFP+
GABA
Dopamine
GABAAR
Cl-
GPe neuronal
dendrite
b
Dorsolateral (DL)
Ventrolateral (VL)
Dorsomedial (DM)
Ventromedial (VM)
TH
Bassoon
RFP
GABAAR
TH
Bassoon
GABAAR
Merge
50 µm
5 µm
RFP+ GABAAR+ co-localization: Striatopallidal GABAergic synapse
TH+ Bassoon+ co-localization: Presynaptic marker for DA bouton
c
TH+ area (x10² µm²)
TH+ Bassoon+ area (x10 µm²)
DL, VM GPe
VL, DM GPe
d
RFP+ area (x10³ µm²)
RFP+ GABAAR+ area (x10² µm²)
e
Nearest neighbor distance
: RFP+ GABAAR+ to TH+ Bassoon+
Cumulative frequency (%)
Distance (µm)
DL, VM GPe
VL, DM GPe

**Extended Data Fig. 3 | The spatial relationship between putatively functional DA boutons and striatopallidal synapses**

**a,** A schematic illustration depicting fluorescently labeled striatopallidal axons (RFP), $GABA_AR$, dopaminergic axons (TH), and Bassoon. **b,** Representative enhanced confocal images of striatopallidal GABAergic synapses and presynaptic markers for DA boutons. **c,** Summary statistics of TH+ area (left) (n = 36 images from 3 mice per GPe subgroup, unpaired t-test; DL-VM GPe 202.39 ± 20.02 $\mu m^2$, VL-DM GPe 188.73 ± 13.40 $\mu m^2$) and Bassoon+ area colocalized with TH+ area (right) (DL-VM GPe 27.22 ± 44.44 $\mu m^2$, VL-DM GPe 19.59 ± 21.70 $\mu m^2$). **d,** Summary statistics of RFP+ area (left) (n = 36 images from 3 mice per GPe subgroup, unpaired t-test; DL-VM GPe 1054.57 ± 25.72 $\mu m^2$, VL-DM GPe 1071.03 ± 26.91 $\mu m^2$) and $GABA_AR$+ area colocalized with RFP+ area (right) (DL-VM GPe 146.65 ± 28.12 $\mu m^2$, VL-DM GPe 128.39 ± 25.75 $\mu m^2$). **e,** Cumulative plot for the nearest neighbor distance from striatopallidal GABAergic synapses (RFP+ $GABA_AR$+) to presynaptic DA boutons (TH+ Bassoon+) in the GPe subgroups. The data are presented as box-and-whisker plots.

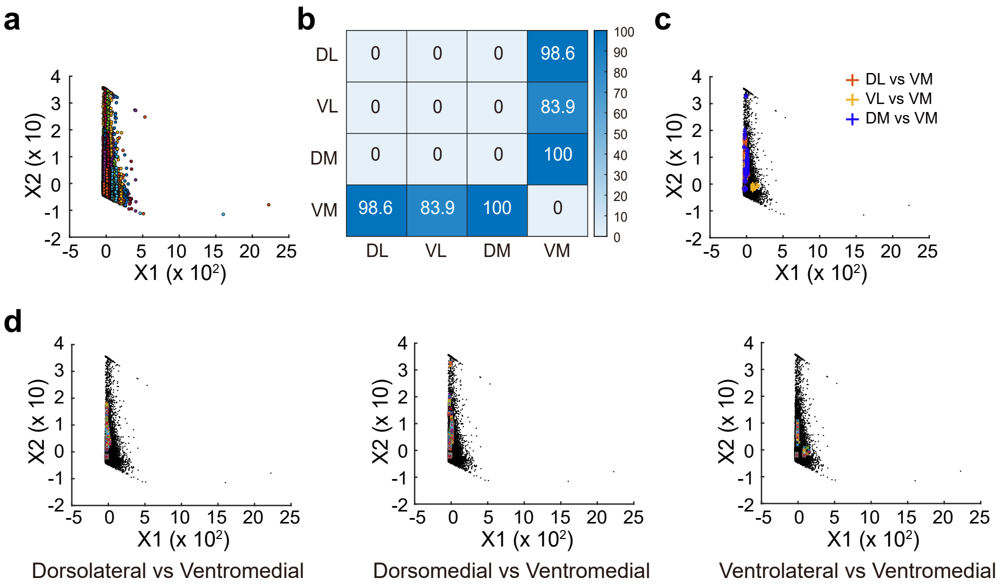
a
X2 (x 10)
X1 (x 10²)
b
DL
VL
DM
VM
0 0 0 98.6
0 0 0 83.9
0 0 0 100
98.6 83.9 100 0
c
DL vs VM
VL vs VM
DM vs VM
X2 (x 10)
X1 (x 10²)
d
Dorsolateral vs Ventromedial
Dorsomedial vs Ventromedial
Ventrolateral vs Ventromedial

**Extended Data Fig. 4 | An AI-based clustering analysis on $Sr^{2+}$-induced quantal responses in the GPe subregions**

**a,** A representative plot of striatopallidal quantal responses divided by two principal components, determined by rise, decay slope, and amplitude. **b,** A table showing the probability that the frequency of quantal responses from two different GPe subregions exhibits significant differences in more than one cluster among approximately 50 clusters. **c,** A representative merged plot of clusters exhibiting differences between the GPe subregions. **d,** Representative plots of clusters exhibiting differences between DL and VM GPe (left), DM and VM GPe (center), and VL and VM GPe (right).

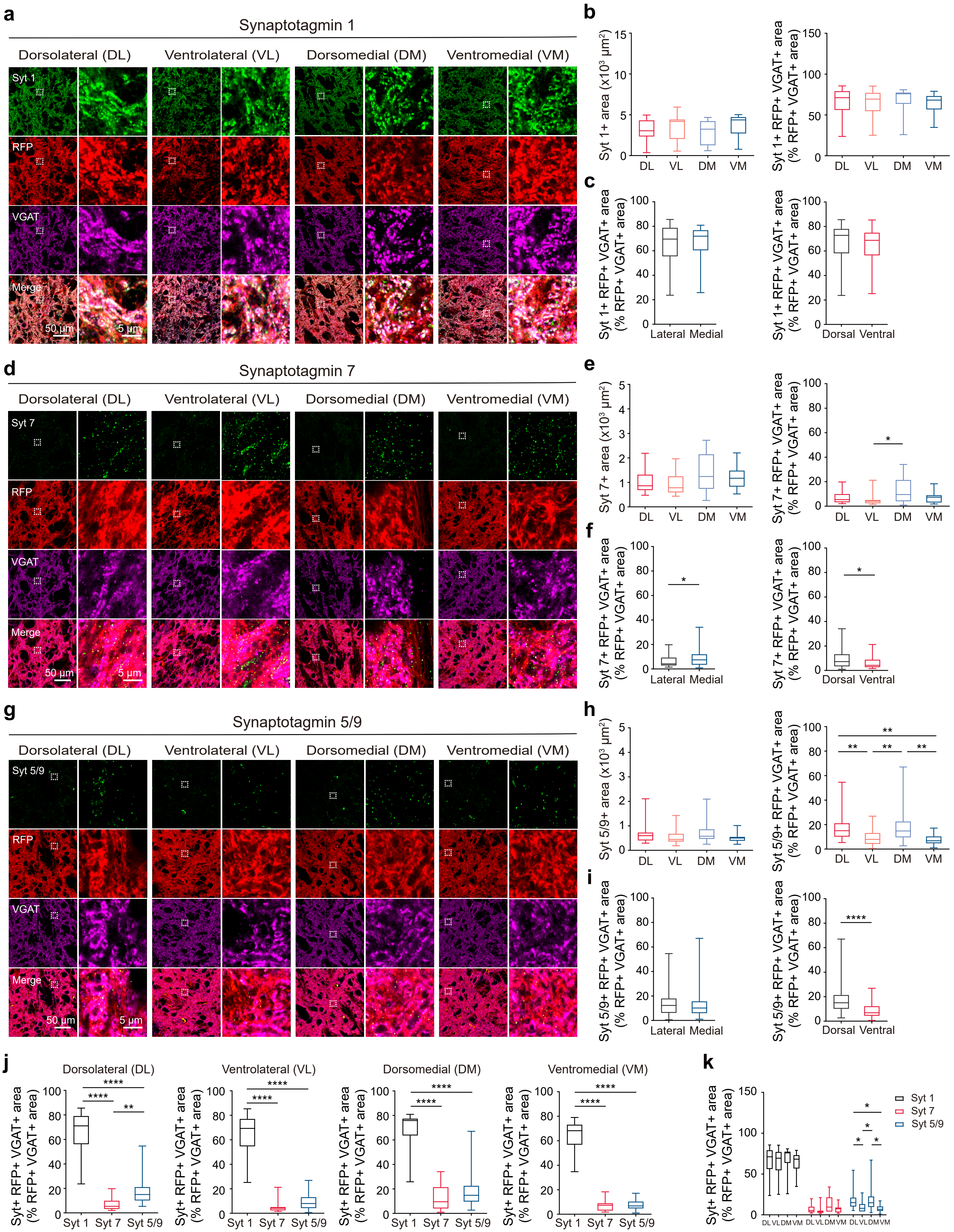

a
Synaptotagmin 1
Dorsolateral (DL)
Ventrolateral (VL)
Dorsomedial (DM)
Ventromedial (VM)
Syt 1
RFP
VGAT
Merge
50 µm
5 µm
b
Syt 1+ area (x10³ µm²)
Syt 1+ RFP+ VGAT+ area (% RFP+ VGAT+ area)
DL VL DM VM
c
Syt 1+ RFP+ VGAT+ area (% RFP+ VGAT+ area)
Lateral Medial
Dorsal Ventral
d
Synaptotagmin 7
Syt 7
e
Syt 7+ area (x10³ µm²)
Syt 7+ RFP+ VGAT+ area (% RFP+ VGAT+ area)
f
Syt 7+ RFP+ VGAT+ area (% RFP+ VGAT+ area)
g
Synaptotagmin 5/9
Syt 5/9
h
Syt 5/9+ area (x10³ µm²)
Syt 5/9+ RFP+ VGAT+ area (% RFP+ VGAT+ area)
i
Syt 5/9+ RFP+ VGAT+ area (% RFP+ VGAT+ area)
j
Dorsolateral (DL)
Ventrolateral (VL)
Dorsomedial (DM)
Ventromedial (VM)
Syt+ RFP+ VGAT+ area (% RFP+ VGAT+ area)
Syt 1 Syt 7 Syt 5/9
k
Syt+ RFP+ VGAT+ area (% RFP+ VGAT+ area)
Syt 1
Syt 7
Syt 5/9
DL VL DM VM

**Extended Data Fig. 5 | Regional variations in the expression of synaptotagmin subtypes at striatopallidal axon terminals in the subregions of the GPe**

**a,d,g,** Representative enhanced confocal images of synaptotagmins (1, 7, 5/9), RFP, and VGAT immunofluorescence in the GPe subregions of Adora2A-Cre;Ai9 mice. **b,e,h,** Summary statistics of synaptotagmin+ area (left) and synaptotagmin+ area on striatopallidal GABAergic presynaptic terminals, normalized to the total area of striatopallidal GABAergic presynaptic terminals (right) in the GPe subregions. **c,f,i,** Summary statistics of synaptotagmin+ area on striatopallidal GABAergic presynaptic terminals, normalized to the total area of striatopallidal GABAergic presynaptic terminals in the lateral-medial GPe (left) and dorsal-ventral GPe (right). **b,** Summary statistics of synaptotagmin 1+ area (left) (n = 20 images from 3 mice per GPe subregion, one-way ANOVA; DL 3104.4 ± 286.2 $\mu m^2$, VL 3589.3 ± 365.3 $\mu m^2$, DM 2906 ± 321 $\mu m^2$, VM 3700 ± 324.4 $\mu m^2$; p = 0.2599) and normalized synaptotagmin 1+ RFP+ VGAT+ area (right) (DL 64.3 ± 4.1%, VL 63.7 ± 3.9%, DM 65.7 ± 4.5%, VM 65.1 ± 2.8%; p = 0.9859). **c,** Summary statistics of normalized synaptotagmin 1+ area in the lateral-medial GPe (left) (n = 40 images from 6 mice per GPe subgroup, unpaired t-test; lateral GPe 64.0 ± 2.8%, medial GPe 65.4 ± 2.6%) and dorsal-ventral GPe (right) (dorsal GPe 65 ± 3%, medial GPe 64.4 ± 2.4%). **e,** Summary statistics of synaptotagmin 7+ area (left) (n = 20 images from 3 mice per GPe subregion, one-way ANOVA; DL 1026.3 ± 101.7 $\mu m^2$, VL 969.3 ± 114.2 $\mu m^2$, DM 1349.5 ± 151.8 $\mu m^2$, VM 1192 ± 97.6 $\mu m^2$; p = 0.1129) and normalized synaptotagmin 7+ RFP+ VGAT+ area (right) (DL 7.4 ± 1.2%, VL 5.6 ± 1.2%, DM 11.7 ± 2.0%, VM 6.6 ± 0.9%; p = 0.0174). **f,** Summary statistics of normalized synaptotagmin 7+ area in the lateral-medial GPe (left) (n = 40 images from 6 mice per GPe subgroup, unpaired t-test; lateral GPe 6.1 ± 0.8%, medial GPe 9.2 ± 1.2%) and dorsal-ventral GPe (right) (dorsal GPe 9.7 ± 1.2%, medial GPe 6.1 ± 0.8%). **h,** Summary statistics of synaptotagmin 5/9+ area (left) (n = 25 images from 3 mice per GPe subregion, one-way ANOVA; DL 698.1 ± 84.7 $\mu m^2$, VL 554 ± 60 $\mu m^2$, DM 722 ± 86.8 $\mu m^2$, VM 500.5 ± 32.9 $\mu m^2$; p = 0.0733) and normalized synaptotagmin 5/9+ RFP+ VGAT+ area (right) (DL 18.0 ± 2.4%, VL 9.1 ± 1.3%, DM 18.2 ± 2.8%, VM 7.7 ± 0.9%; p = 0.0001). **i,** Summary statistics of normalized synaptotagmin 5/9+ area in the lateral-medial GPe (left) (n = 50 images from 6 mice per GPe subgroup, unpaired t-test; lateral GPe 13.5 ± 1.5%, medial GPe 13.1 ± 1.6%) and dorsal-ventral GPe (right) (dorsal GPe 18.1 ± 1.8%, medial GPe 8.4 ± 0.8%). **j,** Summary statistics of synaptotagmin (1, 7, 5/9)+ area on striatopallidal GABAergic presynaptic terminals, normalized to the total area of striatopallidal

GABAergic presynaptic terminals in the DL GPe (one-way ANOVA; Syt-1 64.3 ± 4.1%, Syt-7 7.4 ± 1.2%, Syt-5/9 18.0 ± 2.4%; $p < 0.0001$), VL GPe (Syt-1 63.7 ± 3.9%, Syt-7 5.6 ± 1.2%, Syt-5/9 9.1 ± 1.3%; $p < 0.0001$), DM GPe (Syt-1 65.7 ± 4.5%, Syt-7 11.7 ± 2.0%, Syt-5/9 18.2 ± 2.8%; $p < 0.0001$), and VM GPe (Syt-1 65.1 ± 2.8%, Syt-7 6.6 ± 1.0%, Syt-5/9 7.7 ± 1.0%; $p < 0.0001$). **k,** Summary statistics of synaptotagmin (1, 7, 5/9)+ area on striatopallidal GABAergic presynaptic terminals, normalized to the total area of striatopallidal GABAergic presynaptic terminals in the GPe subregions (ordinary two-way ANOVA with Holm-Sidak's post-hoc multiple comparisons test; Syt-1, DL 64.3 ± 4.1%, VL 63.7 ± 3.9%, DM 65.6 ± 4.4%, VM 65.1 ± 2.8%; Syt-7, DL 7.4 ± 1.2%, VL 5.6 ± 1.2%, DM 11.7 ± 2.0%, VM 6.6 ± 0.9%; Syt-5/9, DL 18.0 ± 2.4%, VL 9.1 ± 1.3%, DM 18.2 ± 2.8%, VM 7.7 ± 0.9%; GPe subregion, $p = 0.0149$, synaptotagmin subtype, $p < 0.0001$, interaction, $p = 0.2354$). The data are presented as box-and-whisker plots. $^{*}p < 0.05$, $^{**}p < 0.01$, $^{****}p < 0.0001$.

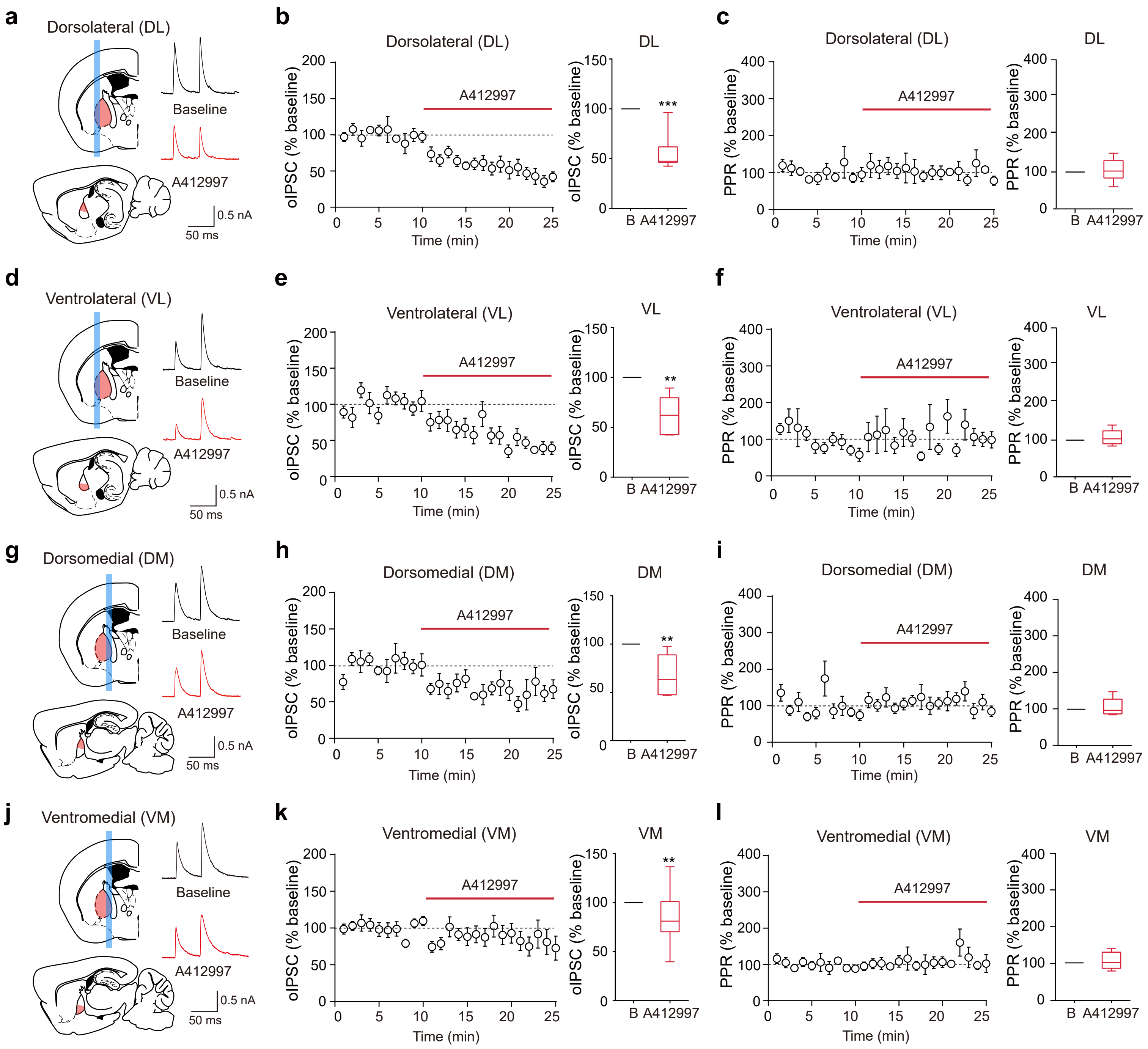
a
Dorsolateral (DL)
Baseline
A412997
0.5 nA
50 ms
b
Dorsolateral (DL)
A412997
oIPSC (% baseline)
Time (min)
DL
***
B
A412997
c
Dorsolateral (DL)
A412997
PPR (% baseline)
Time (min)
DL
d
Ventrolateral (VL)
Baseline
A412997
0.5 nA
50 ms
e
Ventrolateral (VL)
A412997
oIPSC (% baseline)
Time (min)
VL
**
f
Ventrolateral (VL)
PPR (% baseline)
VL
g
Dorsomedial (DM)
Baseline
A412997
h
Dorsomedial (DM)
DM
**
i
Dorsomedial (DM)
DM
j
Ventromedial (VM)
Baseline
A412997
k
Ventromedial (VM)
VM
**
l
Ventromedial (VM)
VM

**Extended Data Fig. 6 | The selective D4R agonist A-412997 universally attenuates striatopallidal transmission across all subregions of the GPe, without affecting the PPR**

**a,d,g,j,** Schematic illustrations of the GPe subregions and representative recording traces before and after the bath application of A-412997 (100 nM). **b,e,h,k,** Normalized oIPSC amplitude plots (left) and summary statistics of normalized oIPSCs (right) in the GPe subregions. **c,f,i,l,** Normalized PPR plots (left) and summary statistics of normalized PPR (right) in the GPe subregions. **b,c,** Summary statistics of normalized oIPSCs (n = 7 cells from 4 mice, one sample t-test; Baseline 100%, A-412997 56.44 ± 7.04%) and PPRs (Baseline 100%, A-412997 103.02 ± 10.93%) in the DL GPe. **e,f,** Summary statistics of normalized oIPSCs (n = 6 cells from 6 mice, one sample t-test; Baseline 100%, A-412997 62.47 ± 8.17%) and PPRs (Baseline 100%, A-412997 107.85 ± 8.15%) in the VL GPe. **h,i,** Summary statistics of normalized oIPSCs (n = 7 cells from 4 mice, one sample t-test; Baseline 100%, A-412997 67.77 ± 8.13%) and PPRs (Baseline 100%, A-412997 107.65 ± 9.13%) in the DM GPe. **k,l,** Summary statistics of normalized oIPSCs (n = 11 cells from 7 mice, one sample t-test; Baseline 100%, A-412997 75.54 ± 5.39%) and PPRs (Baseline 100%, A-412997 109.47 ± 7.89%) in the DM GPe. The data are presented as box-and-whisker plots. **p < 0.01, ***p < 0.001.

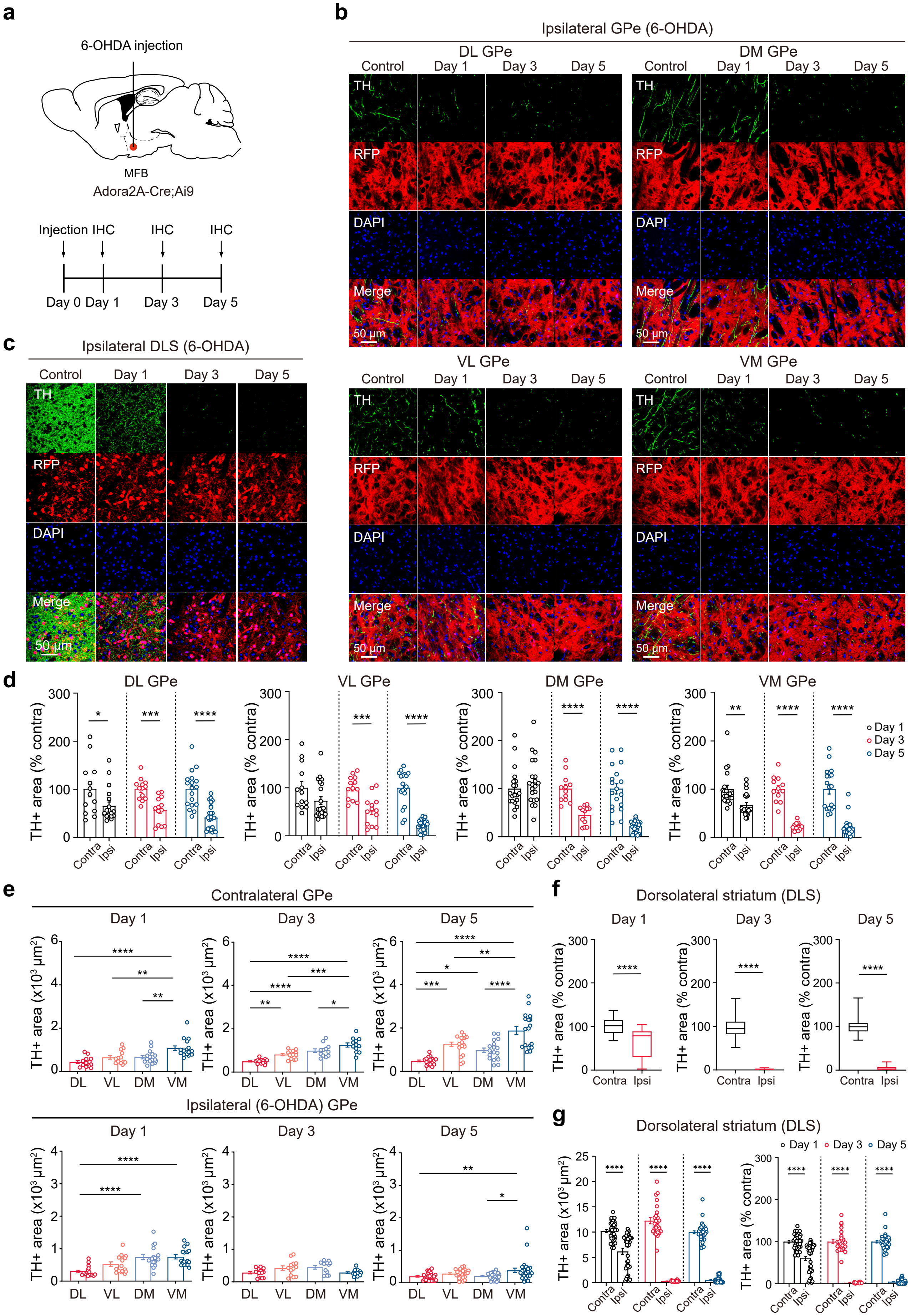
a
6-OHDA injection
MFB
Adora2A-Cre;Ai9
Injection IHC
IHC
IHC
Day 0 Day 1
Day 3
Day 5
b
Ipsilateral GPe (6-OHDA)
DL GPe
DM GPe
VL GPe
VM GPe
Control
Day 1
Day 3
Day 5
TH
RFP
DAPI
Merge
50 µm
c
Ipsilateral DLS (6-OHDA)
d
DL GPe
VL GPe
DM GPe
VM GPe
TH+ area (% contra)
Contra
Ipsi
Day 1
Day 3
Day 5
e
Contralateral GPe
Ipsilateral (6-OHDA) GPe
TH+ area (x10³ µm²)
DL
VL
DM
VM
f
Dorsolateral striatum (DLS)
g
Dorsolateral striatum (DLS)

**Extended Data Fig. 7 | 6-OHDA-induced degeneration of dopaminergic axons exhibits regional heterogeneity across the subregions of the GPe**

**a,** A schematic illustration describing the unilateral injection of 6-OHDA into the MFB of Adora2A-Cre;Ai9 mice. **b,** Representative confocal images of striatopallidal axons (RFP) and TH+ dopaminergic axons in the GPe subregions of ipsilateral (6-OHDA) hemisphere at 1, 3, and 5 days post-6-OHDA injection. **c,** Representative confocal images of TH+ dopaminergic axons and iMSNs (RFP) in the DLS of ipsilateral (6-OHDA) hemispheres at 1, 3, and 5 days post-6-OHDA injection. **d,** Summary statistics of TH+ area normalized to the contralateral TH+ area in the DL GPe (n = 13-18 images from 4 mice at each time point, unpaired t-test; Day 1 Contra 100 ± 15.24%, Day 1 Ipsi 65.76 ± 7.92%, $p = 0.0381$; Day 3 Contra 100 ± 6.06%, Day 3 Ipsi 57.09 ± 7.59%, $p = 0.0002$; Day 5 Contra 100 ± 8.85%, Day 5 Ipsi 41.34 ± 4.96%, $p < 0.0001$), VL GPe (n = 13-26 images from 3-5 mice at each time point; Day 1 Contra 100 ± 12.90%, Day 1 Ipsi 73.06 ± 8.28%, $p = 0.0743$; Day 3 Contra 100 ± 6.70%, Day 3 Ipsi 52.52 ± 8.19%, $p = 0.0002$; Day 5 Contra 100 ± 8.43%, Day 5 Ipsi 22.62 ± 2.14%, $p < 0.0001$), DM GPe (n = 12-26 images from 3-5 mice at each time point; Day 1 Contra 100 ± 9.31%, Day 1 Ipsi 113.90 ± 10.31%, $p = 0.3250$; Day 3 Contra 100 ± 8.46%, Day 3 Ipsi 45.18 ± 5.51%, $p < 0.0001$; Day 5 Contra 100 ± 11.02%, Day 5 Ipsi 21.41 ± 2.15%, $p < 0.0001$), and VM GPe (n = 12-26 images from 3-5 mice at each time point; Day 1 Contra 100 ± 8.08%, Day 1 Ipsi 67.33 ± 5.12%, $p = 0.0014$; Day 3 Contra 100 ± 7.72%, Day 3 Ipsi 22.74 ± 2.28%, $p < 0.0001$; Day 5 Contra 100 ± 10.07%, Day 5 Ipsi 20.35 ± 3.61%, $p < 0.0001$). **e,** Summary statistics of TH+ area in the contralateral GPe (top) 1 day after 6-OHDA injection (n = 13-16 images from 4 mice per GPe subregion, one-way ANOVA; DL 433.6 ± 66.1 $\mu m^2$, VL 652.1 ± 84.1 $\mu m^2$, DM 655.5 ± 73.5 $\mu m^2$, VM 1074.0 ± 104.4 $\mu m^2$; $p < 0.0001$), 3 days after 6-OHDA injection (n = 12-13 images from 3 mice per GPe subregion; DL 494 ± 29.9 $\mu m^2$, VL 810.7 ± 54.4 $\mu m^2$, DM 990.4 ± 83.8 $\mu m^2$, VM 1252 ± 96.6 $\mu m^2$; $p < 0.0001$), and 5 days after 6-OHDA injection (n = 18 images from 4 mice per GPe subregion; DL 476.8 ± 42.2 $\mu m^2$, VL 1240 ± 104.5 $\mu m^2$, DM 965.8 ± 106.4 $\mu m^2$, VM 1877 ± 189 $\mu m^2$; $p < 0.0001$). Summary statistics of TH+ area in the ipsilateral GPe (bottom) 1 day after 6-OHDA injection (n = 16 images from 4 mice per GPe subregion, one-way ANOVA; DL 297.8 ± 40.1 $\mu m^2$, VL 525.1 ± 66.3 $\mu m^2$, DM 739.8 ± 81.0 $\mu m^2$, VM 748.4 ± 65.6 $\mu m^2$; $p < 0.0001$), 3 days after 6-OHDA injection (n = 12-13 images from 3 mice per GPe subregion; DL 282 ± 37.5 $\mu m^2$, VL 425.7 ± 66.4 $\mu m^2$, DM 447.4 ± 54.6 $\mu m^2$, VM 284.6 ± 28.5 $\mu m^2$; $p = 0.0313$), and 5 days after 6-OHDA injection (n = 26 images from 5 mice per GPe subregion; DL 197.1 ± 23.6 $\mu m^2$, VL 280.5 ± 26.6 $\mu m^2$, DM 206.8 ± 20.7 $\mu m^2$, VM 381.9 ± 67.7 $\mu m^2$; $p = 0.0044$). **f,** Summary statistics of TH+ area normalized to the contralateral TH+ area in the DLS 1 day after 6-OHDA injection (n = 35 images from 5 mice per each hemisphere, unpaired t-test; Contra 100 ± 3.07%, Ipsi 60.26 ± 5.26%), 3 days after 6-OHDA injection (n = 24 images from 4 mice per each hemisphere; Contra 100 ± 5.07%, Ipsi 2.25 ± 0.21%), and 5 day after 6-OHDA injection (n = 36 images from 5 mice per each hemisphere; Contra 100 ± 2.92%, Ipsi 4.80 ± 0.64%). **g,** Summary statistics of TH+ area in the contralateral and ipsilateral DLS (left) (n = 24-36 images from 4-5 mice per each hemisphere at each time point, ordinary two-way ANOVA with Holm-Sidak's post-hoc multiple comparisons test; Day 1 Contra 10194.34 ± 313.41 $\mu m^2$, Day 1 Ipsi 6143 ± 536.32 $\mu m^2$, Day 3 Contra 12232.27 ± 619.55 $\mu m^2$, Day 3 Ipsi 275.46 ± 25.24 $\mu m^2$, Day 5 Contra 9950.94 ± 290.40 $\mu m^2$, Day 5 Ipsi 477.20 ± 63.38 $\mu m^2$; 6-OHDA, $p < 0.0001$, post-injection time point, $p < 0.0001$, interaction, $p < 0.0001$) and TH+ area of the contralateral and ipsilateral DLS normalized to the contralateral TH+ area (right) (n = 24-36 images from 4-5 mice per each hemisphere at each time point, unpaired t-test; Day 1 Contra 100 ± 3.07%, Day 1 Ipsi 60.26 ± 5.26%, $p < 0.0001$; Day 3 Contra 100 ± 5.07%, Day 3 Ipsi 2.25 ± 0.21%, $p < 0.0001$; Day 5 Contra 100 ± 2.92%, Day 5 Ipsi 4.80 ± 0.64%, $p < 0.0001$) at 1, 3, and 5 days post-6-OHDA injection. The data are presented as box-and-whisker plots or mean ± SEM. *$p < 0.05$, **$p < 0.01$, ***$p < 0.001$, ****$p < 0.0001$.

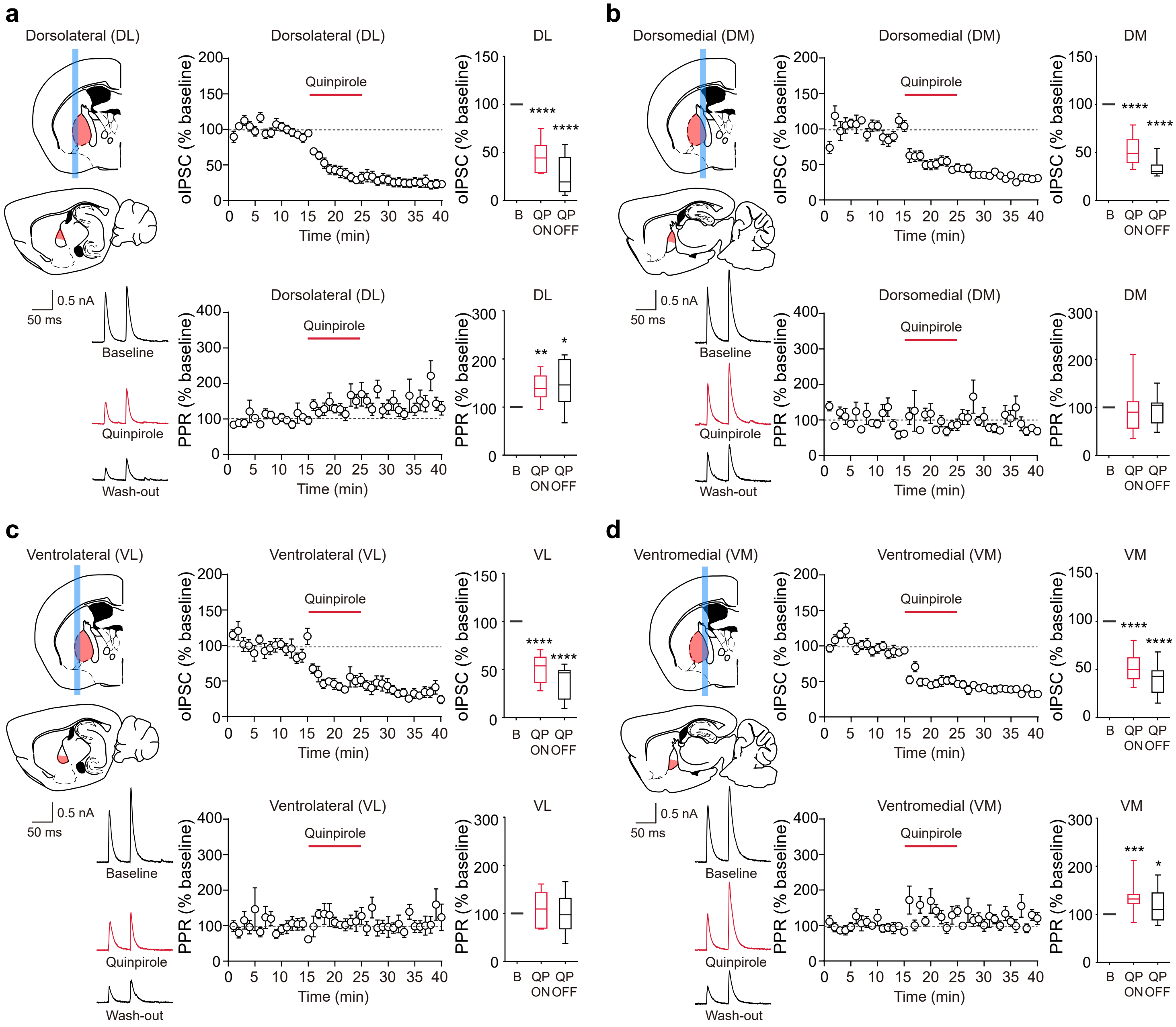

a
Dorsolateral (DL)
0.5 nA
50 ms
Baseline
Quinpirole
Wash-out
Dorsolateral (DL)
Quinpirole
oIPSC (% baseline)
Time (min)
DL
****
****
B
QP ON
QP OFF
Dorsolateral (DL)
PPR (% baseline)
**
*
b
Dorsomedial (DM)
Quinpirole
oIPSC (% baseline)
PPR (% baseline)
Time (min)
DM
****
****
B
QP ON
QP OFF
0.5 nA
50 ms
Baseline
Quinpirole
Wash-out
c
Ventrolateral (VL)
Quinpirole
oIPSC (% baseline)
PPR (% baseline)
Time (min)
VL
****
****
B
QP ON
QP OFF
0.5 nA
50 ms
Baseline
Quinpirole
Wash-out
d
Ventromedial (VM)
Quinpirole
oIPSC (% baseline)
PPR (% baseline)
Time (min)
VM
****
****
***
*
B
QP ON
QP OFF
0.5 nA
50 ms
Baseline
Quinpirole
Wash-out

**Extended Data Fig. 8 | Striatopallidal synaptic transmission in the contralateral hemisphere of 6-OHDA-lesioned mice recapitulates region-specific dopaminergic modulation in the subregions of the GPe**

**a-d,** Schematic illustrations of the GPe subregions of the contralateral hemisphere (top left), representative recording traces (bottom left), oIPSC amplitude (top center) and PPR plots (bottom center) normalized to baseline during bath application of quinpirole (10 μM), and summary statistics of normalized oIPSCs (top right) and PPRs (bottom right). **a,** Summary statistics of normalized oIPSCs (n = 10 cells from 9 mice, one sample t-test; Baseline 100%, QP ON 44.34 ± 5.09%, QP OFF 26.77 ± 6.32%) and PPRs (Baseline 100%, QP ON 139.90 ± 9.38%, QP OFF 147 ± 14.87%) in the DL GPe. **b,** Summary statistics of normalized oIPSCs (n = 11 cells from 11 mice, one sample t-test; Baseline 100%, QP ON 53.38 ± 4.65%, QP OFF 34.37 ± 2.79%) and PPRs (Baseline 100%, QP ON 95.11 ± 13.94%, QP OFF 97.97 ± 9.61%) in the DM GPe. **c,** Summary statistics of normalized oIPSCs (n = 9 cells from 9 mice, one sample t-test; Baseline 100%, QP ON 51.45 ± 4.93%, QP OFF 37.66 ± 5.80%) and PPRs (Baseline 100%, QP ON 108.90 ± 12.28%, QP OFF 99.99 ± 14.06%) in the VL GPe. **d,** Summary statistics of normalized oIPSCs (n = 15 cells from 12 mice, one sample t-test; Baseline 100%, QP ON 51.44 ± 3.43%, QP OFF 39.53 ± 3.71%) and PPRs (Baseline 100%, QP ON 134 ± 8.13%, QP OFF 117.30 ± 8.07%) in the VM GPe. The data are presented as box-and-whisker plots. $*p < 0.05$, $**p < 0.01$, $***p < 0.001$, $****p < 0.0001$.

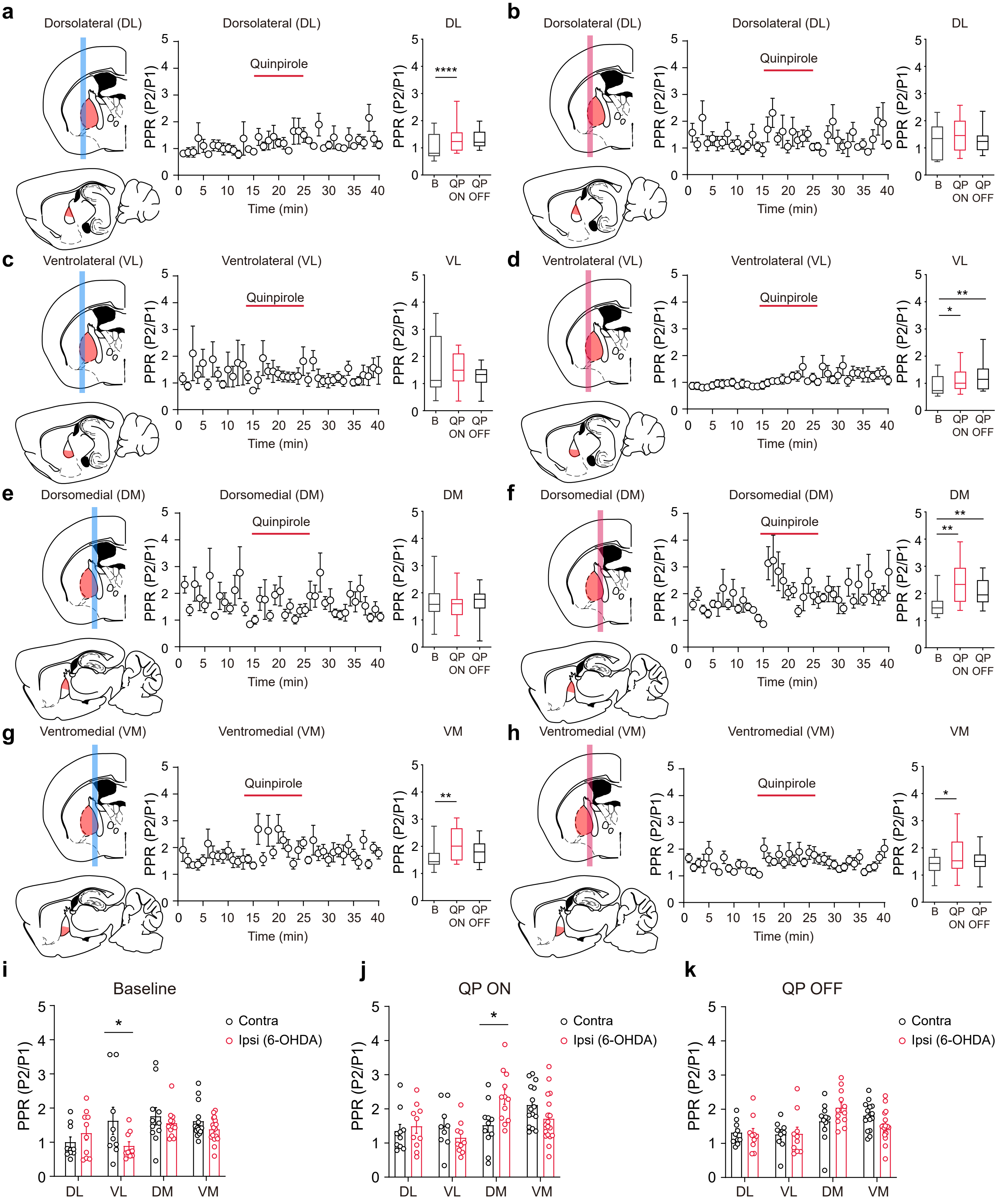

a
Dorsolateral (DL)
Quinpirole
PPR (P2/P1)
Time (min)
DL
****
B
QP ON
QP OFF
b
Dorsolateral (DL)
Quinpirole
DL
c
Ventrolateral (VL)
Quinpirole
VL
d
Ventrolateral (VL)
Quinpirole
VL
*
**
e
Dorsomedial (DM)
Quinpirole
DM
f
Dorsomedial (DM)
Quinpirole
DM
**
**
g
Ventromedial (VM)
Quinpirole
VM
**
h
Ventromedial (VM)
Quinpirole
VM
*
i
Baseline
Contra
Ipsi (6-OHDA)
DL
VL
DM
VM
*
j
QP ON
Contra
Ipsi (6-OHDA)
*
k
QP OFF
Contra
Ipsi (6-OHDA)

**Extended Data Fig. 9 | DA depletion negates quinpirole-induced PPR elevation in the DL-VM subregions of the GPe while increasing PPR in the VL-DM GPe subregions**

**a-h,** Schematic illustrations of the GPe subregions of the contralateral hemisphere or ipsilateral (6-OHDA) hemisphere (left), PPR plots (middle), and summary statistics of PPR (right) before and after bath application of quinpirole (10 μM). **a,** Summary statistics of PPRs (n = 10 cells from 9 mice, repeated measures one-way ANOVA with Holm-Sidak's post-hoc multiple comparisons test; Baseline 1 ± 0.15, QP ON 1.35 ± 0.19, QP OFF 1.32 ± 0.11, p = 0.044) in the contralateral DL GPe. **b,** Summary statistics of PPRs (n = 10 cells from 10 mice; Baseline 1.27 ± 0.21, QP ON 1.49 ± 0.21, QP OFF 1.28 ± 0.16, p = 0.3099) in the ipsilateral DL GPe. **c,** Summary statistics of PPRs (n = 9 cells from 9 mice; Baseline 1.63 ± 0.40, QP ON 1.50 ± 0.22, QP OFF 1.26 ± 0.14, p = 0.3736) in the contralateral VL GPe. **d,** Summary statistics of PPRs (n = 10 cells from 9 mice; Baseline 0.89 ± 0.12, QP ON 1.15 ± 0.15, QP OFF 1.28 ± 0.20, p = 0.0027) in the ipsilateral VL GPe. **e,** Summary statistics of PPRs (n = 11 cells from 11 mice; Baseline 1.76 ± 0.25, QP ON 1.52 ± 0.20, QP OFF 1.64 ± 0.17, p = 0.5647) in the contralateral DM GPe. **f,** Summary statistics of PPRs (n = 12 cells from 12 mice; Baseline 1.56 ± 0.12, QP ON 2.35 ± 0.21, QP OFF 2.05 ± 0.15, p = 0.0012) in the ipsilateral DM GPe. **g,** Summary statistics of PPRs (n = 15 cells from 12 mice; Baseline 1.61 ± 0.13, QP ON 2.11 ± 0.16, QP OFF 1.81 ± 0.11, p = 0.0038) in the contralateral VM GPe. **h,** Summary statistics of PPRs (n = 18 cells from 16 mice; Baseline 1.38 ± 0.08, QP ON 1.71 ± 0.16, QP OFF 1.52 ± 0.10, p = 0.0270) in the ipsilateral VM GPe. **i,** Summary statistics of PPRs in the GPe subregions of the contralateral and ipsilateral hemispheres (ordinary two-way ANOVA with Holm-Sidak's post-hoc multiple comparisons test; Contra DL 1 ± 0.15, Ipsi DL 1.27 ± 0.21, Contra VL 1.63 ± 0.40, Ipsi VL 0.90 ± 0.12, Contra DM 1.76 ± 0.25, Ipsi DM 1.55 ± 0.12, Contra VM 1.61 ± 0.13, Ipsi VM 1.38 ± 0.08; GPe subregion, p = 0.0294, 6-OHDA, p = 0.0846, interaction, p = 0.1024) during baseline recording. **j,** Summary statistics of PPRs in the GPe subregions of the contralateral and ipsilateral hemispheres (ordinary two-way ANOVA with Holm-Sidak's post-hoc multiple comparisons test; Contra DL 1.35 ± 0.19, Ipsi DL 1.50 ± 0.21, Contra VL 1.50 ± 0.22, Ipsi VL 1.15 ± 0.15, Contra DM 1.52 ± 0.20, Ipsi DM 2.35 ± 0.22, Contra VM 2.11 ± 0.16, Ipsi VM 1.71 ± 0.16; GPe subregion, p = 0.0017, 6-OHDA, p = 0.6825, interaction, p = 0.0045) during QP application. **k,** Summary statistics of PPRs in the GPe subregions of the contralateral and ipsilateral hemispheres (ordinary two-way ANOVA with Holm-Sidak's post-hoc multiple comparisons test; Contra DL 1.32 ± 0.11, Ipsi DL 1.28 ± 0.16, Contra VL 1.27 ± 0.14,

Ipsi VL 1.28 ± 0.20, Contra DM 1.64 ± 0.17, Ipsi DM 2.05 ± 0.15, Contra VM 1.81 ± 0.11, Ipsi VM 1.52 ± 0.10; GPe subregion, $p = 0.0002$, 6-OHDA, $p = 0.8181$, interaction, $p = 0.0764$) after QP application. The data are presented as box-and-whisker plots or mean ± SEM. *$p < 0.05$, **$p < 0.01$, ****$p < 0.0001$.

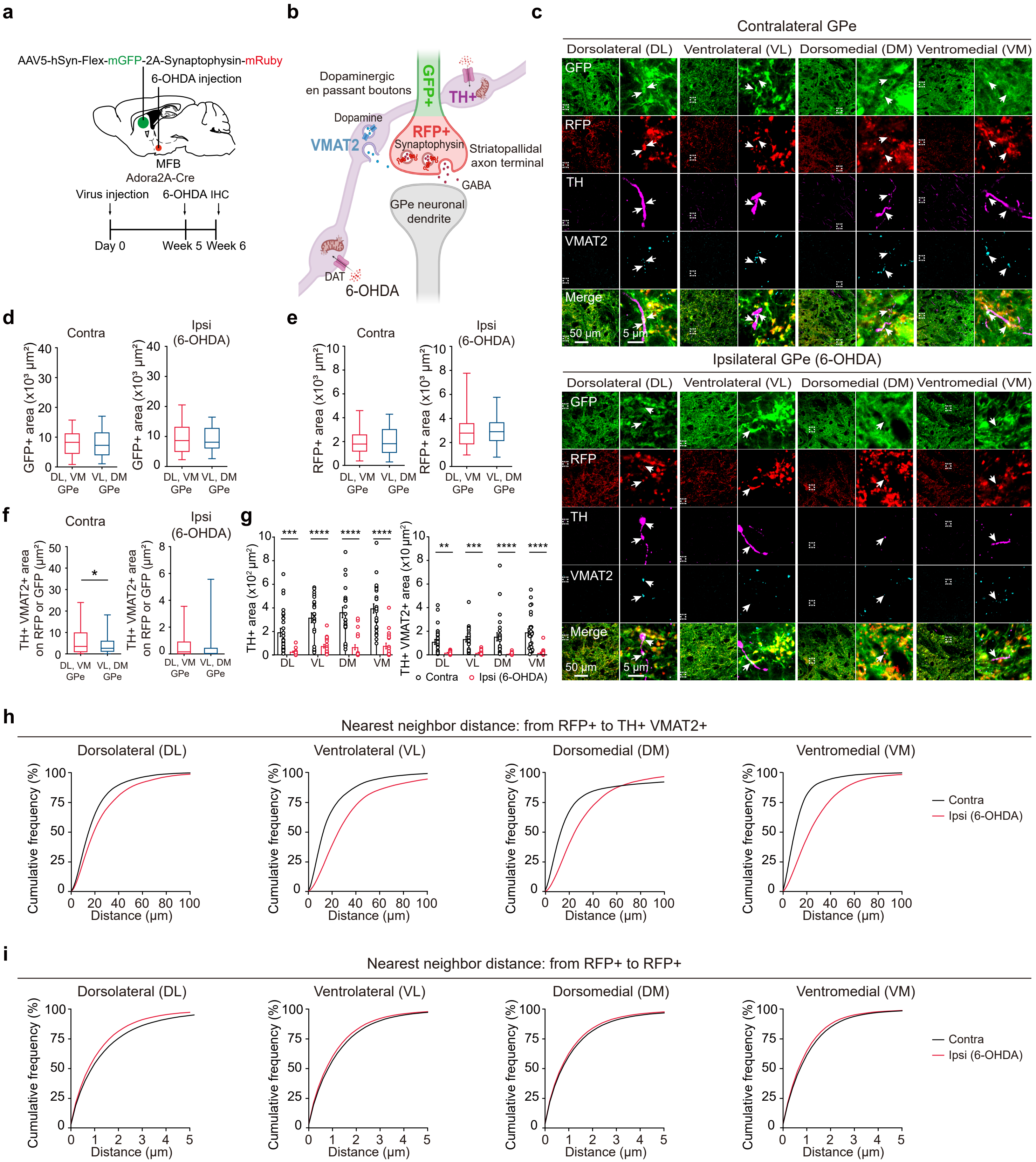

a
AAV5-hSyn-Flex-mGFP-2A-Synaptophysin-mRuby
6-OHDA injection
MFB
Adora2A-Cre
Virus injection
6-OHDA IHC
Day 0
Week 5
Week 6
b
Dopaminergic en passant boutons
GFP+
TH+
Dopamine
VMAT2
RFP+
Synaptophysin
Striatopallidal axon terminal
GABA
GPe neuronal dendrite
DAT
6-OHDA
c
Contralateral GPe
Dorsolateral (DL)
Ventrolateral (VL)
Dorsomedial (DM)
Ventromedial (VM)
GFP
RFP
TH
VMAT2
Merge
50 μm
5 μm
Ipsilateral GPe (6-OHDA)
d
Contra
Ipsi (6-OHDA)
GFP+ area (x10³ μm²)
DL, VM GPe
VL, DM GPe
e
RFP+ area (x10³ μm²)
f
TH+ VMAT2+ area on RFP or GFP (μm²)
*
g
TH+ area (x10² μm²)
TH+ VMAT2+ area (x10 μm²)
***
****
**
DL
VL
DM
VM
Contra
Ipsi (6-OHDA)
h
Nearest neighbor distance: from RFP+ to TH+ VMAT2+
Cumulative frequency (%)
Distance (μm)
i
Nearest neighbor distance: from RFP+ to RFP+

**Extended Data Fig. 10 | No significant differences in striatopallidal axons and axon terminals were observed between the GPe subgroups after DA depletion**

**a,** A schematic illustration depicting the injection of AAV5-hSyn-Flex-mGFP-2A-Synaptophysin-mRuby virus into the striatum of Adora2A-Cre mice, along with the unilateral injection of 6-OHDA into the MFB. **b,** A schematic illustration depicting fluorescently labeled striatopallidal axons (GFP), axon terminals (RFP), dopaminergic axons (TH), and vesicular monoamine transporter 2 (VMAT2) with 6-OHDA injection. **c,** Representative enhanced confocal images of striatopallidal axon terminals and potential dopaminergic boutons (TH+ VMAT2+) in the GPe subregions of the contralateral and ipsilateral (6-OHDA) hemispheres. **d,** Summary statistics of virally expressed GFP+ area in the GPe subgroups of the contralateral (left) ($n = 46$ images from 7 mice per GPe subgroup, unpaired t-test; DL-VM GPe $7755.1 \pm 543.2$ $\mu m^2$, VL-DM GPe $7922 \pm 677.9$ $\mu m^2$) and ipsilateral hemispheres (right) (DL-VM GPe $9384.3 \pm 766.8$ $\mu m^2$, VL-DM GPe $8864.2 \pm 588.1$ $\mu m^2$). **e,** Summary statistics of virally expressed RFP+ area in the GPe subgroups of the contralateral (left) ($n = 46$ images from 7 mice per GPe subgroup, unpaired t-test; DL-VM GPe $1934.5 \pm 136.1$ $\mu m^2$, VL-DM GPe $2005.9 \pm 170.4$ $\mu m^2$) and ipsilateral hemispheres (right) (DL-VM GPe $3017.2 \pm 235.8$ $\mu m^2$, VL-DM GPe $2912.4 \pm 171.8$ $\mu m^2$). **f,** Summary statistics of VMAT2+ area colocalized with TH+ area on striatopallidal axons (RFP+ or GFP+) in the GPe subgroups of the contralateral (left) ($n = 46$ images from 7 mice per GPe subgroup, unpaired t-test; DL-VM GPe $6.4 \pm 0.9$ $\mu m^2$, VL-DM GPe $3.9 \pm 0.6$ $\mu m^2$) and ipsilateral hemispheres (right) (DL-VM GPe $0.6 \pm 0.1$ $\mu m^2$, VL-DM GPe $0.4 \pm 0.1$ $\mu m^2$). **g,** Summary statistics of TH+ area (left) (ordinary two-way ANOVA with Holm-Sidak's post-hoc multiple comparisons test; Contra DL $190.1 \pm 34.3$ $\mu m^2$, Ipsi DL $25.4 \pm 5.6$ $\mu m^2$, Contra VL $314 \pm 40.8$ $\mu m^2$, Ipsi VL $73.2 \pm 14.7$ $\mu m^2$, Contra DM $360.2 \pm 44.4$ $\mu m^2$, Ipsi DM $64.5 \pm 21.2$ $\mu m^2$, Contra VM $393.2 \pm 38.7$ $\mu m^2$, Ipsi VM $75.4 \pm 23.9$ $\mu m^2$; GPe subregion, $p = 0.0009$, 6-OHDA, $p < 0.0001$, interaction, $p = 0.0911$) and VMAT2+ area colocalized with TH+ area (right) (Contra DL $11 \pm 2.1$ $\mu m^2$, Ipsi DL $1.7 \pm 0.4$ $\mu m^2$, Contra VL $13.2 \pm 2.3$ $\mu m^2$, Ipsi VL $1.6 \pm 0.4$ $\mu m^2$, Contra DM $15.3 \pm 3.5$ $\mu m^2$, Ipsi DM $0.7 \pm 0.2$ $\mu m^2$, Contra VM $19 \pm 3.1$ $\mu m^2$, Ipsi VM $1.5 \pm 0.7$ $\mu m^2$; GPe subregion, $p = 0.3448$, 6-OHDA, $p < 0.0001$, interaction, $p = 0.2763$) in the GPe subregions of the contralateral and ipsilateral hemispheres. **h,i,** Cumulative plots of the nearest neighbor distances from striatopallidal axon terminals (RFP+) to potential dopaminergic boutons (TH+ VMAT2+) or to other striatopallidal axon terminals (RFP+) in the GPe subregions of the contralateral and ipsilateral hemispheres. The data are presented as box-and-whisker plots or mean ± SEM. $^{*}p < 0.05$, $^{**}p < 0.01$, $^{***}p < 0.001$, $^{****}p < 0.0001$.

**Table S1. oIPSC kinetics in striatopallidal synaptic transmission within the GPe**

| | PV+ | FoxP2+ | PV-/FoxP2- |
|---|---|---|---|
| **Baseline PPR (P2/P1)** | 1.63 ± 0.06 | 1.1 ± 0.15 | 1.6 ± 0.09 |
| **Time of Peak (ms)** | 7.14 ± 0.27 | 11.52 ± 0.86 | 7.94 ± 0.57 |
| **Half-width (ms)** | 9.23 ± 0.75 | 18.18 ± 1.75 | 11.5 ± 1.27 |
| **Rise slope (dV/dt)** | 373.59 ± 49.91 | 94.2 ± 27.51 | 311.05 ± 40.1 |
| **Decay slope (dV/dt)** | -31.42 ± 4.08 | -10.7 ± 2.49 | -24.1 ± 2.9 |

### Table S2. All values and statistics for Figures 1 to 8

| FIGURE | | DESCRIPTION | VALUE | N NUMBER | STATISTICS | P VALUE | POST TEST | POST TEST P VALUE |
|---|---|---|---|---|---|---|---|---|
| 1 | A | Schematic illustration | N/A | N/A | N/A | N/A | N/A | N/A |
| | B | Representative fluorescence images (M1 cortex, GPe) | N/A | 15 slices / 4 mice | N/A | N/A | N/A | N/A |
| | | Representative fluorescence images (DLS) | N/A | 22 slices / 4 mice | N/A | N/A | N/A | N/A |
| | C | Representative immunostaining images | N/A | N/A | N/A | N/A | N/A | N/A |
| | D | RFP+ area (DL) | 19920 ± 1618 μm2 | 18 images / 3 mice | Ordinary one-way ANOVA | | Post hoc Holm-Sidak's test | |
| | | RFP+ area (VL) | 18920 ± 1507 μm2 | 18 images / 3 mice | Group effect, F(3, 68) = 1.086 | 0.3612 | Dorsolateral vs. Ventrolateral | 0.8719 |
| | | RFP+ area (DM) | 22350 ± 1271 μm2 | 18 images / 3 mice | | | Dorsolateral vs. Dorsomedial | 0.6966 |
| | | RFP+ area (VM) | 21650 ± 1603 μm2 | 18 images / 3 mice | | | Dorsolateral vs. Ventromedial | 0.8038 |
| | | | | | | | Ventrolateral vs. Dorsomedial | 0.5113 |
| | | | | | | | Ventrolateral vs. Ventromedial | 0.6821 |
| | | | | | | | Dorsomedial vs. Ventromedial | 0.8719 |
| | E | TH+ area (Lateral) | 632.7 ± 84.3 μm2 | 36 images / 6 mice | t(70) = 3.339 (Unpaired t-test) | 0.0013 | N/A | N/A |
| | | TH+ area (Medial) | 1190 ± 144.1 μm2 | 36 images / 6 mice | | | | |
| | | TH+ area (Dorsal) | 570.8 ± 98.9 μm2 | 36 images / 6 mice | t(70) = 4.251 (Unpaired t-test) | | N/A | N/A |
| | | TH+ area (Ventral) | 1252 ± 1216.1 μm2 | 36 images / 6 mice | | < 0.0001 | | |
| | F | TH+ area (DL) | 280 ± 36.6 μm2 | 18 images / 3 mice | Ordinary two-way ANOVA | | Post hoc Holm-Sidak's test | |
| | | TH+ area (VL) | 985.4 ± 115.4 μm2 | 18 images / 3 mice | DV axis effect, F(1, 68) = 21.23 | < 0.0001 | Dorsal:Lateral vs. Dorsal:Medial | 0.0208 |
| | | TH+ area (DM) | 861.7 ± 170.2 μm2 | 18 images / 3 mice | LM axis effect, F(1, 68) = 14.22 | 0.0003 | Dorsal:Lateral vs. Ventral:Lateral | 0.0061 |
| | | TH+ area (VM) | 1519 ± 209.3 μm2 | 18 images / 3 mice | Interaction, F(1, 68) = 14.22 | 0.8713 | Dorsal:Lateral vs. Ventral:Medial | <0.0001 |
| | | | | | | | Dorsal:Medial vs. Ventral:Lateral | 0.5561 |
| | | | | | | | Dorsal:Medial vs. Ventral:Medial | 0.0099 |
| | | | | | | | Ventral:Lateral vs. Ventral:Medial | 0.0258 |
| 2 | A | Schematic illustration | N/A | N/A | N/A | N/A | N/A | N/A |
| | B | Baseline (Normalized oIPSC) | 100% | 40 cells/ 26 mice | N/A | N/A | N/A | N/A |
| | | QP ON (Normalized oIPSC) | 50.60 ± 2.17% | 40 cells/ 26 mice | t(39) = 23.37 (one sample t-test) | < 0.0001 | | |
| | | QP OFF (Normalized oIPSC) | 38.99 ± 2.77% | 40 cells/ 26 mice | t(39) = 14.05 (one sample t-test) | < 0.0001 | | |
| | C | Baseline (PPR) | 1.44 ± 0.10 | 40 cells/ 26 mice | Repeated measures one-way ANOVA | | Post hoc Holm-Sidak's test | N/A |
| | | QP ON (PPR) | 1.89 ± 0.11 | 40 cells/ 26 mice | QP treatment effect, F (2, 78) = 13.68 | < 0.0001 | Baseline vs. QP ON | <0.0001 |
| | | QP OFF (PPR) | 1.69 ± 0.09 | 40 cells/ 26 mice | Individual effect, F (39, 78) = 5.943 | < 0.0001 | Baseline vs. QP OFF | 0.0087 |
| | | | | | | | QP ON vs. QP OFF | 0.0253 |
| | | Baseline (Normalized PPR) | 100% | 40 cells/ 26 mice | N/A | N/A | N/A | N/A |
| | | QP ON (Normalized PPR) | 143.10 ± 8.88% | 40 cells/ 26 mice | t(39) = 4.854 (one sample t-test) | < 0.0001 | | |
| | | QP OFF (Normalized PPR) | 129.80 ± 7.73% | 40 cells/ 26 mice | t(39) = 3.852 (one sample t-test) | 0.0004 | | |
| | D | Baseline (Normalized oIPSC) | 100% | 10 cells/ 9 mice | N/A | N/A | N/A | N/A |
| | | QP ON (Normalized oIPSC) | 54.38 ± 4.60% | 10 cells/ 9 mice | t(9) = 9.929 (one sample t-test) | < 0.0001 | | |
| | | QP OFF (Normalized oIPSC) | 39.33 ± 4.79% | 10 cells/ 9 mice | t(9) = 12.67 (one sample t-test) | < 0.0001 | | |
| | | Baseline (Normalized PPR) | 100% | 10 cells/ 9 mice | N/A | N/A | N/A | N/A |
| | | QP ON (Normalized PPR) | 184 ± 23.37% | 10 cells/ 9 mice | t(9) = 3.593 (one sample t-test) | 0.0058 | | |
| | | QP OFF (Normalized PPR) | 154.10 ± 14.71% | 10 cells/ 9 mice | t(9) = 3.680 (one sample t-test) | 0.0051 | | |
| | E | Baseline (Normalized oIPSC) | 100% | 10 cells/ 8 mice | N/A | N/A | N/A | N/A |
| | | QP ON (Normalized oIPSC) | 45.29 ± 4.53% | 10 cells/ 8 mice | t(9) = 12.07 (one sample t-test) | < 0.0001 | | |
| | | QP OFF (Normalized oIPSC) | 34.79 ± 5.23% | 10 cells/ 8 mice | t(9) = 12.48 (one sample t-test) | < 0.0001 | | |
| | | Baseline (Normalized PPR) | 100% | 10 cells/ 8 mice | N/A | N/A | N/A | N/A |
| | | QP ON (Normalized PPR) | 126.30 ± 15.15% | 10 cells/ 8 mice | t(9) = 1.736 (one sample t-test) | 0.1166 | | |
| | | QP OFF (Normalized PPR) | 135.30 ± 22.82% | 10 cells/ 8 mice | t(9) = 1.545 (one sample t-test) | 0.1567 | | |
| | F | Baseline (Normalized oIPSC) | 100% | 10 cells/ 8 mice | N/A | N/A | N/A | N/A |
| | | QP ON (Normalized oIPSC) | 47.49 ± 4.07% | 10 cells/ 8 mice | t(9) = 12.90 (one sample t-test) | < 0.0001 | | |
| | | QP OFF (Normalized oIPSC) | 35.84 ± 5.35% | 10 cells/ 8 mice | t(9) = 11.98 (one sample t-test) | < 0.0001 | | |
| | | Baseline (Normalized PPR) | 100% | 10 cells/ 8 mice | N/A | N/A | N/A | N/A |
| | | QP ON (Normalized PPR) | 115.90 ± 11.32% | 10 cells/ 8 mice | t(9) = 1.406 (one sample t-test) | 0.1934 | | |
| | | QP OFF (Normalized PPR) | 112.20 ± 10.24% | 10 cells/ 8 mice | t(9) = 1.187 (one sample t-test) | 0.2657 | | |
| | G | Baseline (Normalized oIPSC) | 100% | 10 cells/ 8 mice | N/A | N/A | N/A | N/A |
| | | QP ON (Normalized oIPSC) | 55.24 ± 3.85% | 10 cells/ 8 mice | t(9) = 9.929 (one sample t-test) | < 0.0001 | | |
| | | QP OFF (Normalized oIPSC) | 45.99 ± 6.78% | 10 cells/ 8 mice | t(9) = 12.67 (one sample t-test) | < 0.0001 | | |
| | | Baseline (Normalized PPR) | 100% | 10 cells/ 8 mice | N/A | N/A | N/A | N/A |
| | | QP ON (Normalized PPR) | 146.30 ± 12.77% | 10 cells/ 8 mice | t(9) = 3.624 (one sample t-test) | 0.0055 | | |
| | | QP OFF (Normalized PPR) | 117.50 ± 8.48% | 10 cells/ 8 mice | t(9) = 2.061 (one sample t-test) | 0.0694 | | |
| 3 | A | $Sr^{2+}$ (Amplitude) | 44.48 ± 6.68 pA | 8 cells/ 6 mice | t(7) = 3.070 (Paired t-test) | 0.0181 | N/A | N/A |
| | | $Sr^{2+}$ + QP (Amplitude) | 37.54 ± 5.32 pA | 8 cells/ 6 mice | | | | |
| | | $Sr^{2+}$ (Frequency) | 42.80 ± 9.12 Hz | 8 cells/ 6 mice | t(7) = 2.929 (Paired t-test) | 0.0221 | N/A | N/A |
| | | $Sr^{2+}$ + QP (Frequency) | 31.80 ± 8.30 Hz | 8 cells/ 6 mice | | | | |
| | B | $Sr^{2+}$ (Amplitude) | 52.31 ± 6.07 pA | 16 cells/ 15 mice | t(15) = 2.729 (Paired t-test) | 0.0155 | N/A | N/A |
| | | $Sr^{2+}$ + QP (Amplitude) | 47.91 ± 6.03 pA | 16 cells/ 15 mice | | | | |
| | | $Sr^{2+}$ (Frequency) | 41.03 ± 4.74 Hz | 16 cells/ 15 mice | t(15) = 0.5001 (Paired t-test) | 0.6242 | N/A | N/A |
| | | $Sr^{2+}$ + QP (Frequency) | 40.26 ± 4.35 Hz | 16 cells/ 15 mice | | | | |
| | C | $Sr^{2+}$ (Amplitude) | 52.84 ± 8.51 pA | 7 cells/ 7 mice | t(6) = 3.421 (Paired t-test) | 0.0141 | N/A | N/A |
| | | $Sr^{2+}$ + QP (Amplitude) | 45.05 ± 6.83 pA | 7 cells/ 7 mice | | | | |
| | | $Sr^{2+}$ (Frequency) | 46.85 ± 7.95 Hz | 7 cells/ 7 mice | t(6) = 1.054 (Paired t-test) | 0.3325 | N/A | N/A |
| | | $Sr^{2+}$ + QP (Frequency) | 43.29 ± 8.96 Hz | 7 cells/ 7 mice | | | | |
| | D | $Sr^{2+}$ (Amplitude) | 42.62 ± 3.15 pA | 9 cells/ 8 mice | t(8) = 5.311 (Paired t-test) | 0.0007 | N/A | N/A |
| | | $Sr^{2+}$ + QP (Amplitude) | 32.38 ± 2.64 pA | 9 cells/ 8 mice | | | | |
| | | $Sr^{2+}$ (Frequency) | 54.32 ± 5.36 Hz | 9 cells/ 8 mice | t(8) = 5.031 (Paired t-test) | 0.001 | N/A | N/A |
| | | $Sr^{2+}$ + QP (Frequency) | 28.63 ± 2.38 Hz | 9 cells/ 8 mice | | | | |

| | | | | | | | | |
|---|---|---|---|---|---|---|---|---|
| | E | Cumulative plots | N/A | N/A | N/A | N/A | N/A | N/A |
| | F | Schematic illustration | N/A | N/A | N/A | N/A | N/A | N/A |
| | G | L741626 (Normalized oIPSC) | 100% | 7 cells/ 5 mice | N/A | N/A | N/A | N/A |
| | | L741626 + QP (Normalized oIPSC) | 56.04 ± 5.73% | 7 cells/ 5 mice | t(6) = 7.678 (one sample t-test) | 0.0003 | | |
| | | L741626 (Normalized PPR) | 100% | 7 cells/ 5 mice | N/A | N/A | N/A | N/A |
| | | L741626 + QP (Normalized PPR) | 111.70 ± 12.13% | 7 cells/ 5 mice | t(6) = 0.9653 (one sample t-test) | 0.3717 | | |
| | H | Schematic illustration | N/A | N/A | N/A | N/A | N/A | N/A |
| | I | L741626 (Normalized oIPSC) | 100% | 7 cells/ 4 mice | N/A | N/A | N/A | N/A |
| | | L741626 + QP (Normalized oIPSC) | 61.37 ± 8.55% | 7 cells/ 4 mice | t(6) = 4.518 (one sample t-test) | 0.004 | | |
| | | L741626 (Normalized PPR) | 100% | 7 cells/ 4 mice | N/A | N/A | N/A | N/A |
| | | L741626 + QP (Normalized PPR) | 106.8 ± 8.90% | 7 cells/ 4 mice | t(6) = 0.7609 (one sample t-test) | 0.4755 | | |
| 4 | A | Schematic illustration | N/A | N/A | N/A | N/A | N/A | N/A |
| | B | Schematic illustration | N/A | N/A | N/A | N/A | N/A | N/A |
| | C | Representative immunostaining images | N/A | N/A | N/A | N/A | N/A | N/A |
| | D | D2R+ area (DL,VM GPe) | 109.70 ± 7.91 μm2 | 95 images/ 9 mice | t(188) = 3.133 (Unpaired t-test) | 0.002 | N/A | N/A |
| | | D2R+ area (VL,DM GPe) | 80.39 ± 4.99 μm2 | 95 images/ 9 mice | | | | |
| | | D2R+RFP+ area (DL,VM GPe) | 15.61 ± 1.52 μm2 | 95 images/ 9 mice | t(186) = 2.705 (Unpaired t-test) | 0.0075 | N/A | N/A |
| | | D2R+RFP+ area (VL,DM GPe) | 10.80 ± 0.90 μm2 | 93 images/ 9 mice | | | | |
| | | D2R+GFP+ area (DL,VM GPe) | 21.96 ± 2.80 μm2 | 76 images/ 9 mice | t(182) = 2.426 (Unpaired t-test) | 0.0163 | N/A | N/A |
| | | D2R+GFP+ area (VL,DM GPe) | 14.51 ± 1.66 μm2 | 108 images/ 9 mice | | | | |
| | | D2R+GFP+ or D2R+RFP+ area (DL,VM GPe) | 29.78 ± 3.41 μm2 | 92 images/ 9 mice | t(193) = 2.052 (Unpaired t-test) | 0.0415 | N/A | N/A |
| | | D2R+GFP+ or D2R+RFP+ area (VL,DM GPe) | 21.41 ± 2.36 μm2 | 103 images/ 9 mice | | | | |
| | | GFP+ area (DL,VM GPe) | 163.60 ± 14.08 μm2 | 96 images/ 9 mice | t(193) = 1.173 (Unpaired t-test) | 0.2423 | N/A | N/A |
| | | GFP+ area (VL,DM GPe) | 187.40 ± 14.62 μm2 | 99 images/ 9 mice | | | | |
| | E | Cumulative plots | N/A | N/A | N/A | N/A | N/A | N/A |
| | F | Schematic illustration | N/A | N/A | N/A | N/A | N/A | N/A |
| | G | Schematic illustration | N/A | N/A | N/A | N/A | N/A | N/A |
| | H | Representative immunostaining images | N/A | N/A | N/A | N/A | N/A | N/A |
| | I | D4R+ volume (DL,VM GPe) | 14.58 ± 1.32 μm3 | 47 images/ 6 mice | t(97) = 3.029 (Unpaired t-test) | 0.0031 | N/A | N/A |
| | | D4R+ volume (VL,DM GPe) | 29.06 ± 1.84 μm3 | 52 images/ 6 mice | | | | |
| | | The number of D4R+ fluorescence (DL,VM GPe) | 317.80 ± 30.08 | 47 images/ 6 mice | t(97) = 4.705 (Unpaired t-test) | < 0.0001 | N/A | N/A |
| | | The number of D4R+ fluorescence (VL,DM GPe) | 534.10 ± 32.20 | 52 images/ 6 mice | | | | |
| | | GABA$_A$R+ volume (DL,VM GPe) | 533.50 ± 29.48 μm3 | 47 images/ 6 mice | t(97) = 0.07017 (Unpaired t-test) | 0.9442 | N/A | N/A |
| | | GABA$_A$R+ volume (VL,DM GPe) | 536.60 ± 31.58 μm3 | 52 images/ 6 mice | | | | |
| | J | The number of GABA$_A$R+ fluorescence (DL,VM GPe) | 2686.30 ± 169.60 | 47 images/ 6 mice | t(97) = 0.002606 (Unpaired t-test) | 0.9979 | N/A | N/A |
| | | The number of GABA$_A$R+ fluorescence (VL,DM GPe) | 2685.80 ± 143.30 | 52 images/ 6 mice | | | | |
| | K | Cumulative plots | N/A | N/A | N/A | N/A | N/A | N/A |
| | A | Schematic illustration | N/A | N/A | N/A | N/A | N/A | N/A |
| | B | Baseline oIPSC (0 sec) | 100% | 15 cells/ 13 mice | Repeated measures two-way ANOVA | | Post hoc Holm-Sidak's test | |
| | | Baseline oIPSC (0.05 sec) | 81.10 ± 9.90% | 15 cells/ 13 mice | Pulse train effect, F (9, 126) = 31.42 | < 0.0001 | Baseline vs Scaled QP oIPSC (0 sec) | >0.9999 |
| | | Baseline oIPSC (0.10 sec) | 53.63 ± 9.30% | 15 cells/ 13 mice | QP treatment effect, F (1, 14) = 26.60 | 0.0001 | Baseline vs Scaled QP oIPSC (0.05 sec) | <0.0001 |
| | | Baseline oIPSC (0.15 sec) | 40.39 ± 7.81% | 15 cells/ 13 mice | Interaction, F (9, 126) = 4.639 | < 0.0001 | Baseline vs Scaled QP oIPSC (0.10 sec) | <0.0001 |
| | | Baseline oIPSC (0.20 sec) | 35.77 ± 7.97% | 15 cells/ 13 mice | | | Baseline vs Scaled QP oIPSC (0.15 sec) | <0.0001 |
| | | Baseline oIPSC (0.25 sec) | 32.74 ± 8.83% | 15 cells/ 13 mice | | | Baseline vs Scaled QP oIPSC (0.20 sec) | 0.0147 |
| | | Baseline oIPSC (0.30 sec) | 24.38 ± 5.45% | 15 cells/ 13 mice | | | Baseline vs Scaled QP oIPSC (0.25 sec) | 0.0147 |
| | | Baseline oIPSC (0.35 sec) | 26.99 ± 6.60% | 15 cells/ 13 mice | | | Baseline vs Scaled QP oIPSC (0.30 sec) | <0.0001 |
| | | Baseline oIPSC (0.40 sec) | 31.73 ± 11.77% | 15 cells/ 13 mice | | | Baseline vs Scaled QP oIPSC (0.35 sec) | 0.0005 |
| | | Baseline oIPSC (0.45 sec) | 25.84 ± 6.50% | 15 cells/ 13 mice | | | Baseline vs Scaled QP oIPSC (0.40 sec) | 0.0012 |
| | | Scaled QP oIPSC (0 sec) | 100% | 15 cells/ 13 mice | | | Baseline vs Scaled QP oIPSC (0.45 sec) | 0.0002 |
| | | Scaled QP oIPSC (0.05 sec) | 131.19 ± 16.00% | 15 cells/ 13 mice | | | | |
| | | Scaled QP oIPSC (0.10 sec) | 88.91 ± 11.67% | 15 cells/ 13 mice | | | | |
| | | Scaled QP oIPSC (0.15 sec) | 67.76 ± 10.01% | 15 cells/ 13 mice | | | | |
| | | Scaled QP oIPSC (0.20 sec) | 52.57 ± 7.88% | 15 cells/ 13 mice | | | | |
| | | Scaled QP oIPSC (0.25 sec) | 49.96 ± 7.89% | 15 cells/ 13 mice | | | | |
| | | Scaled QP oIPSC (0.30 sec) | 52.64 ± 12.30% | 15 cells/ 13 mice | | | | |
| | | Scaled QP oIPSC (0.35 sec) | 51.09 ± 10.98% | 15 cells/ 13 mice | | | | |
| | | Scaled QP oIPSC (0.40 sec) | 54.16 ± 11.47% | 15 cells/ 13 mice | | | | |
| | | Scaled QP oIPSC (0.45 sec) | 51.96 ± 10.14% | 15 cells/ 13 mice | | | | |
| | C | Baseline oIPSC (P2/P1) | 0.795 ± 0.089 | 15 cells/ 13 mice | Repeated measures two-way ANOVA | | Post hoc Holm-Sidak's test | |
| | | Baseline oIPSC (P10/P1) | 0.258 ± 0.065 | 15 cells/ 13 mice | QP treatment effect, F (1, 14) = 23.52 | 0.0003 | P2/P1 - P10/P1 | |
| | | QP oIPSC (P2/P1) | 1.259 ± 0.149 | 15 cells/ 13 mice | Peak ratio effect, F (1, 14) = 48.16 | < 0.0001 | Baseline | <0.0001 |
| | | QP oIPSC (P10/P1) | 0.520 ± 0.101 | 15 cells/ 13 mice | Interaction, F (1, 14) = 4.015 | 0.0648 | QP(10uM) | <0.0001 |
| | | | | | | | Baseline - QP(10uM) | |
| | | | | | | | P2/P1 | <0.0001 |
| | | | | | | | P10/P1 | 0.0026 |
| | D | Schematic illustration | N/A | N/A | N/A | N/A | N/A | N/A |
| | E | Baseline oIPSC (0 sec) | 100.00 ± 0.00% | 15 cells/ 13 mice | Repeated measures two-way ANOVA | | Post hoc Holm-Sidak's test | |
| | | Baseline oIPSC (0.05 sec) | 103.70 ± 15.91% | 15 cells/ 13 mice | Pulse train effect, F (9, 126) = 18.31 | < 0.0001 | Baseline vs Scaled QP oIPSC (0 sec) | >0.9999 |
| | | Baseline oIPSC (0.10 sec) | 82.71 ± 19.06% | 15 cells/ 13 mice | QP treatment effect, F (1, 14) = 2.724 | 0.1211 | Baseline vs Scaled QP oIPSC (0.05 sec) | 0.9964 |
| | | Baseline oIPSC (0.15 sec) | 57.37 ± 15.80% | 15 cells/ 13 mice | Interaction, F (9, 126) = 0.8486 | 0.573 | Baseline vs Scaled QP oIPSC (0.10 sec) | 0.4109 |
| | | Baseline oIPSC (0.20 sec) | 48.38 ± 15.34% | 15 cells/ 13 mice | | | Baseline vs Scaled QP oIPSC (0.15 sec) | 0.864 |
| | | Baseline oIPSC (0.25 sec) | 38.92 ± 11.41% | 15 cells/ 13 mice | | | Baseline vs Scaled QP oIPSC (0.20 sec) | 0.8571 |
| | | Baseline oIPSC (0.30 sec) | 30.65 ± 9.27% | 15 cells/ 13 mice | | | Baseline vs Scaled QP oIPSC (0.25 sec) | 0.0709 |
| | | Baseline oIPSC (0.35 sec) | 25.47 ± 7.32% | 15 cells/ 13 mice | | | Baseline vs Scaled QP oIPSC (0.30 sec) | 0.9313 |
| | | Baseline oIPSC (0.40 sec) | 21.02 ± 5.24% | 15 cells/ 13 mice | | | Baseline vs Scaled QP oIPSC (0.35 sec) | 0.9964 |
| | | Baseline oIPSC (0.45 sec) | 14.89 ± 2.80% | 15 cells/ 13 mice | | | Baseline vs Scaled QP oIPSC (0.40 sec) | 0.864 |
| | | Scaled QP oIPSC (0 sec) | 100.00 ± 0.00% | 15 cells/ 13 mice | | | Baseline vs Scaled QP oIPSC (0.45 sec) | 0.864 |
| | | Scaled QP oIPSC (0.05 sec) | 102.28 ± 11.26% | 15 cells/ 13 mice | | | | |

| | | | | | | | | |
|---|---|---|---|---|---|---|---|---|
| | | Scaled QP oIPSC (0.10 sec) | 96.91 ± 16.72% | 15 cells/ 13 mice | | | | |
| | | Scaled QP oIPSC (0.15 sec) | 65.95 ± 16.61% | 15 cells/ 13 mice | | | | |
| | | Scaled QP oIPSC (0.20 sec) | 57.57 ± 17.76% | 15 cells/ 13 mice | | | | |
| | | Scaled QP oIPSC (0.25 sec) | 59.08 ± 18.59% | 15 cells/ 13 mice | | | | |
| | | Scaled QP oIPSC (0.30 sec) | 35.79 ± 8.77% | 15 cells/ 13 mice | | | | |
| | | Scaled QP oIPSC (0.35 sec) | 24.76 ± 4.76% | 15 cells/ 13 mice | | | | |
| | | Scaled QP oIPSC (0.40 sec) | 29.07 ± 6.20% | 15 cells/ 13 mice | | | | |
| | | Scaled QP oIPSC (0.45 sec) | 23.18 ± 5.08% | 15 cells/ 13 mice | | | | |
| | F | Baseline oIPSC (P2/P1) | 1.037 ± 0.159 | 15 cells/ 13 mice | Repeated measures two-way ANOVA | | Post hoc Holm-Sidak's test | |
| | | Baseline oIPSC (P10/P1) | 0.149 ± 0.028 | 15 cells/ 13 mice | QP treatment effect, F (1, 14) = 0.4500 | 0.5132 | P2/P1 - P10/P1 | |
| | | QP oIPSC (P2/P1) | 1.023 ± 0.113 | 15 cells/ 13 mice | Peak ratio effect, F (1, 14) = 47.37 | < 0.0001 | Baseline | <0.0001 |
| | | QP oIPSC (P10/P1) | 0.232 ± 0.051 | 15 cells/ 13 mice | Interaction, F (1, 14) = 1.189 | 0.2939 | QP(10uM) | <0.0001 |
| | | | | | | | Baseline - QP(10uM) | |
| | | | | | | | P2/P1 | 0.824 |
| | | | | | | | P10/P1 | 0.3749 |
| | G | Schematic illustration | | N/A | N/A | N/A | N/A | N/A |
| | H | Baseline oIPSC (0 sec) | 100.00 ± 0.00% | 16 cells/ 14 mice | Repeated measures two-way ANOVA | | Post hoc Holm-Sidak's test | |
| | | Baseline oIPSC (0.05 sec) | 165.43 ± 7.47% | 16 cells/ 14 mice | Pulse train effect, F (9, 135) = 4.607 | < 0.0001 | Baseline vs Scaled QP oIPSC (0 sec) | >0.9999 |
| | | Baseline oIPSC (0.10 sec) | 188.33 ± 12.19% | 16 cells/ 14 mice | QP treatment effect, F (1, 15) = 0.2218 | 0.6445 | Baseline vs Scaled QP oIPSC (0.05 sec) | 0.9749 |
| | | Baseline oIPSC (0.15 sec) | 190.52 ± 17.69% | 16 cells/ 14 mice | Interaction, F (9, 135) = 1.125 | 0.3497 | Baseline vs Scaled QP oIPSC (0.10 sec) | 0.1195 |
| | | Baseline oIPSC (0.20 sec) | 178.87 ± 21.44% | 16 cells/ 14 mice | | | Baseline vs Scaled QP oIPSC (0.15 sec) | 0.9656 |
| | | Baseline oIPSC (0.25 sec) | 165.80 ± 22.42% | 16 cells/ 14 mice | | | Baseline vs Scaled QP oIPSC (0.20 sec) | 0.2537 |
| | | Baseline oIPSC (0.30 sec) | 159.53 ± 23.76% | 16 cells/ 14 mice | | | Baseline vs Scaled QP oIPSC (0.25 sec) | 0.9696 |
| | | Baseline oIPSC (0.35 sec) | 158.96 ± 28.00% | 16 cells/ 14 mice | | | Baseline vs Scaled QP oIPSC (0.30 sec) | 0.9749 |
| | | Baseline oIPSC (0.40 sec) | 152.83 ± 30.35% | 16 cells/ 14 mice | | | Baseline vs Scaled QP oIPSC (0.35 sec) | 0.9658 |
| | | Baseline oIPSC (0.45 sec) | 146.68 ± 31.74% | 16 cells/ 14 mice | | | Baseline vs Scaled QP oIPSC (0.40 sec) | 0.9749 |
| 5 | | Scaled QP oIPSC (0 sec) | 100.00 ± 0.00% | 16 cells/ 14 mice | | | Baseline vs Scaled QP oIPSC (0.45 sec) | 0.9437 |
| | | Scaled QP oIPSC (0.05 sec) | 161.76 ± 10.79% | 16 cells/ 14 mice | | | | |
| | | Scaled QP oIPSC (0.10 sec) | 209.78 ± 21.78% | 16 cells/ 14 mice | | | | |
| | | Scaled QP oIPSC (0.15 sec) | 183.08 ± 21.99% | 16 cells/ 14 mice | | | | |
| | | Scaled QP oIPSC (0.20 sec) | 197.26 ± 21.36% | 16 cells/ 14 mice | | | | |
| | | Scaled QP oIPSC (0.25 sec) | 171.50 ± 22.91% | 16 cells/ 14 mice | | | | |
| | | Scaled QP oIPSC (0.30 sec) | 161.37 ± 25.07% | 16 cells/ 14 mice | | | | |
| | | Scaled QP oIPSC (0.35 sec) | 165.67 ± 27.88% | 16 cells/ 14 mice | | | | |
| | | Scaled QP oIPSC (0.40 sec) | 157.26 ± 32.40% | 16 cells/ 14 mice | | | | |
| | | Scaled QP oIPSC (0.45 sec) | 155.47 ± 33.47% | 16 cells/ 14 mice | | | | |
| | I | Baseline oIPSC (P2/P1) | 1.654 ± 0.075 | 16 cells/ 14 mice | Repeated measures two-way ANOVA | | Post hoc Holm-Sidak's test | |
| | | Baseline oIPSC (P10/P1) | 1.467 ± 0.317 | 16 cells/ 14 mice | QP treatment effect, F (1, 15) = 0.06557 | 0.8014 | P2/P1 - P10/P1 | |
| | | QP oIPSC (P2/P1) | 1.618 ± 0.108 | 16 cells/ 14 mice | Peak ratio effect, F (1, 15) = 0.1687 | 0.6871 | Baseline | 0.1466 |
| | | QP oIPSC (P10/P1) | 1.555 ± 0.335 | 16 cells/ 14 mice | Interaction, F (1, 15) = 0.8001 | 0.3852 | QP(10uM) | 0.781 |
| | | | | | | | Baseline - QP(10uM) | |
| | | | | | | | P2/P1 | 0.7151 |
| | | | | | | | P10/P1 | 0.6229 |
| | J | Schematic illustration | | N/A | N/A | N/A | N/A | N/A |
| | K | Baseline oIPSC (0 sec) | 100.00 ± 0.00% | 17 cells/ 15 mice | Repeated measures two-way ANOVA | | Post hoc Holm-Sidak's test | |
| | | Baseline oIPSC (0.05 sec) | 152.01 ± 10.32% | 17 cells/ 15 mice | Pulse train effect, F (9, 144) = 19.30 | < 0.0001 | Baseline vs Scaled QP oIPSC (0 sec) | >0.9999 |
| | | Baseline oIPSC (0.10 sec) | 167.70 ± 15.34% | 17 cells/ 15 mice | QP treatment effect, F (1, 16) = 13.37 | 0.0021 | Baseline vs Scaled QP oIPSC (0.05 sec) | 0.001 |
| | | Baseline oIPSC (0.15 sec) | 159.77 ± 13.98% | 17 cells/ 15 mice | Interaction,F (9, 144) = 4.609 | < 0.0001 | Baseline vs Scaled QP oIPSC (0.10 sec) | <0.0001 |
| | | Baseline oIPSC (0.20 sec) | 134.11 ± 12.43% | 17 cells/ 15 mice | | | Baseline vs Scaled QP oIPSC (0.15 sec) | <0.0001 |
| | | Baseline oIPSC (0.25 sec) | 119.70 ± 13.64% | 17 cells/ 15 mice | | | Baseline vs Scaled QP oIPSC (0.20 sec) | <0.0001 |
| | | Baseline oIPSC (0.30 sec) | 96.48 ± 12.29% | 17 cells/ 15 mice | | | Baseline vs Scaled QP oIPSC (0.25 sec) | 0.0006 |
| | | Baseline oIPSC (0.35 sec) | 79.90 ± 12.00% | 17 cells/ 15 mice | | | Baseline vs Scaled QP oIPSC (0.30 sec) | 0.0274 |
| | | Baseline oIPSC (0.40 sec) | 69.39 ± 13.79% | 17 cells/ 15 mice | | | Baseline vs Scaled QP oIPSC (0.35 sec) | 0.0038 |
| | | Baseline oIPSC (0.45 sec) | 60.09 ± 12.49% | 17 cells/ 15 mice | | | Baseline vs Scaled QP oIPSC (0.40 sec) | 0.0027 |
| | | Scaled QP oIPSC (0 sec) | 100.00 ± 0.00% | 17 cells/ 15 mice | | | Baseline vs Scaled QP oIPSC (0.45 sec) | 0.0274 |
| | | Scaled QP oIPSC (0.05 sec) | 199.06 ± 13.45% | 17 cells/ 15 mice | | | | |
| | | Scaled QP oIPSC (0.10 sec) | 253.46 ± 33.42% | 17 cells/ 15 mice | | | | |
| | | Scaled QP oIPSC (0.15 sec) | 243.57 ± 37.74% | 17 cells/ 15 mice | | | | |
| | | Scaled QP oIPSC (0.20 sec) | 204.30 ± 26.10% | 17 cells/ 15 mice | | | | |
| | | Scaled QP oIPSC (0.25 sec) | 168.62 ± 20.64% | 17 cells/ 15 mice | | | | |
| | | Scaled QP oIPSC (0.30 sec) | 128.56 ± 16.88% | 17 cells/ 15 mice | | | | |
| | | Scaled QP oIPSC (0.35 sec) | 120.95 ± 21.55% | 17 cells/ 15 mice | | | | |
| | | Scaled QP oIPSC (0.40 sec) | 112.36 ± 19.90% | 17 cells/ 15 mice | | | | |
| | | Scaled QP oIPSC (0.45 sec) | 90.87 ± 16.98% | 17 cells/ 15 mice | | | | |
| | L | Baseline oIPSC (P2/P1) | 1.520 ± 0.103 | 17 cells/ 15 mice | Repeated measures two-way ANOVA | | Post hoc Holm-Sidak's test | |
| | | Baseline oIPSC (P10/P1) | 0.601 ± 0.125 | 17 cells/ 15 mice | QP treatment effect, F (1, 16) = 31.09 | < 0.0001 | P2/P1 - P10/P1 | |
| | | QP oIPSC (P2/P1) | 1.991 ± 0.135 | 17 cells/ 15 mice | Peak ratio effect, F (1, 16) = 28.65 | < 0.0001 | Baseline | <0.0001 |
| | | QP oIPSC (P10/P1) | 0.909 ± 0.170 | 17 cells/ 15 mice | Interaction, F (1, 16) = 2.160 | 0.1611 | QP(10uM) | <0.0001 |
| | | | | | | | Baseline - QP(10uM) | |
| | | | | | | | P2/P1 | <0.0001 |
| | | | | | | | P10/P1 | 0.0012 |
| | | Baseline P10/P1 (DL) | 0.2584 ± 0.0650 | 15 cells/ 13 mice | Ordinary one-way ANOVA | | Post hoc Holm-Sidak's test | |
| | | Baseline P10/P1 (VL) | 0.1489 ± 0.0280 | 15 cells/ 13 mice | Group effect, F (3, 59) = 11.18 | < 0.0001 | DL vs. VL | 0.6728 |
| | | Baseline P10/P1 (DM) | 1.4670 ± 0.3174 | 16 cells/ 14 mice | | | DL vs. DM | <0.0001 |
| | | Baseline P10/P1 (VM) | 0.6009 ± 0.1249 | 17 cells/ 15 mice | | | DL vs. VM | 0.3219 |
| | | | | | | | VL vs. DM | <0.0001 |
| | | | | | | | VL vs. VM | 0.2114 |
| | | | | | | | DM vs. VM | 0.0034 |

| | | | | | | | | |
|---|---|---|---|---|---|---|---|---|
| | M | Baseline P10/P1 (DL) | 0.258 ± 0.065 | 15 cells/ 13 mice | Repeated measures two-way ANOVA | | Post hoc Holm-Sidak's test | |
| | | QP P10/P1 (DL) | 0.520 ± 0.101 | 15 cells/ 13 mice | QP treatment effect, F (1, 59) = 14.14 | 0.2442 | Baseline - QP(10uM) | |
| | | Baseline P10/P1 (VL) | 0.149 ± 0.028 | 15 cells/ 13 mice | GPe subregion effect, F (3, 59) = 10.00 | 0.0004 | DL | 0.0353 |
| | | QP P10/P1 (VL) | 0.232 ± 0.051 | 15 cells/ 13 mice | Interaction, F (3, 59) = 1.426 | < 0.0001 | VL | 0.604 |
| | | Baseline P10/P1 (DM) | 1.467 ± 0.317 | 16 cells/ 14 mice | | | DM | 0.604 |
| | | QP P10/P1 (DM) | 1.555 ± 0.335 | 16 cells/ 14 mice | | | VM | 0.0075 |
| | | Baseline P10/P1 (VM) | 0.601 ± 0.125 | 17 cells/ 15 mice | | | | |
| | | QP P10/P1 (VM) | 0.909 ± 0.170 | 17 cells/ 15 mice | | | | |
| | N | Charge ratio (DL) | 1.2260 ± 0.0914 | 15 cells/ 13 mice | Ordinary one-way ANOVA | | Post hoc Holm-Sidak's test | |
| | | Charge ratio (VL) | 0.9444 ± 0.0654 | 15 cells/ 13 mice | Group effect, F (3, 59) = 2.972 | 0.0389 | DL vs. VL | 0.0604 |
| | | Charge ratio (DM) | 1.0430 ± 0.0767 | 16 cells/ 14 mice | | | DL vs. DM | 0.2982 |
| | | Charge ratio (VM) | 1.1770 ± 0.0586 | 17 cells/ 15 mice | | | DL vs. VM | 0.6375 |
| | | | | | | | VL vs. DM | 0.5797 |
| | | | | | | | VL vs. VM | 0.131 |
| | | | | | | | DM vs. VM | 0.4675 |
| | | Charge ratio (DL,VM GPe) | 1.20 ± 0.0522 | 32 cells/ 28 mice | t(61) = 2.815 (Unpaired t-test) | 0.0066 | N/A | N/A |
| | | Charge ratio (VL,DM GPe) | 0.9951 ± 0.0506 | 31 cells/ 27 mice | | | | |
| 6 | A | Schematic illustration | N/A | N/A | N/A | N/A | N/A | N/A |
| | B | Baseline (Normalized oIPSC) | 100% | 10 cells/ 10 mice | N/A | N/A | N/A | N/A |
| | | QP ON (Normalized oIPSC) | 49.95 ± 7.23% | 10 cells/ 10 mice | t(9) = 6.926 (one sample t-test) | < 0.0001 | | |
| | | QP OFF (Normalized oIPSC) | 31.65 ± 5.41% | 10 cells/ 10 mice | t(9) = 12.63 (one sample t-test) | < 0.0001 | | |
| | | Baseline (Normalized PPR) | 100% | 10 cells/ 10 mice | N/A | N/A | N/A | N/A |
| | | QP ON (Normalized PPR) | 132.70 ± 15.88% | 10 cells/ 10 mice | t(9) = 2.061 (one sample t-test) | 0.0693 | | |
| | | QP OFF (Normalized PPR) | 117.40 ± 13.59% | 10 cells/ 10 mice | t(9) = 1.278 (one sample t-test) | 0.2332 | | |
| | C | Baseline (Normalized oIPSC) | 100% | 12 cells/ 12 mice | N/A | N/A | N/A | N/A |
| | | QP ON (Normalized oIPSC) | 52.51 ± 5.44% | 12 cells/ 12 mice | t(11) = 8.734 (one sample t-test) | < 0.0001 | | |
| | | QP OFF (Normalized oIPSC) | 41.80 ± 7.14% | 12 cells/ 12 mice | t(11) = 8.150 (one sample t-test) | < 0.0001 | | |
| | | Baseline (Normalized PPR) | 100% | 12 cells/ 12 mice | N/A | N/A | N/A | N/A |
| | | QP ON (Normalized PPR) | 153.30 ± 12.68% | 12 cells/ 12 mice | t(11) = 4.208 (one sample t-test) | 0.0015 | | |
| | | QP OFF (Normalized PPR) | 136.0 ± 10.86% | 12 cells/ 12 mice | t(11) = 3.312 (one sample t-test) | 0.0069 | | |
| | D | Baseline (Normalized oIPSC) | 100% | 10 cells/ 9 mice | N/A | N/A | N/A | N/A |
| | | QP ON (Normalized oIPSC) | 48.86 ± 8.03% | 10 cells/ 9 mice | t(9) = 6.369 (one sample t-test) | 0.0001 | | |
| | | QP OFF (Normalized oIPSC) | 31.79 ± 6.97% | 10 cells/ 9 mice | t(9) = 9.790 (one sample t-test) | < 0.0001 | | |
| | | Baseline (Normalized PPR) | 100% | 10 cells/ 9 mice | N/A | N/A | N/A | N/A |
| | | QP ON (Normalized PPR) | 132.50 ± 9.07% | 10 cells/ 9 mice | t(9) = 3.584 (one sample t-test) | 0.0059 | | |
| | | QP OFF (Normalized PPR) | 142.50 ± 7.04% | 10 cells/ 9 mice | t(9) = 6.033 (one sample t-test) | 0.0002 | | |
| | E | Baseline (Normalized oIPSC) | 100% | 18 cells/ 16 mice | N/A | N/A | N/A | N/A |
| | | QP ON (Normalized oIPSC) | 56.77 ± 3.77% | 18 cells/ 16 mice | t(17) = 11.47 (one sample t-test) | < 0.0001 | | |
| | | QP OFF (Normalized oIPSC) | 41.40 ± 4.70% | 18 cells/ 16 mice | t(17) = 12.47 (one sample t-test) | < 0.0001 | | |
| | | Baseline (Normalized PPR) | 100% | 18 cells/ 16 mice | N/A | N/A | N/A | N/A |
| | | QP ON (Normalized PPR) | 122.0 ± 6.69% | 18 cells/ 16 mice | t(17) = 3.293 (one sample t-test) | 0.0043 | | |
| | | QP OFF (Normalized PPR) | 111.70 ± 5.74% | 18 cells/ 16 mice | t(17) = 2.046 (one sample t-test) | 0.0566 | | |
| | F | oIPSC DL (Contra) | 44.347 ± 5.088% | 10 cells/ 10 mice | Ordinary two-way ANOVA | | Post hoc Holm-Sidak's test | |
| | | oIPSC DL (Ipsi) | 49.954 ± 7.226% | 10 cells/ 10 mice | GPe subregoin effect, F (3, 87) = 0.7165 | 0.5448 | Control - 6-OHDA | |
| | | oIPSC VL (Contra) | 51.447 ± 4.930% | 10 cells/ 9 mice | 6-OHDA effect, F (1, 87) = 0.2493 | 0.6188 | DL | 0.8661 |
| | | oIPSC VL (Ipsi) | 48.861 ± 8.030% | 10 cells/ 9 mice | Interaction, F (3, 87) = 0.3151 | 0.8144 | VL | 0.9391 |
| | | oIPSC DM (Contra) | 53.383 ± 4.646% | 12 cells/ 12 mice | | | DM | 0.9391 |
| | | oIPSC DM (Ipsi) | 52.514 ± 5.437% | 12 cells/ 12 mice | | | VM | 0.8661 |
| | | oIPSC VM (Contra) | 51.443 ± 3.434% | 18 cells/ 16 mice | | | | |
| | | oIPSC VM (Ipsi) | 56.773 ± 3.770% | 18 cells/ 16 mice | | | | |
| | | PPR DL (Contra) | 139.859 ± 9.380% | 10 cells/ 10 mice | Ordinary two-way ANOVA | | Post hoc Holm-Sidak's test | |
| | | PPR DL (Ipsi) | 132.732 ± 15.879% | 10 cells/ 10 mice | GPe subregoin effect, F (3, 87) = 0.6508 | 0.5846 | Control - 6-OHDA | |
| | | PPR VL (Contra) | 108.932 ± 12.277% | 10 cells/ 9 mice | 6-OHDA effect, F (1, 87) = 4.062 | 0.047 | DL | 0.6677 |
| | | PPR VL (Ipsi) | 132.508 ± 9.071% | 10 cells/ 9 mice | Interaction, F (3, 87) = 4.725 | 0.0042 | VL | 0.4262 |
| | | PPR DM (Contra) | 95.106 ± 13.944% | 12 cells/ 12 mice | | | DM | 0.0012 |
| | | PPR DM (Ipsi) | 153.346 ± 12.678% | 12 cells/ 12 mice | | | VM | 0.5868 |
| | | PPR VM (Contra) | 134.012 ± 8.134% | 18 cells/ 16 mice | | | | |
| | | PPR VM (Ipsi) | 122.040 ± 6.694% | 18 cells/ 16 mice | | | | |
| | G | oIPSC DL (Contra) | 26.774 ± 6.324% | 10 cells/ 10 mice | Ordinary two-way ANOVA | | Post hoc Holm-Sidak's test | |
| | | oIPSC DL (Ipsi) | 31.651 ± 5.413% | 10 cells/ 10 mice | GPe subregoin effect, F (3, 87) = 1.649 | 0.184 | Control - 6-OHDA | |
| | | oIPSC VL (Contra) | 37.657 ± 5.798% | 10 cells/ 9 mice | 6-OHDA effect,F (1, 87) = 0.2811 | 0.5974 | DL | 0.8769 |
| | | oIPSC VL (Ipsi) | 31.790 ± 6.968% | 10 cells/ 9 mice | Interaction,F (3, 87) = 0.4742 | 0.701 | VL | 0.8769 |
| | | oIPSC DM (Contra) | 34.370 ± 2.790% | 12 cells/ 12 mice | | | DM | 0.8079 |
| | | oIPSC DM (Ipsi) | 41.805 ± 7.140% | 12 cells/ 12 mice | | | VM | 0.8769 |
| | | oIPSC VM (Contra) | 39.535 ± 3.705% | 18 cells/ 16 mice | | | | |
| | | oIPSC VM (Ipsi) | 41.398 ± 4.70% | 18 cells/ 16 mice | | | | |
| | | PPR DL (Contra) | 147.0 ± 14.870% | 10 cells/ 10 mice | Ordinary two-way ANOVA | | Post hoc Holm-Sidak's test | |
| | | PPR DL (Ipsi) | 117.371 ± 13.592% | 10 cells/ 10 mice | GPe subregoin effect, F (3, 87) = 1.159 | 0.3303 | Control - 6-OHDA | |
| | | PPR VL (Contra) | 99.991 ± 14.058 | 10 cells/ 9 mice | 6-OHDA effect, F (1, 87) = 2.402 | 0.1248 | DL | 0.1152 |
| | | PPR VL (Ipsi) | 142.466 ± 7.039% | 10 cells/ 9 mice | Interaction, F (3, 87) = 5.356 | 0.002 | VL | 0.0362 |
| | | PPR DM (Contra) | 97.972 ± 9.605% | 12 cells/ 12 mice | | | DM | 0.0362 |
| | | PPR DM (Ipsi) | 135.955 ± 10.857% | 12 cells/ 12 mice | | | VM | 0.6452 |
| | | PPR VM (Contra) | 117.347 ± 8.066% | 18 cells/ 16 mice | | | | |
| | | PPR VM (Ipsi) | 111.744 ± 5.741% | 18 cells/ 16 mice | | | | |
| | A | Schematic illustration | N/A | N/A | N/A | N/A | N/A | N/A |
| | B | Representative immunostaining images | N/A | N/A | N/A | N/A | N/A | N/A |
| | | D2R+ area (DL,VM GPe) | 122.0 ± 6.91 μm2 | 120 images/ 11 mice | t(238) = 6.651 (Unpaired t-test) | < 0.0001 | N/A | N/A |
| | | D2R+ area (VL,DM GPe) | 67.76 ± 4.38 μm2 | 120 images/ 11 mice | | | | |

| | | | | | | | | |
|---|---|---|---|---|---|---|---|---|
| 7 | C | D2R+RFP+ area (DL,VM GPe) | 16.23 ± 1.11 µm2 | 120 images/ 11 mice | t(238) = 5.581 (Unpaired t-test) | < 0.0001 | N/A | N/A |
| | | D2R+RFP+ area (VL,DM GPe) | 8.83 ± 0.73 µm2 | 120 images/ 11 mice | | | | |
| | | D2R+GFP+ area (DL,VM GPe) | 19.92 ± 2.11 µm2 | 120 images/ 11 mice | t(238) = 3.075 (Unpaired t-test) | 0.0023 | N/A | N/A |
| | | D2R+GFP+ area (VL,DM GPe) | 12.29 ± 1.30 µm2 | 120 images/ 11 mice | | | | |
| | | D2R+GFP+ or D2R+RFP+ area (DL,VM GPe) | 27.79 ± 2.34 µm2 | 120 images/ 11 mice | t(238) = 4.137 (Unpaired t-test) | < 0.0001 | N/A | N/A |
| | | D2R+GFP+ or D2R+RFP+ area (VL,DM GPe) | 15.97 ± 1.65 µm2 | 120 images/ 11 mice | | | | |
| | D | D2R+ area (DL,VM GPe) | 104.0 ± 9.10 µm2 | 120 images/ 11 mice | t(238) = 0.5151 (Unpaired t-test) | 0.6068 | N/A | N/A |
| | | D2R+ area (VL,DM GPe) | 97.36 ± 9.25 µm2 | 120 images/ 11 mice | | | | |
| | | D2R+RFP+ area (DL,VM GPe) | 21.56 ± 2.44 µm2 | 120 images/ 11 mice | t(238) = 1.825 (Unpaired t-test) | 0.0687 | N/A | N/A |
| | | D2R+RFP+ area (VL,DM GPe) | 15.94 ± 1.85 µm2 | 120 images/ 11 mice | | | | |
| | | D2R+GFP+ area (DL,VM GPe) | 6.18 ± 0.83 µm2 | 120 images/ 11 mice | t(238) = 1.082 (Unpaired t-test) | 0.2801 | N/A | N/A |
| | | D2R+GFP+ area (VL,DM GPe) | 5.09 ± 0.55 µm2 | 120 images/ 11 mice | | | | |
| | | D2R+GFP+ or D2R+RFP+ area (DL,VM GPe) | 25.74 ± 2.44 µm2 | 120 images/ 11 mice | t(238) = 2.230 (Unpaired t-test) | 0.0263 | N/A | N/A |
| | | D2R+GFP+ or D2R+RFP+ area (VL,DM GPe) | 18.91 ± 1.83 µm2 | 120 images/ 11 mice | | | | |
| | E | Edge-corrected Ripley's H function analysis | N/A | N/A | N/A | N/A | N/A | N/A |
| | F | Edge-corrected Ripley's H function analysis | N/A | N/A | N/A | N/A | N/A | N/A |
| | G | D2R+ area (Contra DL,VM GPe) | 122.01 ± 6.91 µm2 | 120 images/ 11 mice | Ordinary two-way ANOVA | | Post hoc Holm-Sidak's test | |
| | | D2R+ area (Ipsi DL,VM GPe) | 107.96 ± 9.16 µm2 | 120 images/ 11 mice | GPe subregoin effect, F (1, 476) = 12.61 | 0.0004 | Control - 6-OHDA | |
| | | D2R+ area (Contra VL,DM GPe) | 67.76 ± 4.38 µm2 | 120 images/ 11 mice | 6-OHDA effect,F (1, 476) = 0.7258 | 0.3946 | DL,VM GPe | 0.2754 |
| | | D2R+ area (Ipsi VL,DM GPe) | 97.36 ± 9.25 µm2 | 120 images/ 11 mice | Interaction,F (1, 476) = 5.714 | 0.0171 | VL,DM GPe | 0.0448 |
| | | D2R+RFP+ area (Contra DL,VM GPe) | 16.23 ± 1.11 µm2 | 120 images/ 11 mice | Ordinary two-way ANOVA | | Post hoc Holm-Sidak's test | |
| | | D2R+RFP+ area (Ipsi DL,VM GPe) | 21.56 ± 2.44 µm2 | 120 images/ 11 mice | GPe subregoin effect, F (1, 476) = 9.586 | 0.002 | Control - 6-OHDA | |
| | | D2R+RFP+ area (Contra VL,DM GPe) | 8.83 ± 0.73 µm2 | 120 images/ 11 mice | 6-OHDA effect,F (1, 476) = 8.747 | 0.0032 | DL,VM GPe | 0.0729 |
| | | D2R+RFP+ area (Ipsi VL,DM GPe) | 15.94 ± 1.85 µm2 | 120 images/ 11 mice | Interaction,F (1, 476) = 0.1824 | 0.6695 | VL,DM GPe | 0.0345 |
| | H | Edge-corrected Ripley's H function analysis | N/A | N/A | N/A | N/A | N/A | N/A |
| 8 | A | Schematic illustration | N/A | N/A | N/A | N/A | N/A | N/A |
| | B | Schematic illustration | N/A | N/A | N/A | N/A | N/A | N/A |
| | C | Baseline ($Ca^{2+}$ channel $P_{open}$) | 0.5097 ± 0.0349 | 10 models | Wilcoxon matched-pairs signed rank test | 0.002 | N/A | N/A |
| | | QP ($Ca^{2+}$ channel $P_{open}$) | 0.2664 ± 0.0405 | 10 models | Sum of positive, negative ranks: 0.000 , -55.00 | | | |
| | | Baseline ($Ca^{2+}$ concentration) | 0.6020 ± 0.1056 µM | 10 models | Wilcoxon matched-pairs signed rank test | 0.002 | N/A | N/A |
| | | QP ($Ca^{2+}$ concentration) | 0.3679 ± 0.0908 µM | 10 models | Sum of positive, negative ranks: 0.000 , -55.00 | | | |
| | D | Baseline ($Ca^{2+}$ channel $P_{open}$) | 0.5167 ± 0.0479 | 10 models | Wilcoxon matched-pairs signed rank test | 0.4316 | N/A | N/A |
| | | QP ($Ca^{2+}$ channel $P_{open}$) | 0.5203 ± 0.0486 | 10 models | Sum of positive, negative ranks: 36.00 , -19.00 | | | |
| | | Baseline ($Ca^{2+}$ concentration) | 0.4880 ± 0.1019 µM | 10 models | Wilcoxon matched-pairs signed rank test | 0.3223 | N/A | N/A |
| | | QP ($Ca^{2+}$ concentration) | 0.4960 ± 0.1090 µM | 10 models | Sum of positive, negative ranks: 38.00 , -17.00 | | | |
| | E | Baseline ($Ca^{2+}$ channel $P_{open}$) | 0.5549 ± 0.0434 | 10 models | Wilcoxon matched-pairs signed rank test | 0.2324 | N/A | N/A |
| | | QP ($Ca^{2+}$ channel $P_{open}$) | 0.5390 ± 0.0389 | 10 models | Sum of positive, negative ranks: 15.00 , -40.00 | | | |
| | | Baseline ($Ca^{2+}$ concentration) | 0.8195 ± 0.0764 µM | 10 models | Wilcoxon matched-pairs signed rank test | 0.2227 | N/A | N/A |
| | | QP ($Ca^{2+}$ concentration) | 0.7952 ± 0.0680 µM | 10 models | Sum of positive, negative ranks: 15.00 , -40.00 | | | |
| | F | Baseline ($Ca^{2+}$ channel $P_{open}$) | 0.4178 ± 0.0614 | 10 models | Wilcoxon matched-pairs signed rank test | 0.002 | N/A | N/A |
| | | QP ($Ca^{2+}$ channel $P_{open}$) | 0.2778 ± 0.0421 | 10 models | Sum of positive, negative ranks: 0.000 , -55.00 | | | |
| | | Baseline ($Ca^{2+}$ concentration) | 0.5936 ± 0.0883 µM | 10 models | Wilcoxon matched-pairs signed rank test | 0.002 | N/A | N/A |
| | | QP ($Ca^{2+}$ concentration) | 0.4043 ± 0.0636 µM | 10 models | Sum of positive, negative ranks: 0.000 , -55.00 | | | |
| | G | Baseline ($Ca^{2+}$ channel $P_{open}$) | 0.4732 ± 0.0353 | 9 models | Wilcoxon matched-pairs signed rank test | 0.1641 | N/A | N/A |
| | | QP ($Ca^{2+}$ channel $P_{open}$) | 0.4407 ± 0.0354 | 9 models | Sum of positive, negative ranks: 10.00 , -35.00 | | | |
| | | Baseline ($Ca^{2+}$ concentration) | 0.5746 ± 0.0812 µM | 9 models | Wilcoxon matched-pairs signed rank test | 0.1523 | N/A | N/A |
| | | QP ($Ca^{2+}$ concentration) | 0.5404 ± 0.0736 µM | 9 models | Sum of positive, negative ranks: 10.00 , -35.00 | | | |
| | H | Baseline ($Ca^{2+}$ channel $P_{open}$) | 0.5309 ± 0.0280 | 9 models | Wilcoxon matched-pairs signed rank test | 0.0078 | N/A | N/A |
| | | QP ($Ca^{2+}$ channel $P_{open}$) | 0.4528 ± 0.0358 | 9 models | Sum of positive, negative ranks: 1.000 , -44.00 | | | |
| | | Baseline ($Ca^{2+}$ concentration) | 0.6650 ± 0.0935 µM | 9 models | Wilcoxon matched-pairs signed rank test | 0.0078 | N/A | N/A |
| | | QP ($Ca^{2+}$ concentration) | 0.5680 ± 0.0802 µM | 9 models | Sum of positive, negative ranks: 1.000 , -44.00 | | | |
| | I | Baseline ($Ca^{2+}$ channel $P_{open}$) | 0.4341 ± 0.0639 | 9 models | Wilcoxon matched-pairs signed rank test | 0.0039 | N/A | N/A |
| | | QP ($Ca^{2+}$ channel $P_{open}$) | 0.3318 ± 0.0608 | 9 models | Sum of positive, negative ranks: 0.000 , -45.00 | | | |
| | | Baseline ($Ca^{2+}$ concentration) | 0.4544 ± 0.0682 µM | 9 models | Wilcoxon matched-pairs signed rank test | 0.0039 | N/A | N/A |
| | | QP ($Ca^{2+}$ concentration) | 0.3648 ± 0.0651 µM | 9 models | Sum of positive, negative ranks: 0.000 , -45.00 | | | |
| | J | Baseline ($Ca^{2+}$ channel $P_{open}$) | 0.5001 ± 0.0314 | 17 models | Wilcoxon matched-pairs signed rank test | < 0.0001 | N/A | N/A |
| | | QP ($Ca^{2+}$ channel $P_{open}$) | 0.3607 ± 0.0370 | 17 models | Sum of positive, negative ranks: 1.000 , -152.0 | | | |
| | | Baseline ($Ca^{2+}$ concentration) | 0.6953 ± 0.0625 µM | 17 models | Wilcoxon matched-pairs signed rank test | < 0.0001 | N/A | N/A |
| | | QP ($Ca^{2+}$ concentration) | 0.5048 ± 0.0506 µM | 17 models | Sum of positive, negative ranks: 1.000 , -152.0 | | | |
